# Origins and metabolic evolution of multifunctionality in *Metarhizium robertsii*: linking phenotypic diversity to ecological niche plasticity

**DOI:** 10.64898/2026.01.15.699721

**Authors:** Huiyu Sheng, Raymond J St. Leger

## Abstract

This study investigates the evolution of fungi with complex multifunctional ecological roles using early and recently diverged lineages of *Metarhizium robertsii* as a model. The early diverged strains are characterized by slower insect killing, extensive pre-mortem fungal proliferation within hosts, and prolific sporulation, whereas recently diverged strains show rapid growth, a “kill and consume” strategy linked to destruxin toxin production, and enhanced plant root colonization. Metabolic assays demonstrated that recently diverged strains utilize a broader range of carbon sources, supporting their ecological versatility as pathogens, endophytes, and saprophytes. Germination rates on insect cuticles and plant roots strongly correlate with each other, and with virulence to *Drosophila* and beetles (*Tenebrio molitor*, *Popillia japonica*), highlighting nutritional flexibility as a key driver of ecological adaptation. Immune response assays in *Drosophila* revealed that virulence differences among strains are also partly mediated by host immune activation. These findings suggest that nutritional mode shifts underpin the evolutionary trajectory from specialist insect pathogen to plant associations and enhanced insect virulence within *M. robertsii,* providing a valuable model for studying fungal ecological plasticity and informing the development of fungal biofertilizers and biopesticides.

## Introduction

Fungi perform diverse and essential ecological roles, including decomposing organic matter, forming symbiotic relationships with plants, acting as pathogens, and driving nutrient cycling. These functions are vital for maintaining healthy ecosystems across various habitats. Fungi also have significant commercial value; for example, entomopathogenic fungi are widely used to control agricultural pests and disease vectors (1–4). Some entomopathogenic fungi also form facultative associations with plants as endophytes. These fungi transfer nutrients from soil and insect cadavers to plants, enhance plant stress tolerance, and activate plant defenses against pathogens (5)(6–8). *Metarhizium* species are particularly notable as effective plant-colonizing insect pathogens, making them keystone species in natural ecosystems(9) that contribute to sustainability goals by promoting soil health, biodiversity, and plant growth(4,10–13). This symbiosis benefits both partners as the plants provide shelter, nutrients and access to insect hosts(14), contributing to *Metarhizium*’s high soil populations, reaching up to 10^^6^ colony-forming units per gram of soil (15).

*Metarhizium* species thus exemplify ecologically versatile “multifunctional” fungi that challenge traditional fungal categorizations into distinct ecological niches as pathogen, saprophyte, or endophyte (16,17). Selosse et al (17) suggest that the multifunctional roles exhibited by *Metarhizium* are linked to the frequent evolutionary transitions between animal and plant hosts within the Clavicipitaceae family. Consistent with ecological flexibility, the extent of *Metarhizium*’s interactions varies widely: some species have narrow insect host ranges, while others infect many insect species (18,19). Even closely related strains can differ in their phenotypic versatility, affecting their pathogenicity, saprophytic ability, and stress tolerance (20). Phylogenetic analyses confirm specialist and generalist lineages intermingle (21) indicating that transitioning between life strategies is an inherent trait of *Metarhizium*. However, the specific mechanisms driving this ecological flexibility remain largely unknown. Similarly, in other fungi, the ease with which these transitions occur is still debated (22).

The premise behind the current study is that examining phenotypic plasticity among closely related strains will shed light on the origins of multifunctionality. *Metarhizium robertsii* is the most extensively studied *Metarhizium* species at the physiological level and serves as a model for addressing questions of broad biological significance, such as the origins of new diseases and lifestyles (20,23). Certain strains, such as *M. robertsii* 2575 (Mr2575) can colonize the rhizosphere (the soil layer influenced by root secretions) and rhizoplane (root surface) (7,24,25) and have been shown to promote plant growth in laboratory and field studies(7,24,26). However, the physiological and genetic bases for these traits and the extent of intraspecific variation are not well understood(24).

Using phylogenomic methodologies and experimental data on functional specialization, this study clarifies the evolutionary relationships among eight *M. robertsii* strains, revealing phenotypic and ecological diversity that traces the evolution of multifunctionality. The results highlight the importance of nutritional flexibility in enabling opportunistic colonization of both plants and insects, and reveal a correlation rather than trade-off between increased insect virulence and improved plant root colonization. These findings demonstrate that the combined genomic and experimental data from *M. robertsii* strains provides a valuable resource for studying functional evolution within a fungal species, advancing understanding of niche plasticity, co-evolution with hosts, and informing the development of fungal biofertilizers and biopesticides.

## Results

### M. robertsii phylogeny

The evolutionary trajectory of *M. robertsii* was investigated through a phylogenetic analysis using 8653 single-copy orthologous sequences obtained from eight sequenced *M. robertsii* strains (Fig 1). *M. anisopliae* and *M. robertsii* are members of the PARB species complex of closely related *Metarhizium* species and they diverged approximately 7.1 million years ago (MYA)(19). Using *M. anisopliae* ARSEF 549 as an outgroup (19), we calculated *M. robertsii* strain Mr727, isolated from a katydid (Tettigoniidae) cricket, was the most basal lineage, diverging from other strains around 3.66 million years ago (S1 Table). Mr1120, isolated from soil baited with greater wax moths (*Galleria mellonella*), diverged 2.62 MYA. Mr1046 and Mr1878, both isolated from the Japanese beetle (*Popillia japonica*), diverged 3.22 MYA and 1.46 MYA, respectively. Mr2547 isolated from the related European chafer, *Rhizotrogus majalis*, diverged 1.47 MYA and Mr1-16, isolated from rhizosphere soil on Sorghum roots diverged 1.41 MYA. Strains Mr23 and Mr2575, isolated from Elaterid and Curculionid beetles respectively, diverged approximately 0.29 MYA. Based on phylogenetic relationships and divergence times, *M. robertsii* strains were classified into two groups: early diverged strains (≥ 2.62 MYA) comprising Mr727, Mr1046, and Mr1120, and recently diverged strains (≤ 1.47 MYA) comprising Mr2547, Mr1878, Mr1-16, Mr2575, and Mr23 (Fig 1). The early diverged strain exhibited less clustering (Fig 1), reflecting greater time to accumulate genetic differences and evolve independently.

**Fig 1.**
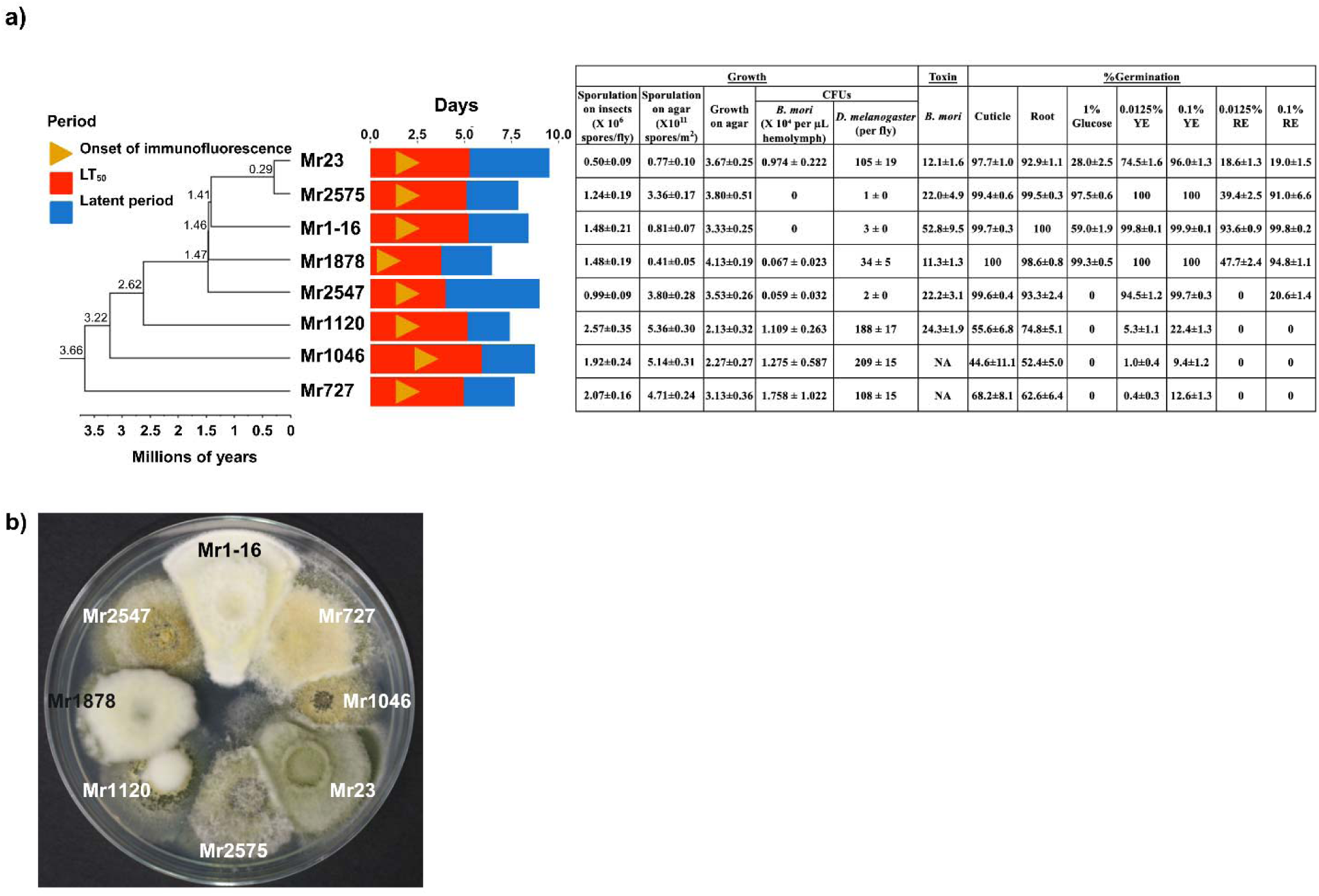
A phylogenomic tree was constructed for eight *M. robertsii* strains. a) The *M. robertsii* strains were separated into early diverged (Mr727, Mr1046, and Mr1120); and recently diverged (Mr1878, Mr2547, Mr1-16, Mr2575, and Mr23) groups. Right of the tree; yellow arrowheads, red bars and blue bars denote the onset of immune response (detected by Drs-GFP fluorescence), virulence (LT_50_) and latent period (the interval between death and sporulation), respectively, as measured in female *Drosophila melanogaster* line Drs-GFP flies. Growth metrics for each strain included sporulation on fly cadavers and agar, growth rate on Potato Dextrose Agar (PDA), fungal proliferation in moribund *Bombyx mori* larvae, and maximum growth in hemolymph (quantified as CFUs) in Drs-GFP females. Toxin production was evaluated by the knockdown (KD) time in *B. mori* using media filtrate. Germination rates were recorded after 12 hours on insect cuticle (fly wings) and plant (*Arabidopsis*) roots, and after 18 hours in media containing glucose (1%), yeast extract (YE) and root exudate (RE) with both YE and RE at 0.0125% or 0.1%. b) The eight *M. robertsii* strains showing phenotypic diversity in growth and sporulation after 14 days growth (27 °C) on PDA.

### *M. robertsii* strains exhibit considerable heterogeneity in virulence profiles and sporulation on cadavers

The *Drosophila* Genetic Reference Panel (DGRP) comprises 192 wild-type *Drosophila melanogaster* lines collected in Raleigh, North Carolina (27). Based on previous work in our laboratory (23), we selected two DGRP fly lines for bioassays: one resistant to *M. anisopliae* ARSEF 549 (Ma549), DGRP RAL 808, and one susceptible, DGRP RAL 321, along with a Drosomycin-GFP (Drs-GFP) reporter line that is intermediate in resistance (23). We verified that RAL 808 is more resistant to Ma549 than RAL 321 and observed similar resistance patterns (RAL 808>Drs-GFP>RAL 321) across all eight *M. robertsii* strains tested (S1 and S2a Figs). Virulence was measured by LT_50_ (time to 50% mortality), showing varying lethality of *M. robertsii* strains against RAL 321 and RAL 808 flies, with notable sex differences, especially in the resistant RAL 808 flies. In RAL 321 females, LT_50_ values ranged from 2.52 (Mr1878) to 3.24 days (Mr1046); and in males, from 2.99 (Mr1878) to 3.67 days (Mr1046). For RAL 808 females, LT_50_ values ranged from 4.36 (Mr1878) to 6.98 days (Mr1046), and in males from 5.24 days (Mr1878) to 8.94 days (Mr1046). In Drs-GFP females LT_50_ values ranged from 3.77 (Mr1878) to 5.93 days (Mr1046); and in males, from 3.71 (Mr1878) to 5.56 days (Mr1046). Thus, males in both DGRP lines were more resistant than females, while in the Drs-GFP line females were more resistant than males (S2a Fig). Nonetheless, strong correlations between males and females across all three fly lines (S2 Table) suggest shared genetic factors influencing resistance.

Resistance profiles across the three *Drosophila* lines to the eight *M. robertsii* strains showed a slightly stronger correlation among males (average: r = 0.77, p = 0.043) than among females (average: r = 0.68, p = 0.066), indicating a more consistent pattern of resistance across male individuals. Plotting LT_50_ values by *M. robertsii* strain and fly line revealed that the less virulent strains had steeper slopes (Fig 2) suggesting that host resistance plays a larger role in determining infection outcomes for these strains. In contrast, highly virulent strains quickly overwhelm host defenses regardless of resistance variation in the three fly lines. Supporting this observation, the slope of LT_50_ across fly lines was positively correlated with average *Drosophila* LT_50_ (males, r = 0.88, p = 0.0041; females, r = 0.83, p = 0.011).

**Fig 2.**
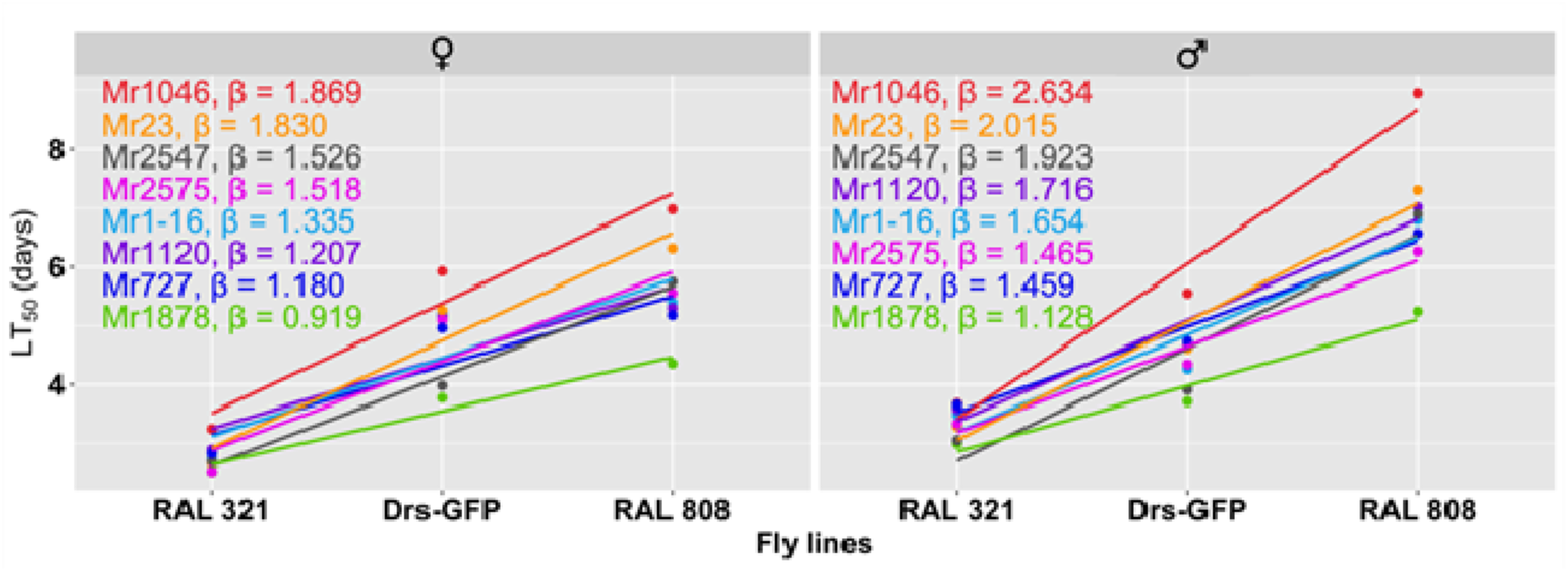
Virulence trends of eight *M. robertsii* strains across three *D. melanogaster* lines (RAL 321, Drs-GFP, and RAL 808). Fly lines were listed from left to right in order of increasing resistance to *M. robertsii*. The slope (β) of the trend line for each *M. robertsii* strain was ranked from the steepest to the shallowest. Less virulent strains showed steeper slopes, suggesting that host genotype has a greater influence on infection outcomes when the pathogen is less virulent. LT_50_ values were calculated based on five independent experiments for each pathogen with at least 30 flies per genotype per experiment.

A virulence bioassay using *Tenebrio molitor* beetles and *M. robertsii* strains revealed a strong correlation with mean LT_50_ values across the three *Drosophila* lines (males, r = 0.84, p = 0.0086), similar to the cross-resistance observed between *Drosophila* lines (males, r = 0.77, p = 0.043), indicating that differences in insect species and body size may not account for virulence differences between *M. robertsii* strains.

Since Mr1046 and Mr1878 were originally isolated from *Popillia japonica* beetles, we also tested virulence against this host. We expected that if a pathogen is highly virulent in one beetle species, it might also be highly virulent in another. Contrary to expectations, virulence to *P. japonica* was less strongly correlated with virulence to *T. molitor* [S2 Table, (r = 0.50, p = 0.21), than with virulence to *Drosophila* [mean male LT_50_ values (r = 0.76, p = 0.028)]. However, Mr1046, the least virulent strain against *T. molitor* and *D. melanogaster* (Fig 1) ranked seventh out of eight in virulence to *P. japonica* (Table 1). Conversely, Mr1878 was among the two most virulent strains against *D. melanogaster*, *T. molitor*, and *P. japonica.* The Lethal Dose 50 (LD_50_’s) values for Mr1878 and Mr1046 against *P. japonica* were 2.41 x 10^6^ and 1.05 x 10^7^ spores/mL, respectively, indicating that Mr1046’s low virulence was not offset by a lower lethal spore concentration (S4 Fig). Overall, recently diverged strains (Mr2547, Mr1878, Mr1-16, Mr2575, and Mr23) killed different insects approximately 1.15-fold faster than early diverged strains (Mr727, Mr1046, and Mr1120, S3 Table).

**Table 1.** Rankings of *M. robertsii* strains based on their insect virulence, proliferation within insect hosts, growth and sporulation on agar, toxin production, and germination in nutrient-rich environments or on fly wings, and interaction with plant roots. YE: yeast extraction; RE: root exudate. A ranking of 1 indicates the most virulent, highest toxin-producing, most CFUs, fastest growth, highest spore counts, fastest germinator. Average virulence ranking was based on the average LT_50_ across all insect bioassays.

| Strain<br>S | Virulence to insects |  |  |  |  |  |  |  |  | Growth |  |  |  | Toxi<br>n | %Germination |  |  |  |  |
| --- | --- | --- | --- | --- | --- | --- | --- | --- | --- | --- | --- | --- | --- | --- | --- | --- | --- | --- | --- |
|  | D. melanogaster |  |  |  |  |  | T.<br>molito<br>r | P.<br>japonic<br>a | Averag<br>e | Growt<br>h on<br>agar | Sporulatio<br>n on agar | CFUs |  |  | Cuticle<br>(Drosophil<br>a wings) | Roo<br>t | 1%<br>Glucos<br>e | 0.0125<br>% YE | 0.0125<br>% RE |
|  | ♂ |  |  | ♀ |  |  |  |  |  |  |  | B.<br>mor<br>i | D.<br>melanogast<br>er |  |  |  |  |  |  |
|  | RA<br>L<br>321 | Drs<br>-<br>GF<br>P | RA<br>L<br>808 | RA<br>L<br>321 | Drs<br>-<br>GF<br>P | RA<br>L<br>808 |  |  |  |  |  |  |  |  |  |  |  |  |  |
| Mr23 | 3 | 5 | 7 | 3 | 7 | 7 | 3 | 5 | 6 | 3 | 7 | 4 | 4 | 2 | 5 | 5 | 4 | 5 | 4 |
| Mr2575 | 4 | 4 | 2 | 2 | 4 | 5 | 5 | 4 | 3 | 2 | 5 | 7 | 8 | 3 | 4 | 2 | 2 | 1 | 3 |
| Mr1-16 | 5 | 3 | 4 | 5 | 6 | 4 | 6 | 6 | 5 | 5 | 6 | 7 | 6 | 6 | 2 | 1 | 3 | 3 | 1 |
| Mr1878 | 1 | 1 | 1 | 1 | 1 | 1 | 2 | 1 | 1 | 1 | 8 | 5 | 5 | 1 | 1 | 3 | 1 | 1 | 2 |
| Mr2547 | 2 | 2 | 5 | 4 | 2 | 6 | 1 | 3 | 2 | 4 | 4 | 6 | 7 | 4 | 3 | 4 | 5 | 4 | 5 |
| Mr1120 | 6 | 6 | 6 | 7 | 5 | 3 | 4 | 8 | 7 | 8 | 1 | 3 | 2 | 5 | 7 | 6 | 5 | 6 | 5 |
| Mr1046 | 8 | 8 | 8 | 8 | 8 | 8 | 8 | 7 | 8 | 7 | 2 | 2 | 1 | 7 | 8 | 8 | 5 | 7 | 5 |
| Mr727 | 7 | 7 | 3 | 6 | 3 | 2 | 7 | 2 | 4 | 6 | 3 | 1 | 3 | 7 | 6 | 7 | 5 | 8 | 5 |

Differences in reproductive potential between *M. robertsii* strains, measured by sporulation capacity (spore counts per cadaver), were similar across *D. melanogaster*, *T. molitor*, and *P. japonica* (S2 Table, correlation coefficients between 0.51 and 0.93). This indicates that sporulation on an insect is influenced more by the strain’s intrinsic traits than by the host species. In contrast, there was a strong negative correlation between the latent period (time from host death to initiation of spore production) and final spore counts (S2 Table, r between -0.93 and - 0.64). Overall, early diverged strains killed hosts more slowly but had shorter latent periods post-mortem (p < 0.05) and produced more spores on all three insect species (S3 Table). For example, although Mr1046 took longer to kill than Mr1878 (Welch’s t-test: t = 10.837, p = 0.0076), it sporulated more rapidly after death (t = -54.22, p = 1.241e-12), and produced 68 times more spores (t = 4.59, p = 0.0013) (S5 Fig).

### *M. robertsii* strains show high variation in disease-relevant traits on the cuticle and in the hemocoel

Host mortality results from complex interactions between pathogen traits-such as spore germination capacity and toxin production-and host defenses, including physical barriers like the cuticle and immune responses (20). In the *Metarhizium* infection process, spore germination is a crucial early step that strongly affects infection success (20). Our analysis showed significant differences among the eight *M. robertsii* strains in both germination frequency (One-way ANOVA, p = 9.66e-14) and germling length (One-way ANOVA, p < 2e-16) on fly wings. These two factors were strongly positively correlated (r = 0.82, p = 0.014). The early diverged strains Mr1046, Mr727 and Mr1120 had lower germination rates (Average germination: 56.1%, range: 44.6-68.2%) and produced shorter germlings on fly wings compared to the recently diverged strains (Average germination: 99.3%, range: 97.7-100%) (S6 Fig). Consequently, there was a negative correlation between phylogenetic divergence times and germination rate on cuticle (r = -0.83, p = 0.0112). Furthermore, the LT_50_ values of the eight *M. robertsii* strains against flies, *T. molitor*, and *P. japonica* were significantly negatively correlated with germination ability on fly wings (S4 Table, -0.90 ≤ r ≤ -0.40), indicating that strains with higher germination rates killed hosts faster. We conclude that *M. robertsii* strains differ in their ability to germinate on the fly surface depending on their evolutionary divergence, and that this early germination ability on *Drosophila* cuticle reliably predicts their lethality across multiple insect species.

### Recently diverged strains produce higher levels of destruxins

Mr23 and Mr2575 have been shown to produce destruxins (dtxs), secondary metabolites that cause flaccid paralysis and muscle contraction in insects, and suppress both cellular and humoral immune responses (28) (29). To compare the toxigenicity of *M. robertsii* strains, we used an established protocol of injecting culture filtrates into *Bombyx mori* caterpillars and measuring the time to paralysis(29) (Fig 1). A transgenic Mr2575 strain, dtx-KO Mr2575, that does not produce dtx(30) served as a control source of culture filtrate. Culture filtrates from dtx-KO Mr2575, Mr1046, and Mr727 neither induced paralysis in *B. mori* nor affected righting time (the time for an inverted caterpillar to right itself) within ten minutes post-injection (Welch’s t-test: p > 0.05). In contrast, filtrates from Mr1878, Mr2547, Mr2575, and Mr23 caused permanent paralysis in all tested *B. mori*.

The filtrate from the fast-killing Mr1878 induced the fastest paralysis (knock down time: 11.29 ± 1.33s), which was statistically similar (p>0.05) to Mr23 (12.13 ± 1.58s) but significantly faster (p<0.05) than Mr2547 (22.15 ± 3.10s), Mr2575 (22.03 ± 4.85s), Mr1120 (24.31 ± 1.85s), and Mr1-16 (52.81 ± 9.47s). Some *B. mori* injected with Mr1-16 (5/30) or Mr1120 (2/30) did not become paralyzed. Mr1-16 took significantly longer than Mr1120 to induce paralysis (p < 0.05), and 21/30 *B. mori* had recovered (able to right themselves within 5s) one day after injection.

Although knockdown times showed a positive but statistically non-significant correlation with virulence [average *Drosophila* males (r = 0.30, p = 0.56), *P. japonica* (r = 0.38, p = 0.46), and *T. molitor* (r = 0.67, p = 0.15), further investigation was conducted on toxin production and fungal proliferation within hosts by quantifying colony-forming units (CFUs) in the hemolymph of *B. mori* and female Drs-GFP flies. These showed a strong positive correlation (r = 0.82, p = 0.012), suggesting that each strain possesses traits, such as virulence factors or immune evasion strategies, that allow them to proliferate similarly within the hemocoel of both host species. Alternatively, the immune systems of both insect species may react in a quantitatively similar manner to these pathogens, resulting in similar levels of control (or lack thereof). Non-toxigenic early diverged strains Mr727 and Mr1046 produced approximately three-fold more CFUs than toxigenic strains (*Drosophila* CFU mean ± SE: 159 ± 51 CFU per fly vs. 56 ± 31 CFU per fly; S6 Table). Another early diverged strain, Mr1120, which produced low toxin levels, generated 188 ± 17 CFU, whereas at the other extreme the toxigenic Mr2575 and Mr2547 produced one or two CFU per fly. Interestingly, the recently diverged strain, Mr1-16, which produced low toxin levels, showed minimal CFU proliferation within flies (max CFUs: 3 ± 0).

Due to their large size and soft unpigmented cuticles, caterpillars are more suitable for observational studies than beetles or *Drosophila*. Observations of infected *B. mori* caterpillars’ 12 hours before death revealed *M. robertsii* strains growing both on and beneath cuticle surfaces, with numerous melanized spots indicating penetration sites (Fig 3). All recently diverged strains showed growth on host surfaces, with Mr1-16 showing the most extensive growth (Fig. 3d), similar to its rapid germination and growth on fly wings (S4 Table, S6 Fig), and also destroying large areas of the epidermis beneath penetration sites, as evidenced by green fluorescent fungus associated with the destruction. However, *B. mori* hemolymph contained very few Mr1-16 blastospores (< 3 CFUs in 3uL of hemolymph), similar to results with *Drosophila*, indicating that CFU counts in hemolymph may not accurately reflect biomass for some strains. It should also be noted that CFU counts in *Drosophila* were measured after centrifuging fly homogenates, so fungi attached to the fly cuticle surface or undersurface were not counted. Infections with the high toxin producing Mr1878-GFP, Mr2547-GFP, and Mr2575-mCherry also resulted in low CFU counts.

**Fig 3.**
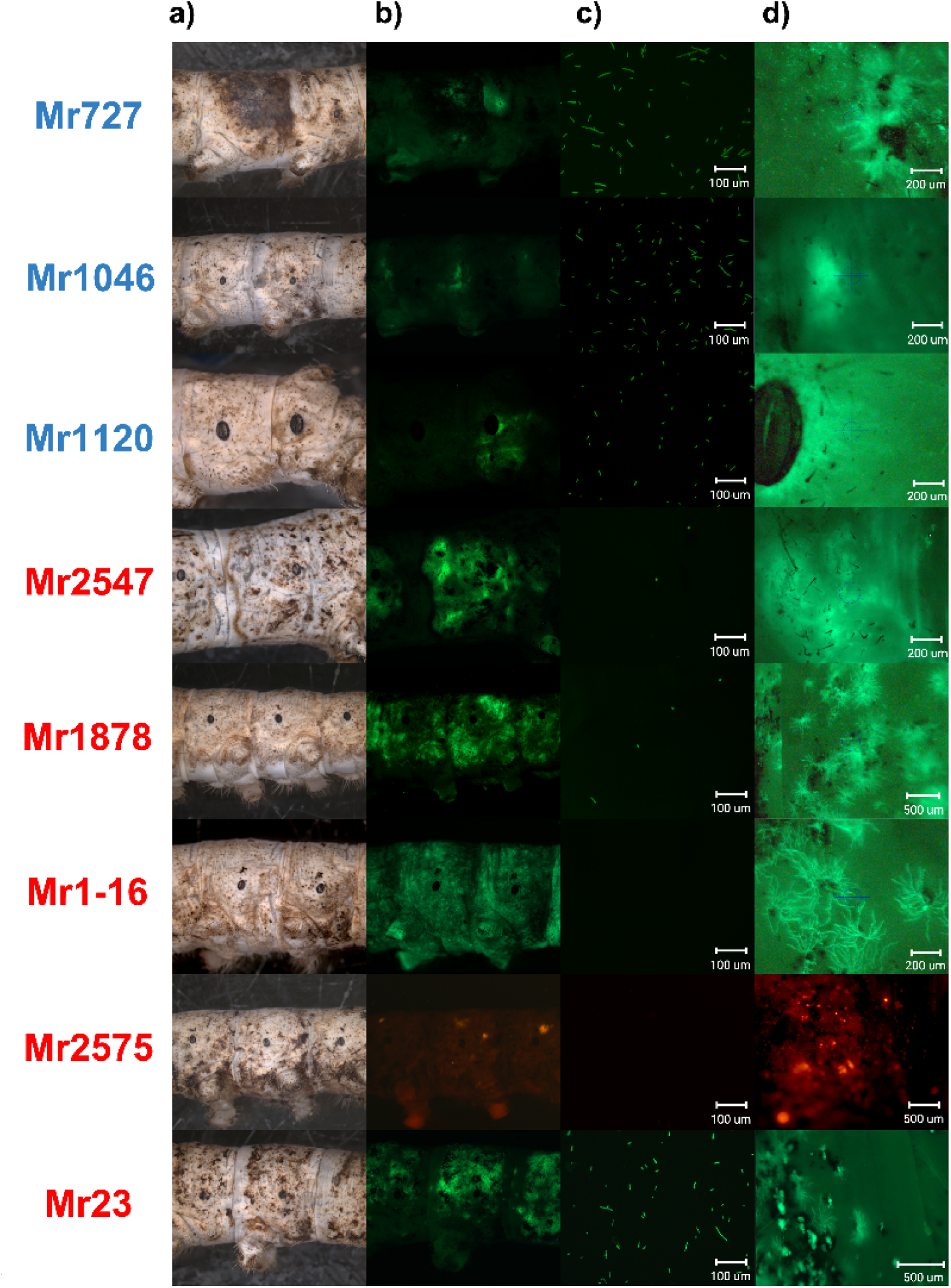
Colonization of *B. mori* by topically applied mCherry- or GFP-tagged *M. robertsii* strains. a) bright field and b) overlay of mCherry or GFP fluorescence of a moribund (unable to right itself within five minutes) *B. mori* caterpillar infected with a *M. robertsii* strain. c) mCherry or GFP fluorescence images of hemolymph extracted from the same *B. mori*. d) growth of mCherry- or GFP-tagged *M. robertsii* on or beneath the cuticle of *B. mori*. Names of early diverged strains are shown in blue and those of recently diverged strains in red. Additional images are shown in S7 Fig.

Early diverged strains (average: 1.381 ± 0.170 X 10^4^ CFUs per uL of hemolymph) showed 6-fold higher CFU counts at the moribund stage compared to most recently diverged strains (average: 0.205 ± 0.0368 X 10^4^ CFUs per uL of hemolymph), except for Mr23 that shows high blastospore counts in both flies and caterpillars (Fig 1).

Slower growing strains on PDA (Mr1046 and Mr1120) displayed little surface growth on *B. mori*, although there were numerous penetration sites shown by melanized spots. In contrast, Mr727 exhibited growth characteristics similar to recently diverged strains, with extensive hyphal development on *B. mori* surfaces and beneath cuticles, although ungerminated conidia clustered on the cuticles suggesting lower germination rates (Fig 3d).

### Recently diverged strains exhibited rapid growth on nutrient media, early germination on cuticle, and low sporulation rates on various substrates

The growth patterns of *M. robertsii* strains were compared across different media: high-nutrient Sabouraud dextrose agar + 1% yeast extraction (SDA+1% YE), medium nutrient PDA, and low-nutrient Czapek-Dox agar (CZA) (S8 Fig). Among these, growth on PDA best simulated the insect environment, as indicated by the most significant negative correlation with LT_50_ values in *Drosophila* (males, r = -0.80, p = 0.018) and other insects (S4 Table). This means strains that grew faster on PDA were usually fast killers. Recently diverged strains grew significantly faster on PDA than early diverged strains (S3 Table, p = 3.01e-6). Rapid growth on PDA was strongly positively correlated with germination on *Drosophila* cuticle (r = 0.92, p = 0.0014). However, because early diverged strains grew slowly but sporulated well, growth on PDA negatively correlated with sporulation on both PDA (r = -0.76, p = 0.02) and insect cadavers (S4 Table).

As previously mentioned, early diverged strains showed high fungal loads within insects so CFU counts negatively correlated with growth on agar (S5 Table, *Drosophila*: r = -0.83, p = 0.01; *B. mori*: r = -0.63, p = 0.094) and germination rate on fly wings (*Drosophila*: r = -0.92, p = 0.0013; *B. mori*: r = -0.78, p = 0.022). Conversely, these CFU counts positively correlated with sporulation on agar (*Drosophila*: r = 0.58, p = 0.13; *B. mori*: r = 0.57, p = 0.14) and in insects (e.g., female RAL 808 flies, *Drosophila*: r = 0.80, p = 0.017; *B. mori*: r = 0.67, p = 0.066).

In summary, early diverged Mr727, Mr1046, and Mr1120 are characterized by poor germination on cuticles, slow growth on agar, extensive proliferation as CFUs inside hosts, and prolific sporulation on insect cadavers and agar. In contrast, recently diverged strains show rapid growth on PDA, early germination and prolific growth on and under cuticles, high virulence, and low sporulation rates on various substrates. Except for Mr1-16, these recently diverged strains produce high levels of destruxins in culture, which may contribute to their lethality despite limited fungal growth inside insect hemolymph.

### *M. robertsii* strains elicit a wide range of responses by the *Drosophila* immune system

We next investigated whether the host immune response might contribute to between *M. robertsii* strain variation in virulence. In *Drosophila*, fungal infections are mainly controlled by the conserved Toll immune pathway. Fungal proteases are detected by the sensor protease Persephone (*Psh*), which activates Toll to produce antimicrobial peptides (AMPs) such as drosomycin (Drs), the principal anti-fungal peptide in *Drosophila* (31).

Using Drosomycin (Drs)-GFP reporter flies, which show GFP fluorescence proportional to drosomycin production and immune activation (32) we observed that the highly virulent strain Mr1878 induced a significant increase in Drs-GFP fluorescence just one day after infection (Welch’s t-test, p ≤ 0.05), demonstrating rapid immune recognition. Most other strains caused significant immune activation by day two, except for Mr1046, which showed a delayed response, becoming significant only by day three (Fig 4). The immune response appears to be linked to fungal penetration of the cuticle, as indicated by correlations between Drs-GFP fluorescence and both cuticle germination (r = -0.64, p = 0.089) and LT_50_ values (r = 0.81, p = 0.014). Overall, these results suggest that *M. robertsii* strains with slower germination, growth and killing rates provoked delayed immune responses compared to virulent strains, and that neither the immune response nor virulence is contingent upon the extensive fungal proliferation in the hemocoel characteristic of early diverged strains (Fig 4, S9 Fig).

**Fig 4.**
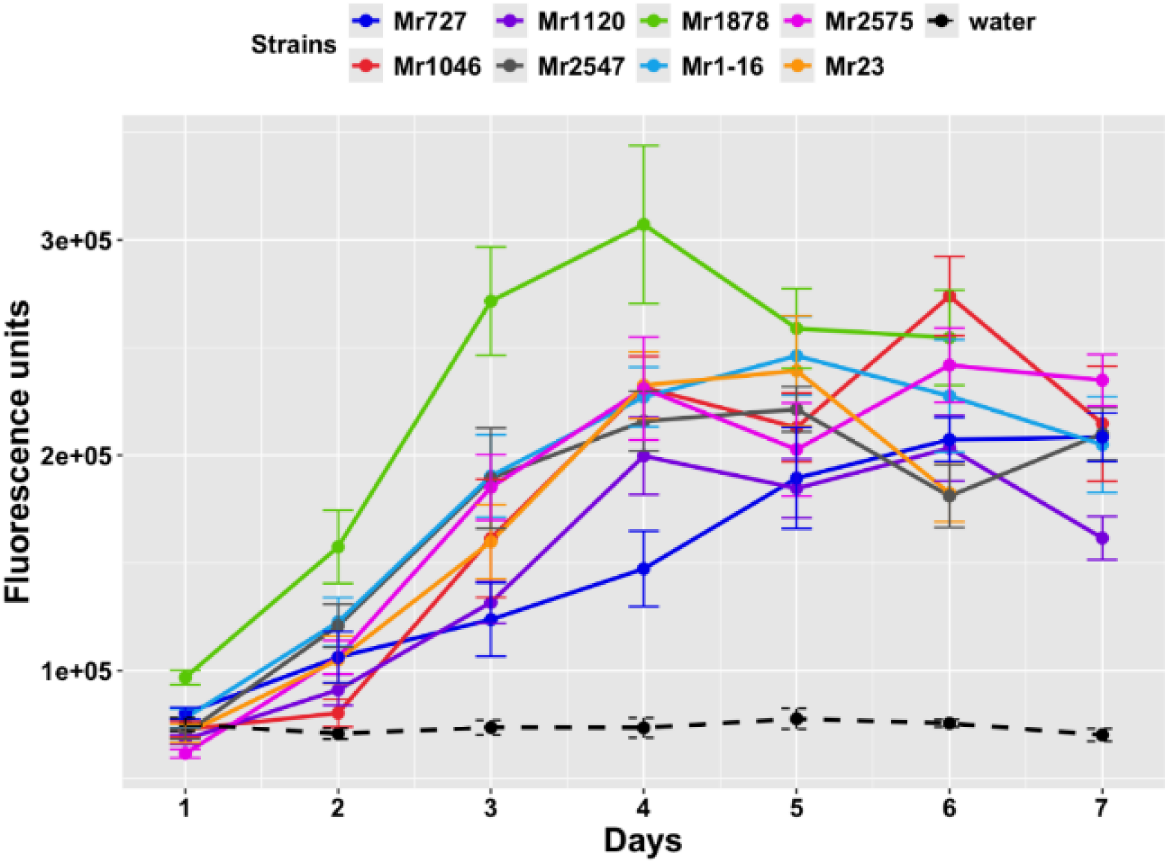
Drs-GFP immunofluorescence response of *D. melanogaster* to infection with different *M. robertsii* strains. Points represent the means of 10 individual flies ± SE. Control flies (dashed line) were treated with water instead of spore suspensions.

### Persephone (Psh)-dependent processes and prophenoloxidases contribute to resistance to *M. robertsii*

Immunocompromised *Drosophila* lacking the pathogen sensor Persephone (Δ*Psh*) died significantly faster when infected with Mr2575(20). In this study, we used Δ*Psh* flies to assess how the insect immune system affects lethality differences among *M. robertsii* strains. Compared to wild-type (*w*^A5001^) flies, Δ*Psh* flies showed significantly reduced survival (Tukey test, p ≤ 0.05) after infection with all eight *M. robertsii* strains (Fig 5 and S10 Fig). The range of LT_50_ values from maximum to minimum among Δ*Psh* flies was considerably narrower—about two-thirds smaller—than in wild-type flies (males: 0.61 days vs. 1.89 days; females: 0.65 days vs. 1.67 days). This suggests that the deletion of *Psh* reduces the differences in virulence observed between *M. robertsii* strains. For example, Mr1046, which killed WT flies comparatively slowly (Tukey’s test, p < 0.05) caused mortality rates similar to other strains in Δ*Psh* flies (Tukey’s test, p > 0.05). These findings indicate that the insect immune system contributes to the variation in survival times following infection by different *M. robertsii* strains. However, if differential sensitivity to Toll pathway products fully explained these differences, then LT_50_ values in Δ*Psh* flies would be consistent across strains. Instead, Mr1878 remained the most pathogenic strain against both WT and Δ*Psh* flies. Compared to WT flies, Δ*Psh* flies succumbed 19% (females) and 13.8% (males) faster when infected with Mr1878, and ∼36% faster in both sexes when infected with Mr1046, consistent with Mr1046 being particularly susceptible to host immunity.

**Fig 5.**
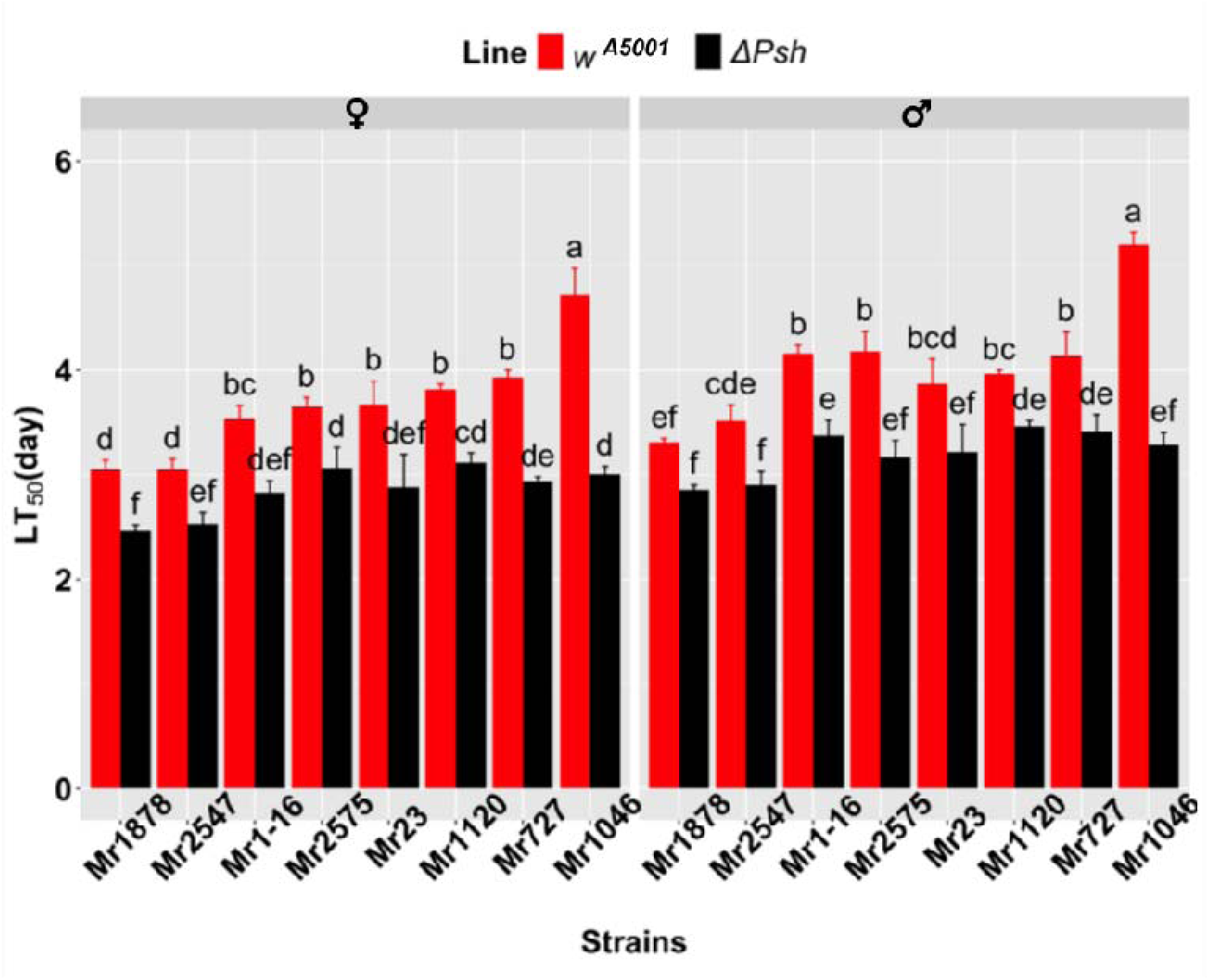
Survival (measured as LT_50_ values post infection with *M. robertsii* strains) of a *Drosophila* fly line disrupted in *Psh* and its isogenic WT background. The different lowercase letters at the top of each bar indicate significant differences by Tukey’s test (Two-way ANOVA, p<0.05). a) LT_50_ values (mean ± SD) of Δ*Psh* and its corresponding WT (*w*^A5001^), each treatment was repeated three times with ∼30 flies per replicate.

In both wild-type (WT) and Δ*Psh* lines, females were more susceptible than males to all *M. robertsii* strains (Log-rank test, p < 0.05). However, the differences in LT_50_ between sexes in Δ*Psh* flies did not correlate with those in WT flies (r = -0.1, p = 0.82). The absence of *Psh* increased the sex-based susceptibility differences for strains Mr1878, Mr1120, Mr23, and Mr727, but reduced these differences for Mr1046, Mr2575, Mr2547, and Mr1-16. This indicates that *Psh* influences the degree of sexual dimorphism in susceptibility to *M. robertsii* infection but does not determine which sex is more vulnerable.

Prophenoloxidases (PPOs), crucial for melanization at infection sites and hemolymph, represent an early host response to *M. robertsii* infection (33,34). The *Drosophila* genome contains three PPO-coding genes (33). To evaluate their role in the immune response against *M. robertsii*, we compared the effects of strains Mr2547 and the slower killing Mr1046 (Fig 6). Both strains killed Δ *PPO^1,2,3^*flies approximately 50% faster than WT flies (*w^1118^*). In contrast, mortality rates in Δ*Psh* flies increased by only 36% (Mr1046) and 17% (Mr2547), indicating that PPOs have a greater impact on survival than *Psh.* Mr2547 killed both WT and Δ*PPO_1,2,3_* flies significantly faster than Mr1046 (Welch’s t-test, p < 0.05), demonstrating that differences in susceptibility to melanization do not explain the strain-specific differences in virulence. Additionally, female WT and Δ*PPO^1,2,3^* flies were significantly more susceptible than males to both Mr1046 and Mr2547 infections (Fig 6, Log-rank test, p < 0.05), suggesting that melanization, like Psh, does not account for the observed sexual dimorphism in susceptibility.

**Fig 6.**
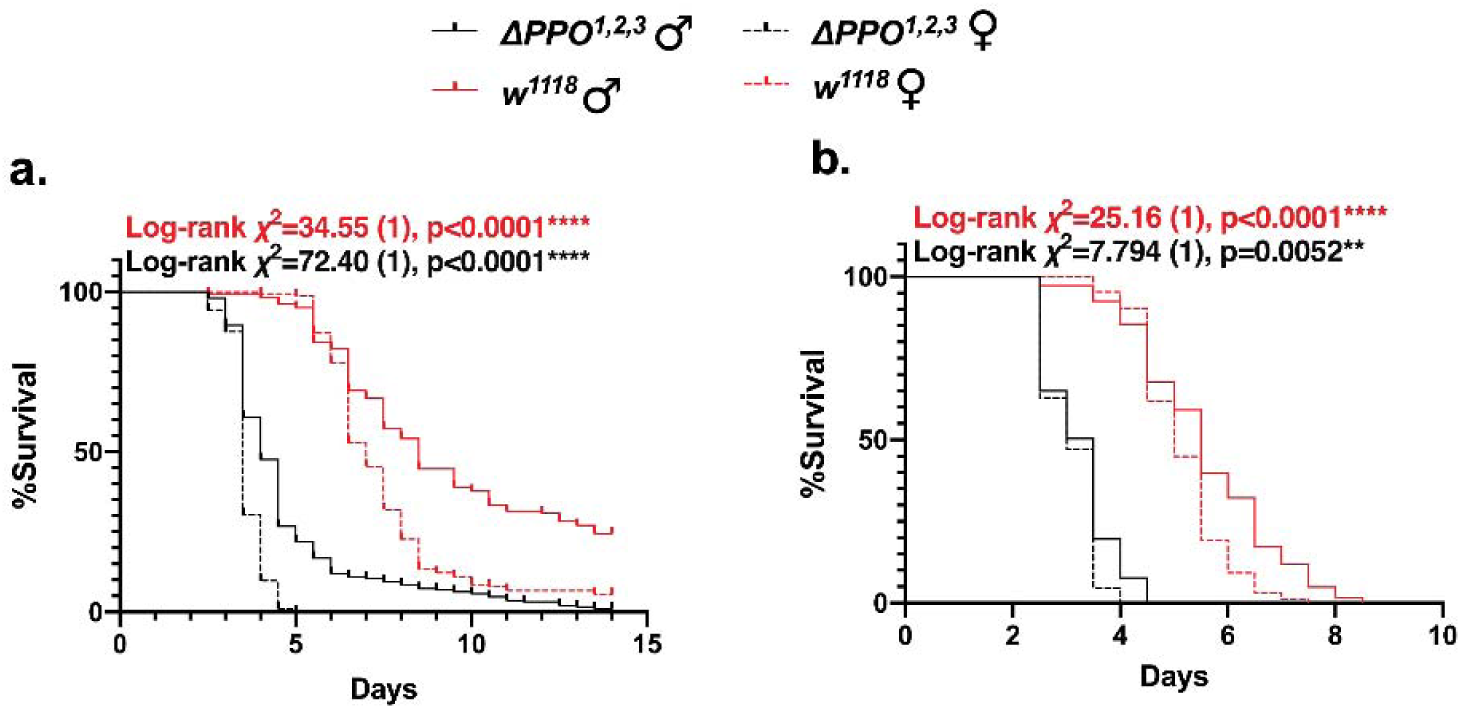
Quantifying the role that melanization plays in resisting infection. a) Survival curves (Kaplan-Meier) of Δ*PPO^1,2,3^*and its isogenic background *w^1118^* against Mr1046 (a) and Mr2547 (b), Each treatment was repeated five times with ∼30 flies per replicate. The significance of differences between males and females was evaluated using log-rank tests with Δ*PPO^1,2,3^* indicated in black and *W^1118^*in red.

### Variation in endophytic ability across strains correlates with their virulence to insects

*M. robertsii* spores must attach to and germinate on plant roots to establish an association. Our analysis found a strong positive correlation between spore attachment and germination on roots (Spearman’s rank correlation: r = 0.88, p = 0.0072). However, this variation was not continuous (Fig 7a), allowing us to classify strains into three groups using Mr2575 as a standard: (1) Mr1046, Mr727, and Mr1120 showed lower attachment and germination than Mr2575; (2) Mr1878, Mr2547, Mr2575, and Mr23 had low to intermediate attachment but high germination rates, (3) Mr1-16 exhibited higher attachment and germination than Mr2575. These patterns aligned with the phylogeny of *M. robertsii*: earlier diverged strains had lower affinity to roots, while recently diverged strains showed increased affinity.

**Fig 7.**
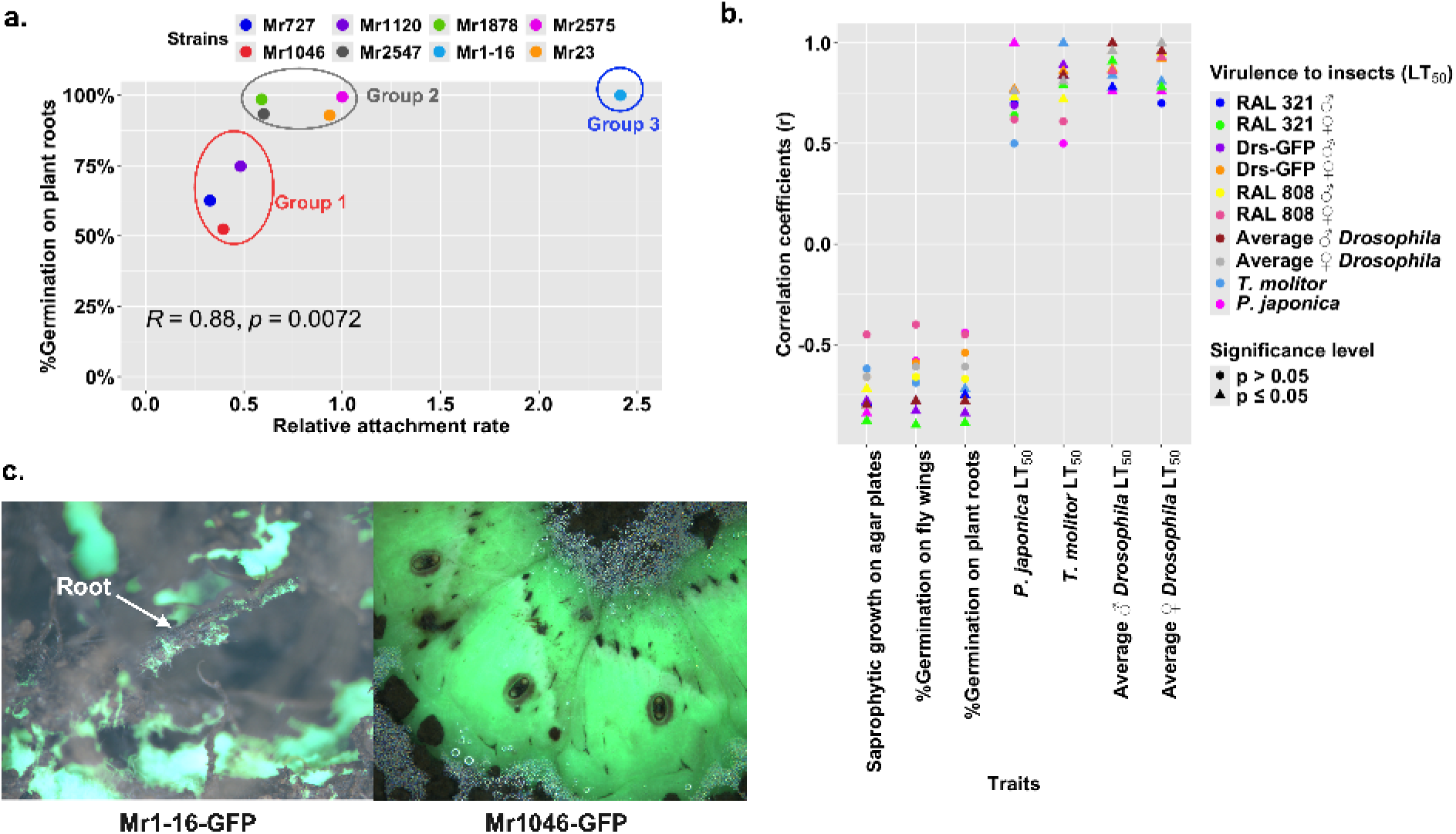
The relationship between *M. robertsii*’s endophytic ability on plant roots and virulence to insects. a) Correlation between attachment and germination rate on *Arabidopsis thaliana* roots, with Mr2575’s rate set as the baseline (1.0) for comparison with the other seven strains. b) Correlation coefficients (r) between the virulence of *M. robertsii* strains to flies, *T. molitor*, and *P. japonica* (LT_50_ values) and their germination ability on plants, fly wings (% germination), or growth on agar. c) Fungal development from *Manduca sexta* cadavers buried in soil. Mr1046-GFP colonized cadavers but produced few emergent hyphae and did not interact with plant material, whereas Mr1-16-GFP produced emergent hyphae and colonized neighboring plant roots (colonized root indicated by arrow). GFP-strains were visualized with epifluorescence, with filters set to detect GFP.

We analyzed the relationships among insect virulence, saprophytic growth, and root attachment/germination. Germination on roots correlated strongly with germination (r = 0.94, p ≤ 0.001) and growth (r = 0.84, p ≤ 0.01) on fly wings, and insect virulence (LT_50_ values) was negatively correlated with root germination (male *Drosophila* r = -0.75, p = 0.033, females r = -0.89, p = 0.0034), indicating that slow killing strains also germinated less on roots (Fig 7b and S4 Table). Thus, insect virulence and endophytic ability are linked through germination traits, with highly insect virulent *M. robertsii* strains, such as Mr1878 and Mr2547 germinating efficiently on plant roots.

To explore lifestyle differences between early and recently diverged *M. robertsii* strains, we buried infected *Manduca sexta* cadavers in soil and monitored fungal development (Fig 7c). Using GFP tagged strains, we observed that Mr1046 extensively colonized cadavers, but the sparse hyphae that emerged from these did not interact with surrounding plant material. In contrast, the strong endophyte Mr1-16 produced hyphae that emerged from cadavers and grew along plant roots to colonize the soil environment (Fig 7c). Cadaver decomposition varied, with Mr2575-infected cadavers showing the most disintegration. This was supported by nematode counts on cadaver surfaces (S7 Table): Mr2575 cadavers had significantly more nematodes (29 ± 3) than those infected by Mr1046 (15 ± 3) (Welch’s t-test, t = 2.56, p = 0.0082), indicating greater breakdown and ecological activity.

### Variation in metabolic profiles across strains corresponds to variation in virulence and endophytism

In previous work, we evaluated the metabolic capabilities of *Metarhizium* species by examining their germination in yeast extract (YE) and glucose solutions (35). In this study, we compared the germination of *M. robertsii* strains on plant roots and fly wings with their germination in YE, glucose, and root exudate (RE) solutions (S11 Fig).

All eight strains germinated in 0.1% YE within 18 hours post-inoculation. However, the early diverged strains Mr1046 and Mr727 did not germinate in 0.0125% YE before 18 hours. The recently diverged strains germinated within 12 hours in 0.1% RE, and except for Mr2547, they also germinated within 18 hours in 1% glucose and within 12 hours in 0.0125% RE. Early diverged strains failed to germinate on 1% glucose and RE at 18 hours. Recently diverged strains not only germinated faster but also showed significantly faster growth (S8 Table).

Germination in RE and YE showed positive correlations (S9-10 Tables, r ≥ 0.73, p < 0.05) with attachment and germination on plant roots (S12 Fig). Furthermore, both germination and germling length on insect cuticles strongly correlated (r ≥ 0.76, p < 0.05) with germination in RE and YE, indicating that robust germination on insect cuticles and plant roots is linked to the ability to germinate and grow efficiently on diverse nutrients. Additionally, germination capacity in RE, YE, and 1% glucose negatively correlated with LT_50_ values (S12 Fig, LT_50_ values versus 0.1% YE: r = -0.87, 0.0125% YE: r = -0.85, 0.1% RE: r = -0.85, 0.0125% RE: r = -0.62, 1% glucose: r = -0.65), meaning strains such as Mr2575, Mr1878, and Mr1-16 that germinate rapidly in RE or YE also germinate quickly on insect cuticles and kill insects faster.

For a phylogenetic comparison of *M. robertsii*’s metabolic capacities, we used Filamentous Fungi plates (FF plate) from Biolog to assess utilization of 95 carbon sources grouped into seven categories: amines and amides, amino acids, carbohydrates, carboxylic acids, water, miscellaneous, and polymers. We collected complete utilization and growth curves for each substrate due to their differing kinetics (S13-28 Figs). Cluster analysis separated the 8 strains into recently diverged and early diverged clusters, with recently diverged strains showing faster growth rates (S29 Fig). Strains Mr2575, Mr1-16, and Mr23 developed a purplish color in the water control well (A1) as early as day 1 (S29c Fig), indicating mitochondrial activity even without added nutrients. These strains also showed the highest substrate utilization on day 1, especially for carbohydrates and amino acids (S30 Fig). However, turbidity-based mycelial growth estimates were similar across strains, suggesting metabolic activity without proportional biomass accumulation. Consequently, the correlation between substrate utilization and biomass production was weaker in recently diverged strains (S11 Table, average Spearman’s r = 0.80) than in early diverged strains (average r = 0.94) on day 1. By day 5, all strains showed a strong correlation between substrate utilization and growth.

By day 4 (Fig 8), Mr2547 and Mr1878 reached utilization levels similar to those of Mr1-16, Mr2575, and Mr23 on day 3, reflecting a slower but sustained metabolic response. Although Mr1046, Mr727 and Mr1120 improved carbon source utilization by day 4, they lagged behind the recently diverged strains, especially in amino acid and carbohydrate metabolism. On day 5, recently diverged strains still exhibited greater substrate utilization and higher biomass production across all carbon source categories (S12 Table), except for polymers (α-cyclodextrin, β-cyclodextrin, dextrin, and glycogen). Apart from polymer**s,** strains that performed well in one nutrient category (amines and amides, amino acids, carbohydrates, carboxylic acids, and miscellaneous) also performed well in others (S13-14 Tables).

**Fig 8.**
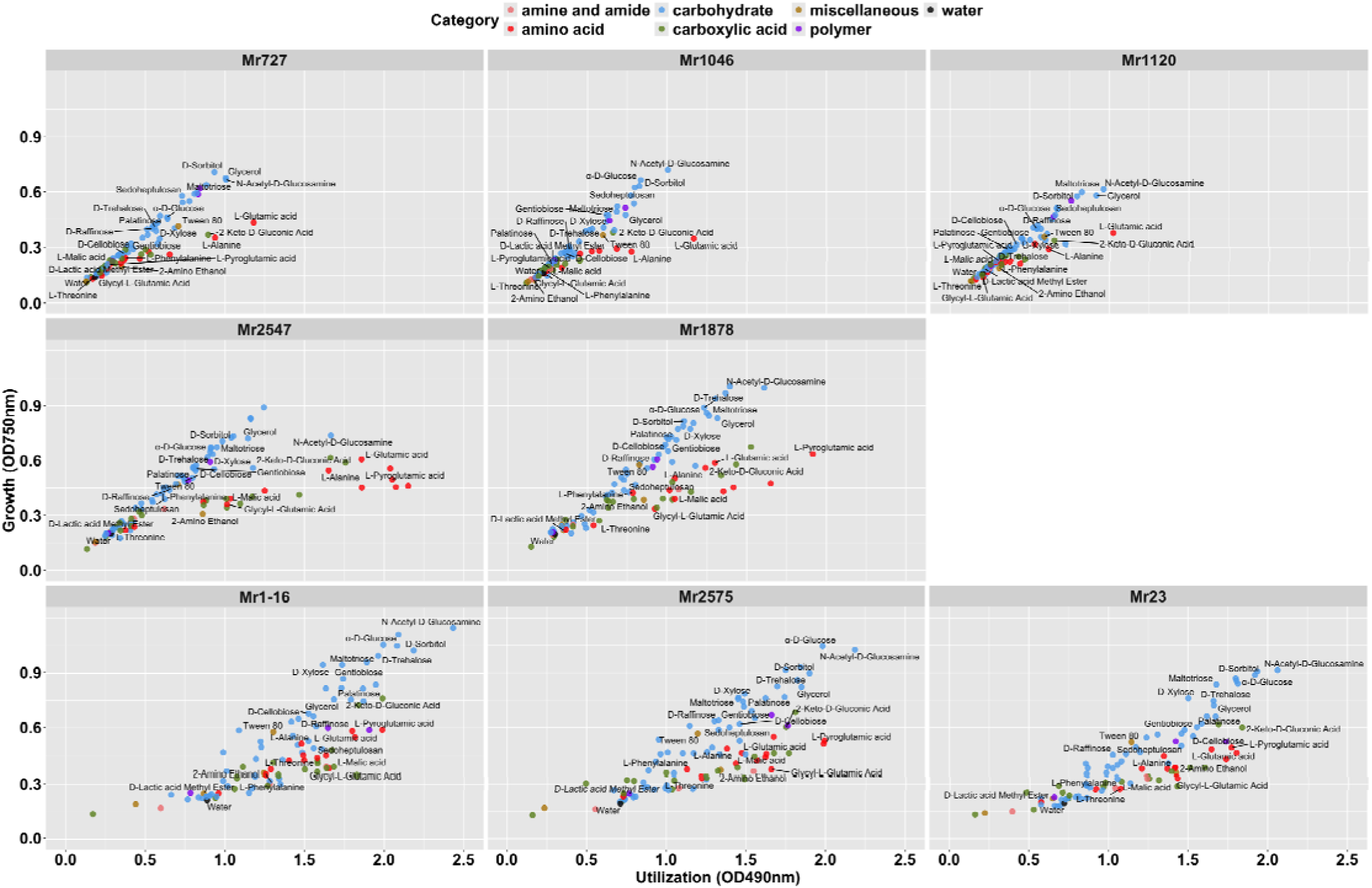
Correlation showing *M. robertsii* FF MicroPlate C-source utilization and growth rates. The utilization rate and growth intensity rate were calculated using absorbance at 490 nm and 750 nm, respectively, on day four. Correlations of C-source utilization and growth rates on day one, two, three, and five were shown in S30-33 figures.

Examining nutrient categories individually, strains Mr-16, Mr2575 and Mr23 utilized carboxylic acids, amino acids and carbohydrates with similar efficiency, but substrates supporting high biomass production were almost exclusively carbohydrates (Fig 8). Mr1-16 was the fastest metabolizer of 69 of the 95 carbon substrates and showed the most growth on 37, with 32 of these (including 21 carbohydrates) among the fastest utilized (linking carbon use to biomass accumulation) (S36 Fig and S15 Table). Mr2575 ranked second, with highest utilization of 12 substrates (including 5 carbohydrates) and highest growth on 25, mostly (14) carbohydrates. Mr23 and Mr1878 showed similar utilization and growth patterns to Mr2575, while Mr2547 had lower carbohydrate utilization but was the fastest utilizer of several amino acids. Early-diverged strains Mr1046, Mr727 and Mr1120 displayed the lowest utilization and growth rates but still preferred carbohydrates (S12 Table).

Among amino acids, glutamic acid (average OD_490_: 1.779) and alanine (average OD_490_: 1.507)-the major amino acid in insect cuticles-were highly utilized by most strains. Mr1-16, Mr2575 and Mr23 efficiently utilized proline, whereas other strains did not. Mr2575 particularly favored pyroglutamic acid, which all recently diverged strains utilized well. Mr1046 and Mr1120 showed little and delayed growth on several amino acids including threonine, and on most carboxylic acids except 2-keto-D-gluconic acid, a generally preferred substrate for *M. robertsii* strains. Quinic acid was highly utilized by most strains (average OD_490_ = 1.845), except Mr1046 and Mr1120 (average OD_490_ = 0.702), indicating an active shikimate pathway for aromatic amino acid production.

Fungi can convert various sugars to the sugar alcohol adonitol as a carbon source(36) and adonitol supported rapid growth of all strains (S18-20 Figs). The amino-sugar N-acetyl-glucosamine, a chitin breakdown product, was among the most highly utilized and growth supporting substrates for all strains. Polymers like glycogen and dextrin, and the primary insect hemolymph sugar, D-trehalose(39), were moderately assimilated by all strains. Other substrates in this cluster (e.g., L-rhamnose, D-tagatose, xylitol, D-arabinose, D-lactose, N-acetyl-galactosamine and N-acetyl mannosamine) less efficiently utilized are rare in insect cuticles or plant cell walls and are not common root exudates. Well utilized sugars such as D-glucose, D-xylose and L-arabinose are major monosaccharides in plant cell walls and root exudates(37).

Unexpectedly, D-galacturonic acid, the major monomer of pectin and a carboxylic acid sugar, supported little or no growth by *M. robertsii* strains (S37 Fig). Blast searching the genomes revealed all strains retained several pectinase families and single homologs (e-value = 0, > 80% similarity) to *Trichoderma reesei* D-galacturonic acid reductases GAR1 and GAR2 (Mr2575 gene accessions: XP_007817691.1 and XP_007824913.1), which mediate D-galacturonate catabolism(38). Cellobiose (the principal breakdown product of cellulose) and gentobiose supported good growth of most *M. robertsii* strains (S18-20 Figs) and can induce cellulase production in some fungi(39). Cellulose is not an FF plate substrate, but *M. robertsii* strains were unable to grow on agar plates containing 2% crystalline cellulose as sole carbon source. Cellobiose and gentobiose are not frequently cited as root exudates suggesting that access to cellobiose and gentobiose may require cohabitation with cellulase-producing microorganisms. The moderately efficient utilization of raffinose may support interactions with plants over longer distances, as seen in Mr2575’s root-directed growth in response to raffinose containing exudate gradients (40).

Maltose, a disaccharide commonly used in *Metarhizium* cultivation and mass production, supported good growth of Mr2575, Mr23, Mr1-16, Mr1878, with weaker and delayed (>72 hours) growth in other strains. Maltotriose (composed of three glucose units), turanose (a sucrose analog), palatinose (a glucose-fructose disaccharide with a α-1,6 glycosidic bond) and sucrose were also better utilized by recently diverged *M. robertsii* strains. All strains grew well on D-sorbitol (average OD_750_: 1.011) and showed growth rates on glycerol (average OD_750_: 0.983) comparable to glucose (average OD_750_: 0.949). Insect hemolymph can contain glycerol at molar concentrations(41).

Early diverged strains sporulated rapidly on an average of 42 substrates by day 7 and on 65 by day 10 (Fig 9). Recently diverged strains Mr2547 and Mr2575 resembled early diverged strains, sporulating on 33 substrates by day 7 and 60 by day 10. In contrast, Mr1878, Mr1-16, and Mr23 sporulated later, on only 6 substrates by day 7 and 18 by day 10. Notably, Mr1-16 failed to sporulate on amino acid carbon substrates, while other strains sporulated on between 3 (Mr23) and 13 (Mr727) by day 10. Mr2575 was among the fastest sporulators on carbohydrates (including plant-related D-cellobiose, D-raffinose, and sucrose) and carboxylic acids, while Mr2547 sporulated rapidly on both carboxylic acids and amino acids (S16 Table). N-acetyl-D-glucosamine was the only substrate on which all strains sporulated by day 10; early diverged strains sporulated on it as early as day six, while Mr1-16 did not sporulate until day nine.

**Fig 9.**
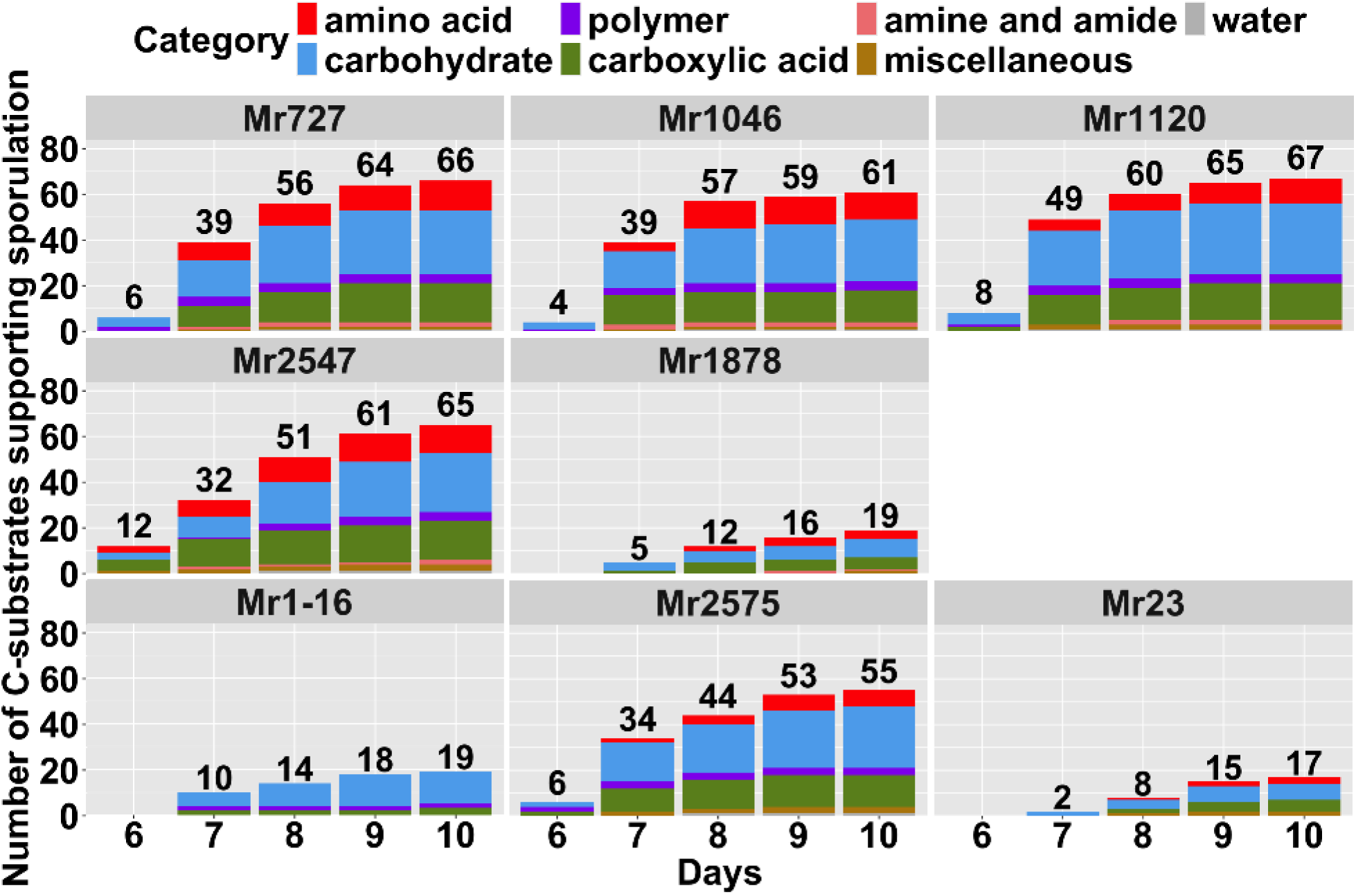
Sporulation of *M. robertsii* on FF MicroPlates. C-substrates were grouped into amino acids, polymers, carbohydrates, carboxylic acids, amines and amides, miscellaneous, and water. Sporulation was monitored for 10 days., and the total number of sporulated C-substrates for each *M. robertsii* strain is indicated above each bar.

Certain substrates reduced metabolic activity and growth compared to water (S17 Table). Mr2547, Mr1-16, Mr2575, and Mr23 were inhibited by glucuronamide, previously found to be an inhibitor of *Trichoderma*(42). Mr2547 and Mr23 were inhibited by D-saccharic acid. Mr1-16 and Mr23 showed reduced metabolic activity and growth on N-acetyl-D-galactosamine, while Mr2547 and Mr2575 showed growth inhibition only. These patterns suggest that certain substrates act as metabolic stressors or are poorly metabolized by specific strains. Mr1-16 and Mr23 showed lower utilization but higher growth on D-arabinose compared to water, suggesting that this substrate may support biomass accumulation through non-respiratory or alternative metabolic pathways. *M. robertsii* strains failed to sporulate on 15 substrates, including glucuronamide, D-galacturonic acid and N-acetyl-D-galactosamine that support little of no growth by some or all strains, as well as α-D-glucose, salicin, maltotriose, maltose, glycerol, gentiobiose, D-mannose, D-mannitol, D-arabitol, amygdalin, adonitol, and Tween 80.

We analyzed the relationships between carbon substrate utilization, virulence, and endophytic ability (S18 Table and S38 Fig). Strains that utilized and grew well on these C-substrates also exhibited high insect virulence and strong germination on fly wings, roots, YE, RE, and 1% glucose (S38 Fig). This suggests these strains are opportunistic fungi capable of efficiently exploiting diverse substrates, exhibiting rapid insecticidal activity, and maintaining strong plant associations.

## Discussion

The eight *M. roberstii* strains form a monophyletic group divided into early and recently diverged lineages, reflecting distinct evolutionary paths between strains that have been diverging from a common ancestor for a long time and strains that have recently diverged. Early diverged strains (Mr727, Mr1046, Mr1120) are characterized as slow-killing pathogens that produce abundant spores, while recently diverged strains (Mr2547, Mr1878, Mr1-16, Mr2575, Mr23) are fast-killing plant root colonizers. By integrating Biolog FF microplate data with experiments on complex plant and insect-related nutrients, we compared carbon and nitrogen utilization across these strains. The observed differences in growth patterns on various carbon sources suggest significant ecological and genetic divergence, particularly in their approach to plant and insect components.

Early diverged strains, especially Mr1046, utilized a narrower range of nutrients and exhibited slower growth on most of the others, consistent with adaptation to nutrient limitations imposed by a single ecological niche. However, they sporulated rapidly and profusely on a broad array of substrates, indicating a strategy focused on persistence through spore production. In contrast, recently diverged strains showed rapid growth on a broad spectrum of nutrients, including insect cuticles and plant roots, demonstrating greater ecological versatility as pathogens, endophytes and saprophytes. This suggests that recently diverged strains prioritize fast colonization and growth to exploit new or variable niches, while early diverged strains emphasize spore production for survival in consistent or resource-limited environments. Nutritional flexibility is critical for saprobic fungi adapting to the heterogeneous soil environment (43). Furthermore, we observed germination speed and growth on insect cuticles and plant roots strongly predict and influence virulence and endophytic potential, with faster germination correlating with quicker insect killing.

Plant roots release a variety of low molecular weight compounds, such as simple sugars (D-glucose, D-maltose, D-mannose, D-fructose, D-xylose, D-ribose, L-arabinose, D-raffinose, sucrose), amino acids, organic acids and phenolics, that shape microbial interactions in the rhizosphere(44)(45). Sugars are particularly important due to their role as a readily available energy source and the carbohydrates listed as root exudates were all very efficiently utilized by Mr1-16, Mr2575 and Mr23 potentially enhancing their plant association capabilities. For example, Mr1-16 begins growing in root exudate (RE) solutions within six hours, aligning with its superior root attachment and germination ability. Many substrates where Mr1-16 showed the fastest growth rate (S36 Fig), including xylose, raffinose, sucrose, fructose, mannose, are major exudate components, while D-gluconic acid is a major hemicellulose component. This ability to germinate in root exudates enhances the potential for associating with plants at a distance, as we previously demonstrated that Mr2575’s grows towards roots following a concentration gradient of raffinose in root exudate (40).

It is unclear which environmental factors can shift a fungal pathogen to a new niche(16). Specifically, for different *Metarhizium* strains we lack adequate knowledge about individual associations with different plants in nature and the mechanisms behind these associations. Most insights come from studies with Mr2575, which shows consistent field and laboratory behavior(24), indicating that *in vitro* studies on individual strains can reflect field conditions. The variation in degrees of root colonization across *M. robertsii* strains would likely produce a spectrum of plant responses. Mr2575 significantly enhances plant growth in the field(24), suggesting that the superior root colonizer Mr1-16 particularly merits further study as a biofertilizer.

Examining interactions with the *Drosophila*, immune system revealed that virulence correlates with the timing of the host immune response: fast-killing strains (e.g., Mr1878, Mr2575) induce early immune activation, whereas slow-killing Mr1046 triggers a delayed response. Using immune-deficient fly mutants, we found that components of the Toll pathway and melanization affect *M. robertsii* virulence but do not explain the observed sexual dimorphism in susceptibility, although *Psh* and other determinants of the Toll pathway account for sexual dimorphism against some other pathogens(46,47). The host’s genetic background influences fungal proliferation for some strains (e.g., Mr1046), but recently diverged strains (Mr2547, Mr2575, Mr1-16) produce few CFUs in *D. melanogaster* or *B. mori*, showing proliferation patterns that are independent of host genotype. Mr23, closely related to Mr2575, shares most phenotypes, but differs by producing more CFUs in insects, germinating less effectively in 0.0125% yeast extract and some other sparce nutrients, and being less virulent to the resistant *Drosophila* line RAL 808. Thus, even among closely related toxin producing strains, a high CFU count does not always equate to high virulence. Compared to Mr2575, the fitness of Mr23 may depend more on a susceptible host and nutrient rich conditions, such as those found in a host. Within-host fungal proliferation, as estimated by CFUs, correlates positively with sporulation on insect cadavers and negatively with saprophytic growth and germination rate on fly wings, indicating a trade-off: early diverged strains kill slowly but their abundant proliferation in the hemolymph before death maximizes reproductive output post-mortem.

The balance between sporulation and hyphal proliferation likely shapes establishment of *Metarhizium* in the soil and rhizosphere. A non-pathogenic but root colonizing mutant of Mr2575 survives better in soils and plants than an insect-pathogenic mutant that could not adhere to roots, confirming roots as a primary habitat(24). Consequently, recently diverged strains may rely more on root colonization for reproduction, reducing selection for heavy sporulation on insects. Extensive hyphal growth on living insects, and particularly on cadavers, as seen in Mr1-16, may facilitate the colonization of nearby plants. This strain resembles the “creeper” phenotype described by Angelone et al (48), that explores habitats, whereas early diverged non-plant colonizing strains act as “sleepers”, producing spores that await hosts. According to our study, these phenotypes align with differences in metabolic plasticity.

It is also plausible that plant and insect surfaces exhibit shared characteristics, enabling adaptations to one to facilitate colonization of the other. The endophytic ability of *M. robertsii* strains is tied to germination and growth on insect cuticles, suggesting that the ability to exploit a wide array of nutrients underpins both insect virulence and plant association. Similarly, ecological and nutritional versatility predicts rhizosphere fitness in bacteria(49), a trait abundantly present in recently diverged *M. robertsii* strains. Consequently, the evolutionary gap between opportunistic plant associations and enhanced virulence to insects may be small, with overlapping genetic “toolkits” for entomopathogenicity and plant colonization. It is conceivable that metabolic flexibility evolved in recently diverged *M. robertsii* strains to enhance competition for limited nutrients across in a specific environment (soil, plants or insects) and has been directed to support multiple other associations. Supporting this, Mr2575 spores fail to infect insects when supplemented with additional nutrients, indicating that insect associations serve nutritional purposes (50).

Nutrition and ecological roles are commonly used to delimit fungal taxa (51–53), implying early and recently diverged strains could be classified as different species. However, reclassifying historical *Metarhizium* species into narrower definitions may lack biologically or practical benefit, especially since current *Metarhizium* PARB clade phylogeny, based on morphology and specific DNA sequences(21), does not fully capture ecological niches.

Whether the shift from saprophyte to plant endophyte is simple for some fungi remains debated(22). In *M. roberstii,* lifestyle transitions are linked to a nutritional shift that also favors the rapid killing of insect hosts and plant associations, indicating these traits may be transient. Therefore, further research into nutrient variability in soil, insects and plants—and its role in driving nutritional shifts—is essential for understanding ecological plasticity in this species. Supporting the notion that lifestyles are transient are differences in the expression of destruxins. Non-toxigenic early evolved *Metarhizium* species such as *M. acridum* lack the destruxin (dtx) cluster (54,55). Several studies suggest that the dtx gene cluster was recruited into the ancestor of the PARB clade of closely related *Metarhizium* species (*M. pingshaense, M. anisopliae, M. robertsii,* and *M. brunneum*) by horizontal helping set the PARB clade on the trajectory to a broad host range(56). Despite inheriting a destruxin gene cluster, Mr1046 and Mr727 exhibit a non-toxigenic phenotype which mimics the phenotype of *M. acridum* by killing hosts through nutrient depletion and mechanical disruption of tissues due to extensive fungal growth(54,55).

In summary, this study confirms *M. roberstii* as a valuable model for exploring fungal ecological plasticity and potential ongoing evolution towards plant endophytism. While further strain analysis is needed, this research highlights clear nutritional mode shifts corresponding to ecological differences between early and recently diverged lineages within this monophyletic group.

## Data availability

The genome sequences used to assemble phylogenies have been deposited in the National Center for Biotechnology Information (NCBI) database [NCBI BioSample accession: SAMN54551730 (*Metarhizium robertsii* ARSEF 1046), SAMN54551731 (*Metarhizium robertsii* ARSEF 1120), SAMN54551729 (*Metarhizium robertsii* ARSEF 727), SAMN54551733 (*Metarhizium robertsii* ARSEF 1878), SAMN54551732 (*Metarhizium robertsii* ARSEF 2547), SAMN54551735 *Metarhizium robertsii* ARSEF 2575) and SAMN54551734 (*Metarhizium robertsii* Mr1-16).

## Acknowledgments

We thank Novogene for genome-sequencing services. This project was supported by the USDA National Institute of Food and Agriculture and Agricultural Research Service Biotechnology Risk Assessment Grant Program (2022-33522-38272 to RJS) and jointly supported by the Plant Biotic Program of the National Science Foundation and the USDA National Institute of Food and Agriculture (DEB 1911777 to RJS). The funders had no role in study design, data collection and analysis, decision to publish, or preparation of the manuscript. Figures were created using BioRender.

## Materials and Methods

### Growth Conditions

RAL 808, RAL 321, Drosomycin-GFP (Drs-GFP), Δ *PPO^1,2,3^*, and its isogenic control lines (*W^1118^*) were obtained from the Bloomington *Drosophila* Stock Center (https://bdsc.indiana.edu/index.html) with Bloomington stock number indicated in parenthesis. RAL 808 (#28238) and RAL 321 (#29655) are wild-type fly lines in the *Drosophila* Genetic Reference Panel and have been previously studied in our lab(23,27). RAL 808 is comparatively resistant and RAL 321 is comparatively susceptible to Ma549(23). Drs-GFP (#55707), Δ *PPO^1,2,3^* (#68387), and *W^1118^* (#5905) have been previously described(33,57). The Persephone-KO line fly line (Δ*Psh*) and its isogenic control line (*w*^A5001^) were kindly donated by Dominique Ferrandon (University of Strasbourg, Strasbourg, France)(57). All fly lines were reared on Cornmeal-molasses-yeast-agar medium containing Tegosept and Propionic acid (Genesee Scientific) at 24 ± 3 °C.

*Tenebrio molitor (T. molitor)* adults were obtained from the Carolina Biological Supply company (https://www.carolina.com). *Popillia japonica* (*P. japonica)* adults were collected using Japanese beetle traps with pheromone lures from June to mid-July.

Seven *M. robertsii* strains (ARSEF 1046, 1120, 1878, 23, 2547, 2575 and 727) were obtained from the USDA Entomopathogenic Fungus Collection (Ithaca, NY, USA). Mr1-16 was isolated from a rhizospheric soil sample collected in Burkina Faso by Dr. Etienne Bilgo, a former Ph.D. student in the laboratory. Fungal cultures were transferred from -80°C stock tubes twelve days before each bioassay and maintained on potato dextrose agar (PDA) at 27°C. Plasmid construction and transformation of GFP or mCherry fluorescent *M. robertsii* strains have been previously described (58).

### Bioassays

To prepare inocula, conidia were harvested from 10-14-days-old potato dextrose agar plates, suspended in sterile distilled water, vortexed for 2 min, and filtered through Miracloth (22-25µm) (Andwin Scientific) to remove mycelia. Spore concentrations were determined using a Neubauer hemocytometer and adjusted to the required concentration using water.

Flies were reared on standard fly food without antibiotics for two days. The infection method was described by Wang et al. (2017)(23). Briefly, we assayed five tubes of ∼30 flies (aged 2-5 days old) per sex and per line with a spore suspension (2.5x10^4^ spores/mL of water) made using 8- to 15-day-old cultures of each *M. robertsii* strain. The infected flies were maintained at 27°C with 85% relative humidity and a 12-hour light-dark cycle, and mortality was recorded twice a day for 14 days. Control flies were treated with distilled water for the bioassay.

Adult *T. molitor*, and *P. japonica* were used to evaluate *M. robertsii* intraspecific variation against a different insect order. At least 10 *T. molitor* or *P. japonica* adults were randomly selected from each colony per replicate, and each treatment was replicated three times. They were infected by 20s submergence in 15mL of an *M. robertsii* spore suspension and then transferred to a container containing organic apples as their diet. They were maintained at 27 °C with 85% relative humidity and a 12-hour light-dark cycle and mortality was recorded every 12 hours for 14 days. The control groups were established using distilled water in place of spore suspensions.

*T. molitor* and *P. japonica* infections were performed using a spore suspension of 2x10^7^ spores/mL. To more precisely determine the LD_50_’s of Mr1046 and Mr1878 on *P. japonica*, various spore suspensions (1x10^5^, 5x10^5^, 1x10^6^, and 5x10^6^ spores/mL) were used, and LD_50_‘s were calculated using the ‘drc’ R package.

The time to death was calculated for each replicate and expressed as the LT_50_. Each treatment’s LT_50_ and standard error (or standard deviation if there were only three replicates) were calculated using Rstudio (version: 4.3.1). Kaplan Meier survival curves were generated using Prism GraphPad (version: 8.4.0, GraphPad Software, Boston, Massachusetts USA, www.graphpad.com), and log-rank tests were performed to compare the survival among treatment groups.

### Fungal growth, latent period, and sporulation capacity

Drs-GFP females were selected for a time-course bioassay of fungal growth in the hemolymph using previously described protocols(23). Every 12 h post-infection, ten flies of each sex were individually homogenized with 45 μl of 0.1% Tween 80. The homogenate was spread onto PDA plates supplemented with oxbile, cetyltrimethylammonium bromide, oxytetracycline, streptomycin, penicillin, chloramphenicol, and cycloheximide.

Flies, *T. molitor*, and *P. japonica* were used to determine the post-mortality pathogen parameters. Ten insect cadavers were randomly collected and transferred to containers with damp cotton balls to provide humidity. At twelve-hour intervals, we recorded the time interval between the death of the host and the duration until the appearance of spores (latent period).

Twenty days after the onset of sporulation, each cadaver was transferred to a tube containing 0.5% Tween 80 (500uL for flies; 2mL for *T. molitor* and *P. Japonica*) and vortexed for two minutes before counting spores using a hemocytometer. Male and female flies were analyzed separately.

The correlations between different fungal life traits were determined using Pearson correlation, with Bonferroni correction applied to adjust for multiple testing.

### Germination on *Drosophila* wings

Fly cadavers were sterilized in 70% ethanol for 5 seconds and air-dried in a sterile flow hood. Ten cadavers per replicate were transferred onto a 0.8% agar plate and co-inoculated with mCherry-tagged Mr2575 (the type strain of *M. robertsii*) and a GFP-tagged *M. robertsii* strain (3.5x10^5^ spores/mL of each strain) and incubated at 27°C and 85% relative humidity. After 12h, the wings were removed and transferred onto a slide, and the percent spore germination was observed using a compound microscope (Zeiss MicroImaging GmbH 37081) with filter sets for GFP (38 HE GFP) or mCherry (20 Rhod). Additionally, ten germlings per strain were randomly selected, and the germling length was measured using ZEISS ZEN software (version: 3.0).

Identify destruxin (dtxs) production in *M. robertsii*

Each *M. robertsii* strain was suspended in 5mL of 0.01% Tween 80 and vortexed for three minutes to separate spores. The spore suspension was then transferred into 45mL of Sabouraud dextrose broth with 0.2% yeast extract and incubated for seven days at room temperature (24-25 °C) in a shaker at 200 RPM(29). The mycelia were filtered and collected, and standard amounts of mycelia (0.5g) were transferred to 30mL of Czapek-Dox broth with peptone (5g/L) and shaken (200 RPM) for seven days at room temperature. The fungal exudate was filtered (pore size: 0.22um) and stored at -4°C. The DTX-KO Mr2575 strain (59) was used as a negative control.

*Bombyx mori* (*B. mori*) eggs were obtained from the Carolina Biological Supply company (https://www.carolina.com) and reared on the silkworm artificial diet to fifth instars. Each fifth instar *B. mori* was injected with 50uL of the filtrate through the base of the second last proleg. A *B. mori* caterpillar was considered knocked down if it could not right itself within five seconds following inversion, and the knockdown time for each *B. mori* was recorded. Each treatment had three replicates of ten insects.

To examine fungal infection morphology, infected larvae were selected at the moribund stage (inability to right themselves within five minutes on three consecutive attempts and died within 12 hours). Moribund *B. mori* instars were observed using a compound microscope (Zeiss MicroImaging GmbH 37081) with filter sets for GFP (38 HE GFP) or mCherry (20 Rhod). To estimate the number of blastospores present in each strain, 3μL of hemolymph from each of five biological replicates per strain was examined under a compound microscope (Zeiss MicroImaging GmbH 37081) with GFP (38 HE GFP) or mCherry (20 Rhod) filters. Some images were captured from the ZEN imaging software interface using the Windows screenshot tool.

### Fungal growth and sporulation on agar

Growth of the eight *M. robertsii* strains was measured on three different media: potato dextrose agar (PDA), Sabouraud dextrose agar + 1% yeast extraction (SDA+1%YE), and Czapek-Dox agar (CZA) 27± 1 ∘C. Both PDA and SDA+1%YE are complex media containing organic sources of nitrogen, whereas CZA contains glucose and sodium nitrate as the sole carbon and nitrogen sources. The SDA medium is particularly nutrient-rich and contains carbohydrates, casein, and peptone. The added yeast provides fungi with proteins, amino acids, vitamins etc. Plates were inoculated with sterilized toothpicks dipped in spore suspensions of each *M. robertsii* strain, and colony diameter and morphology were observed daily for seven days. Replicates were generated using five colonies per plate and three plates per strain. Correlations with other fungal traits were calculated using Pearson’s correlations based on growth rate from day six to day seven on PDA at 27±1°C.

The ability of each strain to grow on 1% crystalline cellulose (Sigma, C-6288), 1% apple pectin (Sigma, P-2157, reported by the manufacturer to contain 76% D-galacturonic acid and not contain sucrose or other sugars), 1% D-glucose (Sigma, G-7528) or 1% D-galacturonic acid (Sigma, 48280) was determined in a minimal Noble agar (1.5%) medium containing 0.2% NaNO_3_, 0.1% KH_2_PO_4_, 0.05% MgSO_4_, and FeSO_4_, ZnSO_4_, CuSO_4_ and MnCl_2_ each at 10mg per 100ml. The media were adjusted to pH 8 (pectin) or 5.5 (other media) before autoclaving.

To compare *M. robertsii* sporulation on agar, each *M. robertsii* strain was suspended in 10mL 0.01% Tween 80 and vortexed for two minutes. Each PDA plate was inoculated with 500uL spore suspension spread using a spreader. Fungal plates were stored at 27 °C for 18 days. Five agar plugs were obtained from each fungal plate using a soil sample probe (sample diameter: 2.2 cm) and individually transferred into a 50mL falcon tube with 10mL 0.01% Tween 80. After vertexing for three minutes, released spores were counted using a hemocytometer. Each treatment consisted of four plates, with 20 replicates in total.

### Quantification of Drosomycin production

The production of Drosomycin in the Drs-GFP flies has been confirmed to be associated with fluorescence intensity using real-time PCR(32). Ten live female Drs-GFP flies were selected per day for each treatment group. Their heads were removed and the bodies were individually transferred into 40 ul of TBS +1mM Dithiothreitol buffer. Each fly was homogenized using an electric homogenizer for 20-30 seconds and centrifuged for one minute at 10000 rpm. Thirty microliters of supernatant from each tube was transferred into a 384-well microplate (Corning Inc.). Fluorescence was quantified using a FilterMax F5 microplate reader (Molecular Devices), and GFP fluorescence was measured in fluorescence units. The control group consisted of flies that were treated with water. Welch’s t-test was performed to compare treatments, and Pearson correlations were performed to assess the relationship between Drosomycin production and other traits.

### Attachment and germination ability on plant roots

Three-day-old seedlings of *Arabidopsis thaliana* were placed in 0.8% w/v agar plates, with their roots suspended in a longitudinal well cut through the agar. The wells contained *mCherry*-tagged *Mr*2575 plus a GFP-tagged *Metarhizium* strain (2.5 x 10^5^ spores/mL of each strain in 2 mL water). Mr2575 was used as a reference standard for comparing each GFP-tagged *M. robertsii* strain. After 12 hours, each seedling was transferred to a microscope slide. The number of spores and germlings of each strain on each root was determined using a compound microscope (Zeiss MicroImaging GmbH 37081) with filter sets for GFP (38 HE-GFP) or mCherry (20 Rhod). A spore was considered to have germinated when it had a germ tube at least half the length of the spore.

### Plant association in soil environment

We assessed the ability of Mr1046-GFP, Mr1-16-GFP and Mr2575-mCherry to associate with plants in a natural soil environment. Fifth instar *Manduca sexta* larvae were injected with 50 μL of a spore suspension at the base of their second to last pair of prolegs. Infected *M. sexta* were reared on damp filter paper for the first 12 hours post-injection to maintain 90% humidity, then transferred to 50 mL tubes containing standard *Manduca* diet.

Following larval death, cadavers were placed in petri dishes and covered with soil and grass roots collected from a wooded area near a river. Observations were conducted using a Zeiss MicroImaging GmbH 37081 compound microscope equipped with a GFP filter set (38 HE GFP). Cadavers infected with Mr2575-mCherry or Mr1046-GFP were examined at 30× magnification, and the number of nematodes on each cadaver was counted (15 replicates per *M. robertsii* strain).

### Germination ability in different nutrient sources

Spores of each strain were transferred into 2mL of either 0.0125% yeast extract solution (YE), 0.1% YE, 1% glucose solution (1% glucose, 0.1% NaNO_3_, 0.05% KH_2_PO_4_, 0.05% MgSO_4_), 0.0125% plant root exudate (RE) or 0.1% RE. Plant root exudates were obtained by culturing wheat roots in sterile water for 30d. The exudate was freeze-dried and resuspended in distilled water to obtain 0.1% and 0.0125% RE solutions. The root germination rates and germling hyphal lengths were calculated after 4.5, 6, 12, and 18 hours from photographs taken using a Leica DMIRB inverted microscope. The germination rate at 18 hours post-inoculation was used to perform Spearman correlations to assess the relationships between germination in solutions and other traits.

### Filamentous Fungi (FF) MicroPlate Biolog™ bioassay

The metabolic profiles of the eight *M. robertsii* isolates were measured using FF plates (Biolog, Inc., Hayward, CA, USA) containing 95 different carbon substrates (C-substrates) and tetrazolium. Conidia were transferred into 2mL of 0.01% Tween 80, washed twice with distilled water, and resuspended by vortexing for 10s in FF Inoculating Fluid (FF-IF, Biolog) to obtain a final spore suspension of 1×10^5^ spores/mL. Then, 100uL of spore suspension was dispensed in each well of an FF plate which was incubated at 27°C in the dark. As described in the manufacturer’s protocol, absorbance at 490 nm (substrate utilization) and 750 nm (fungal growth intensity) were measured every 12 hours for a total of 120 hours using a microplate reader (Agilent, BioTek Epoch 2). The control plate was inoculated with water only. Each treatment group was repeated three times.

The 95 C-substrates were separated into six major groups: amines and amides, amino acids, carbohydrates, carboxylic acids, water, and miscellaneous. Similarities in carbon utilization patterns among *M. robertsii* strains were illustrated using hierarchical cluster analysis (pvclust package in R).

