## Supplementary Figures S1-S38 for "Origins and metabolic evolution of multifunctionality in *Metarhizium robertsii*: linking phenotypic diversity to ecological niche plasticity"

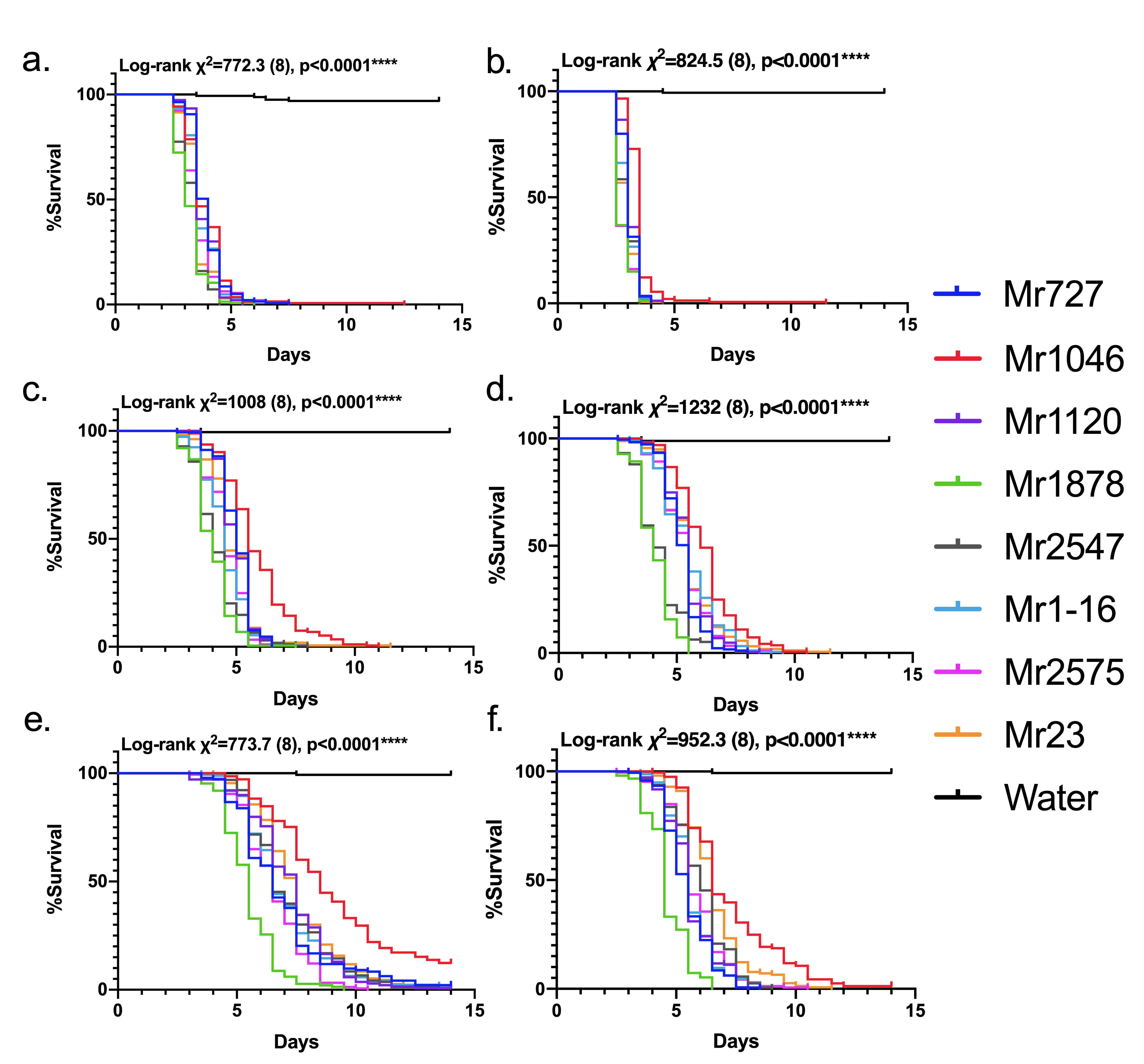

**S1 Figure. Survival curves (Kaplan-Meier) of three *Drosophila melanogaster* lines infected with *M. robertsii* strains.** Log-rank tests were used to determine if there were significant differences among treatments. a) Male RAL 321, b) Female RAL 321, c) Male Drs-GFP, d) Female Drs-GFP, e) Male RAL 808 f), Female RAL 808.

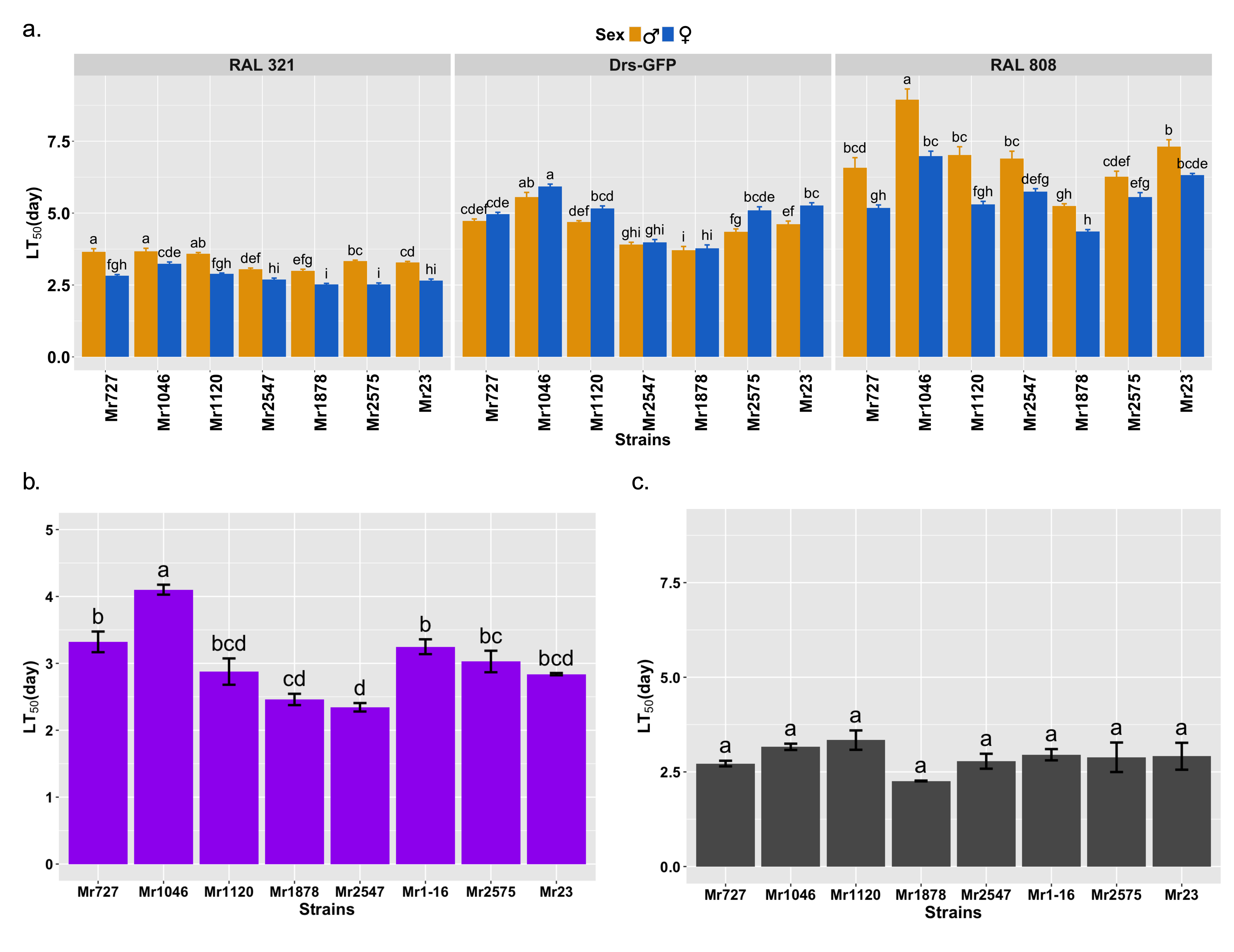

**S2 Figure. LT_50_ values (mean ± SE) of eight *M. robertsii* strains tested against a) three *Drosophila melanogaster* lines (RAL 321, Drs-GFP, and RAL 808), b) *T. molitor*, and c) *P. japonica*.** Different lowercase letters at the top of each bar indicate significant differences by Tukey’s test (One-way ANOVA or Two-way ANOVA, P < 0.05).

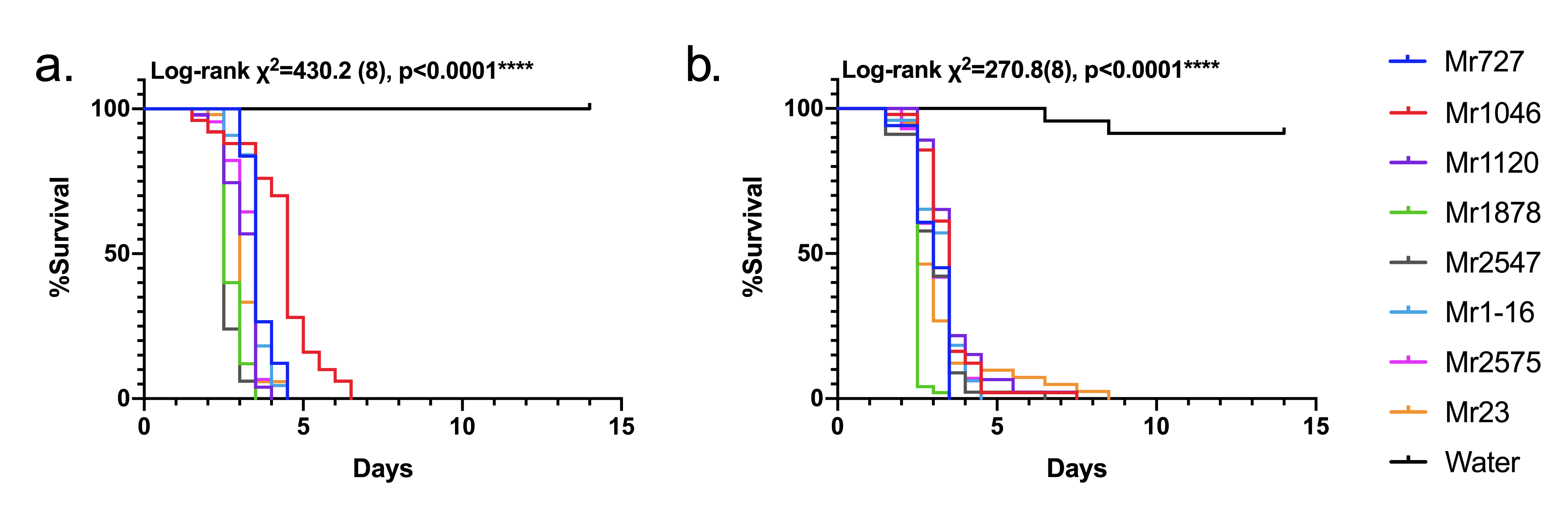

**S3 Figure. Survival curves (Kaplan-Meier) of *Tenebrio molitor* (a) and *Popillia japonica* (b) infected with *M. robertsii* strains.** Log-rank tests were used to determine if there were significant differences among treatments.

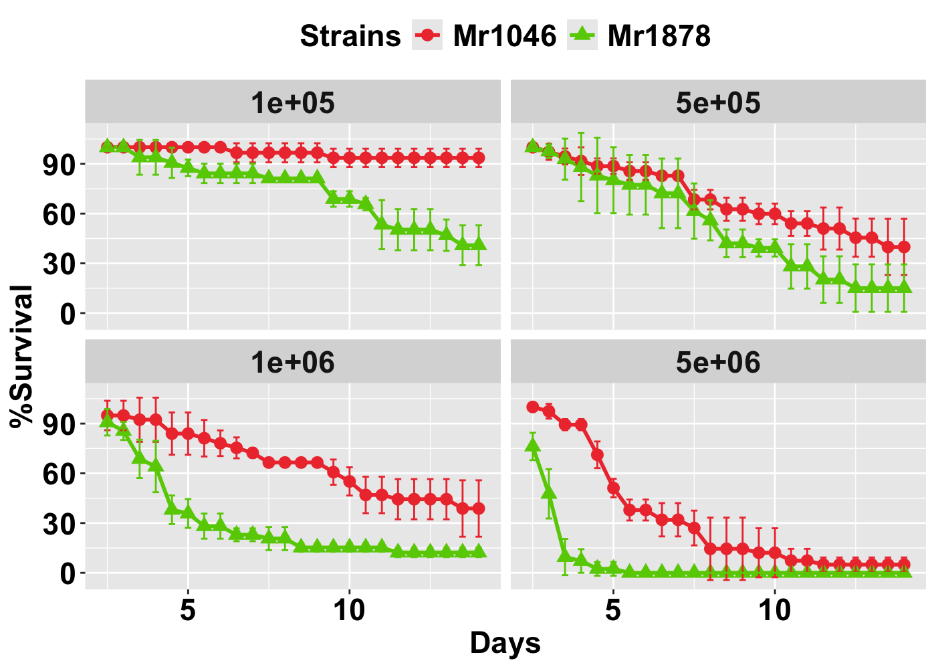

**S4 Figure. Survival curves of *Popillia japonica* infected with either Mr1046 or Mr1878 using various (1x10^5^, 5x10^5^, 1x10^6^, 5x10^6^ spores/mL) spore dosages.** Each treatment was replicated three times.

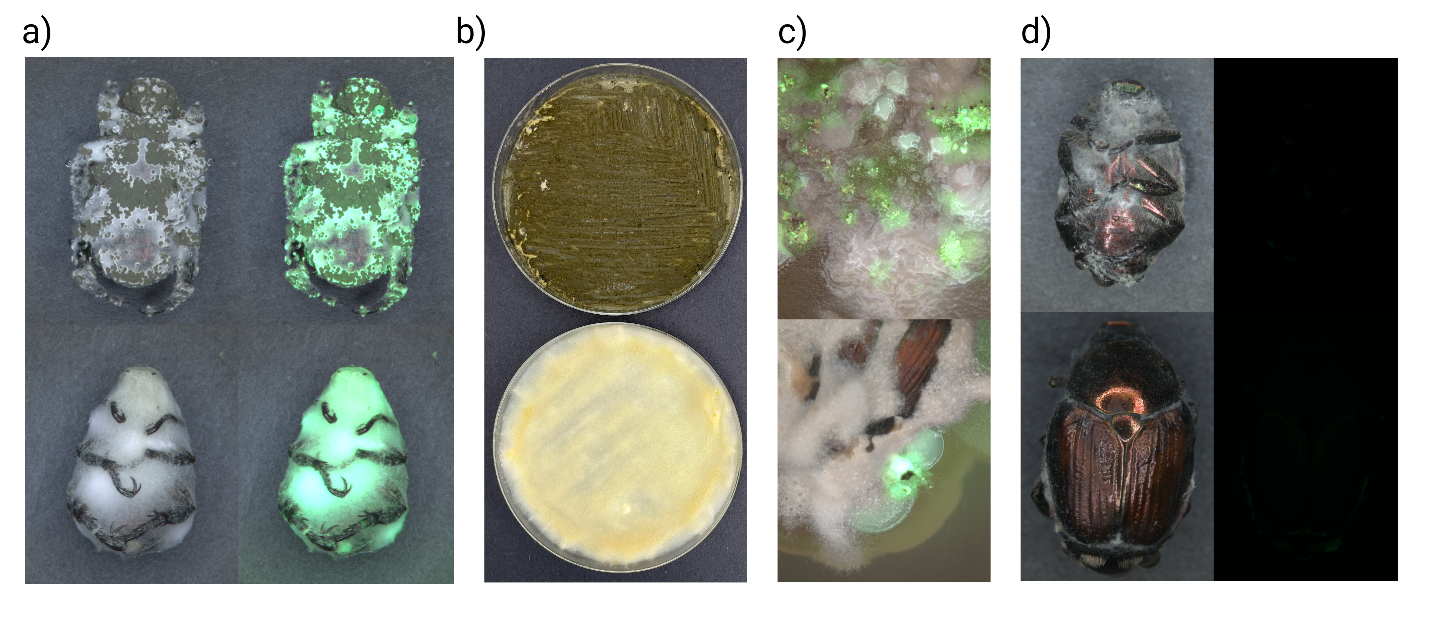

**S5 Figure. Sporulation capacity of *M. robertsii* strains Mr1046-GFP and Mr1878 or Mr1878-GFP.** a) Bright field images (left) and fluorescent images (right) showing the sporulation capacity of Mr1046-GFP (top panels) and Mr1878-GFP (bottom panels) assessed on their original host, *P. japonica*. F**luorescence images** were taken with a filter set for GFP (38 HE GFP). b) Sporulation of Mr1046 (top) and Mr1878 (bottom) on potato dextrose agar (PDA). c) Hemolymph (top). and cuticle (bottom) sampled from P. japonica co-infected with wild-type Mr1878 (non-fluorescent) and green fluorescent Mr1046-GFP. Hemolymph (3uL) was plated out on PDA. Cuticle fragments from cadavers were dissected and placed on PDA. Both Mr1878 WT and Mr1046-GFP can penetrate the insect cuticle, with Mr1878 exhibiting greater penetration (bottom) and better hemolymph colonization (top) than Mr1046-GFP. d) ventral (top) and dorsal (bottom) view of cadaver resulting from mixed infections with wild-type Mr1878 and Mr1046-GFP. The dark panels to the right were taken with a filter set for GFP (38 HE GFP). The absence of GFP signal (green) suggesting cadavers are predominantly covered by non-fluorescent Mr1878.

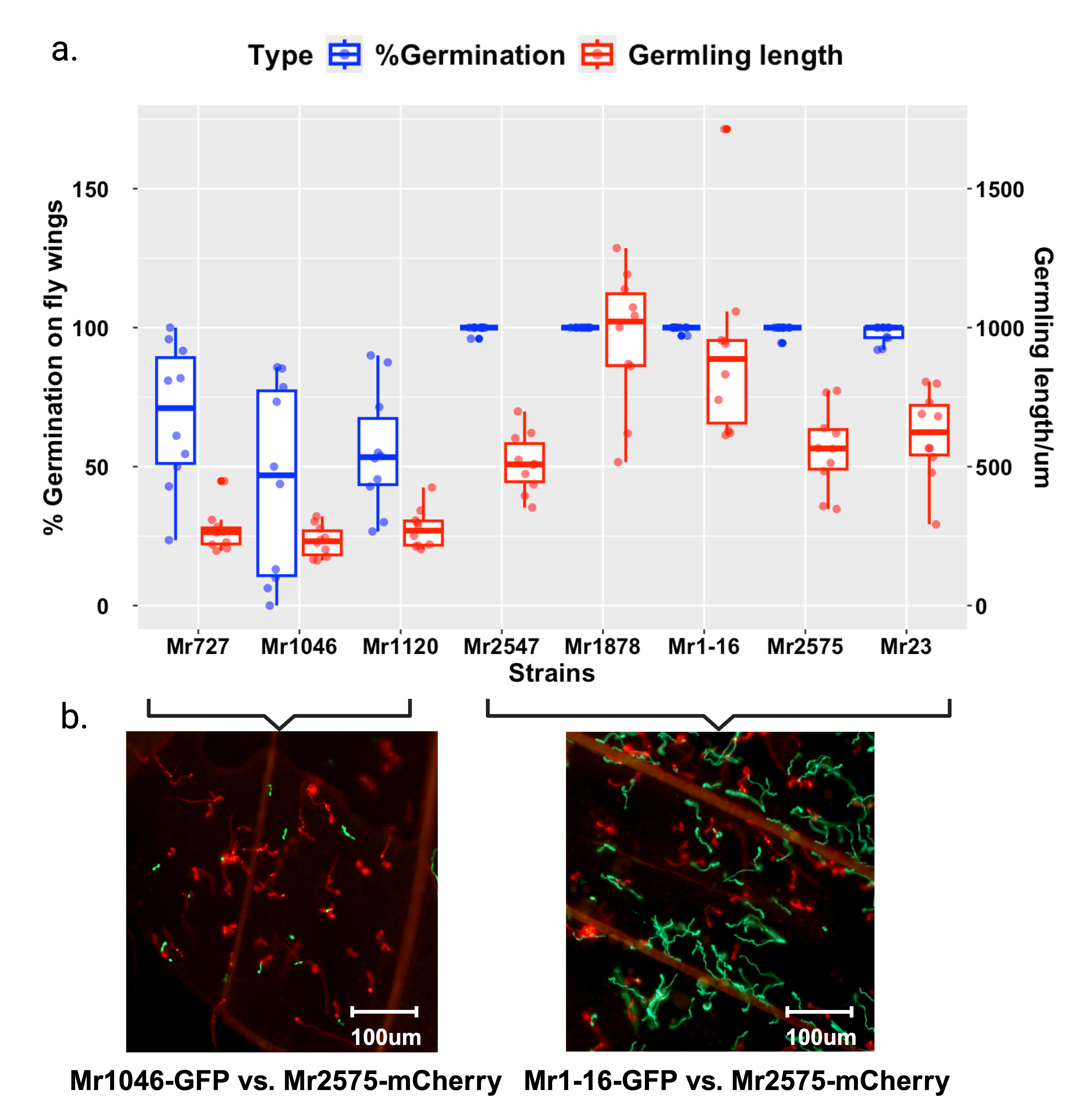

**S6 Figure. Percent germination and germling length of *M. robertsii* strains on insect cuticle after 12 hours incubation at 27 °C.** a) Box plots of percent germination and germling length on fly wings after 12 hours at 27 °C (n = 10 per treatment). b) Compared to Mr2575-mCherry, early diverged, slow-killing Mr1046-GFP (left) showed less attachment (resulting in fewer spores per visual frame), lower germination after 12 h, whereas recently diverged Mr1-16-GFP (right) exhibited higher germination rates and longer germ tubes.

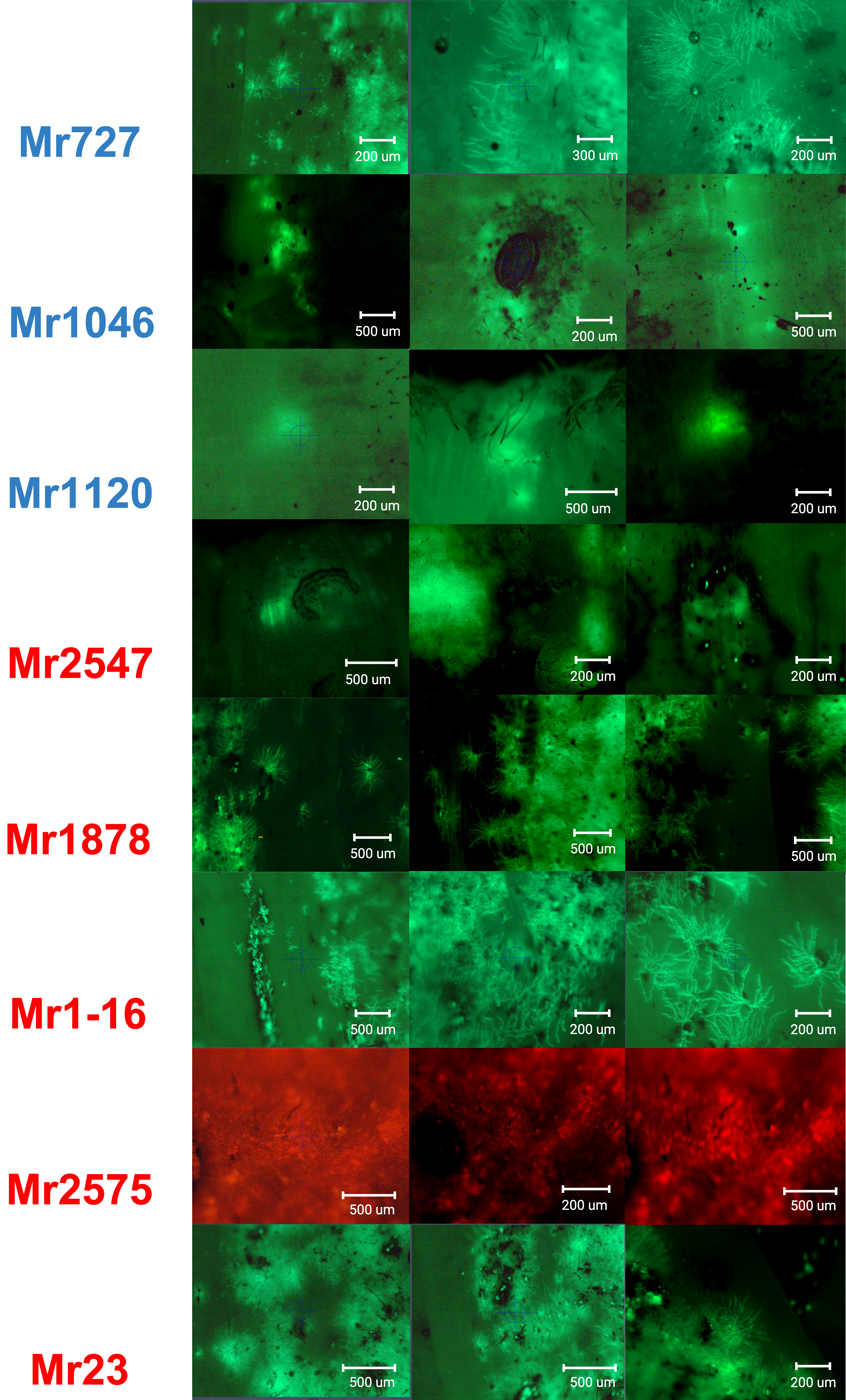

**S7 Fig. Growth of mCherry- or GFP-tagged M. robertsii on or beneath** **B. mori cuticle following topical infection.** Recently diverged strains exhibited extensive growth on the host, with Mr1-16 showing the most hyphal development. In contrast, early diverged Mr1046 and Mr1120 showed little growth. Mr727 displayed growth similar to the recently diverged strains, with extensive hyphal proliferation on and beneath the cuticle, although ungerminated conidia on the cuticle suggest reduced germination. Names of early diverged strains are shown in blue and those of recently diverged strains in red.

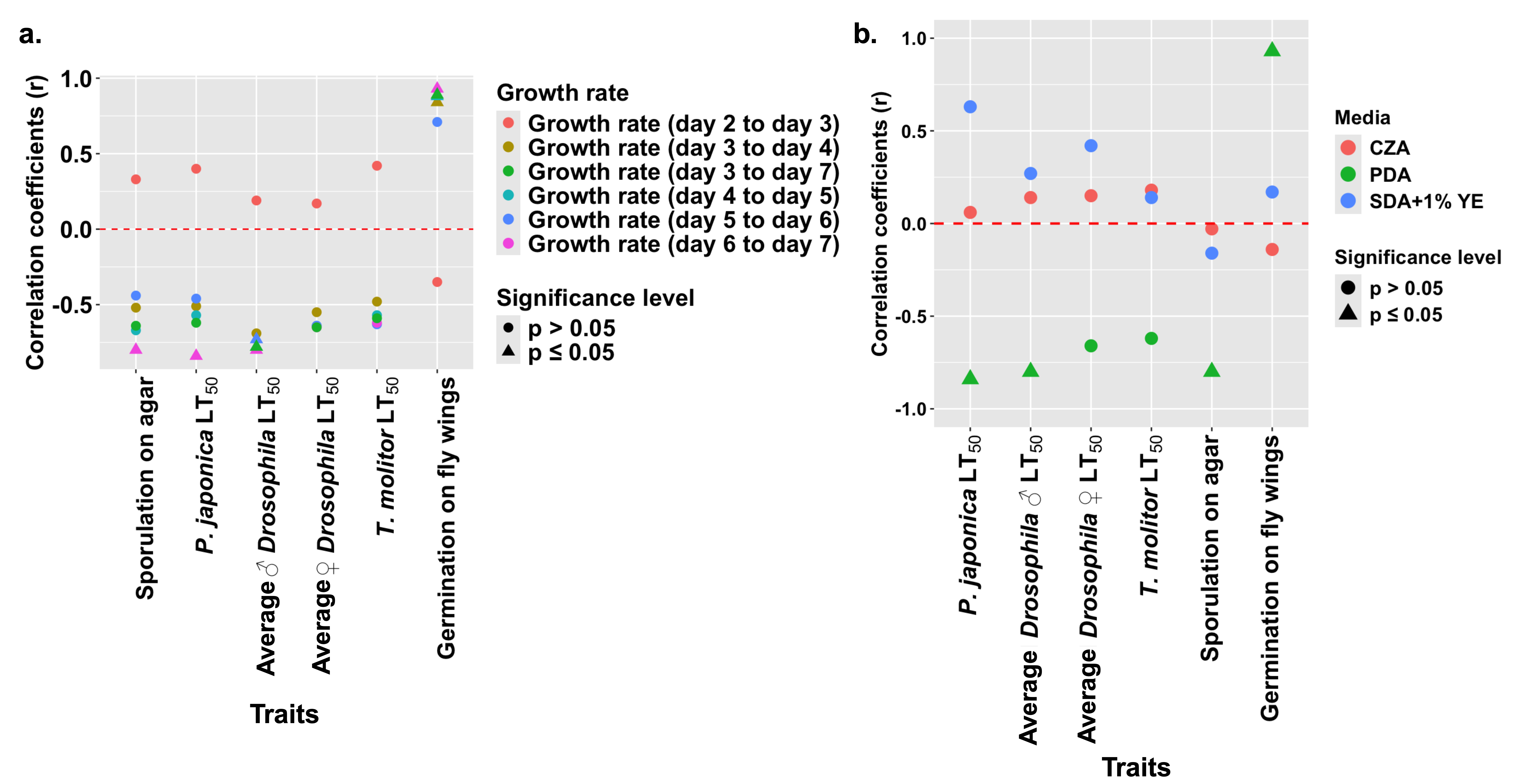

**S8 Figure. a) Correlation coefficients (r) between the growth rate of *M. robertsii* strains on agar, virulence to flies (indicated by LT_50_ values), germination ability on fly wings (indicated by % germination), or sporulation capacity on agar. b) Correlation coefficients (r) between the growth rate of *M. robertsi* strains from day 6 to day 7 on three media and their virulence to flies (average LT_50_ values of three *Drosophila* lines), % germination on fly wings, or sporulation capacity on agar.** The media used are PDA: Potato Dextrose Agar PDA; SDA+1% YE; Sabouraud Dextrose Agar + 1% Yeast Extract; CZA: Czapek-Dox Agar.

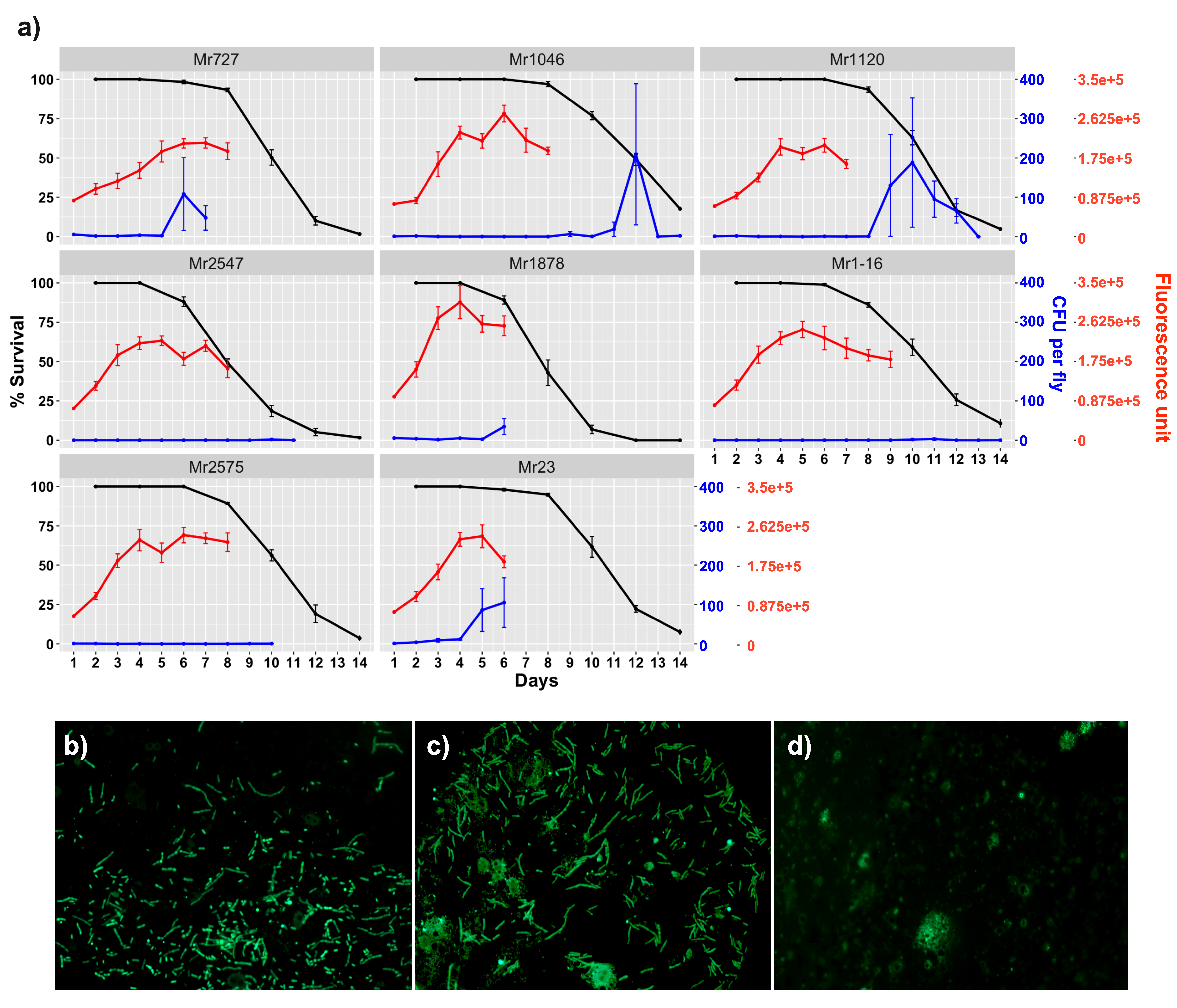

**S9 Figure. The survival curves (black), immunofluorescence responses (red), and fungal proliferation (blue, measured by the number of colony-forming units per fly) in female Drs-GFP flies infected with different *M. robertsii* strains.** a) Data (mean ± SE) was measured using ten randomly selected flies each day. Flies were infected with Mr727 (b), Mr1046 (c), or Mr1-16 (d) at a dose of 2.5 × 10⁴ spores/mL and sampled at the moribund stage.

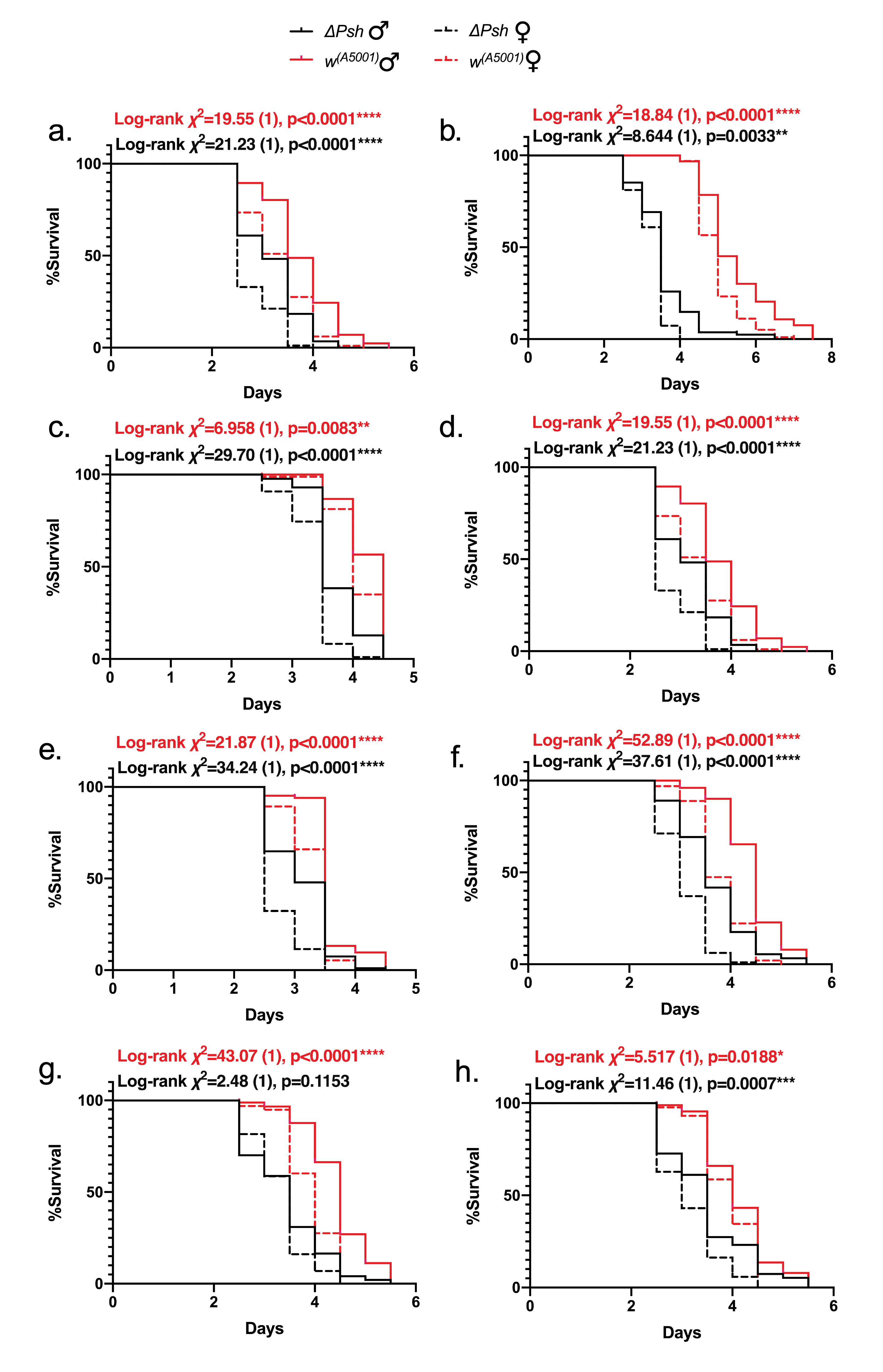

**S10 Fig. Kaplan-Meier survival curves of fly lines disrupted in Δ*Psh* and their isogenic WT background (*w^(A5001)^*) following infections with *M. robertsii* strains.** Log-rank tests were used to determine whether there were significant differences between males and females in each fly line. Δ*Psh* is represented in black and *w^(A5001)^* in red. The fungal lines are a) Mr727, b) Mr1046, c) Mr1120, d) Mr2547, e) Mr1878, f) Mr1-16, g) Mr2575 and h) Mr23.

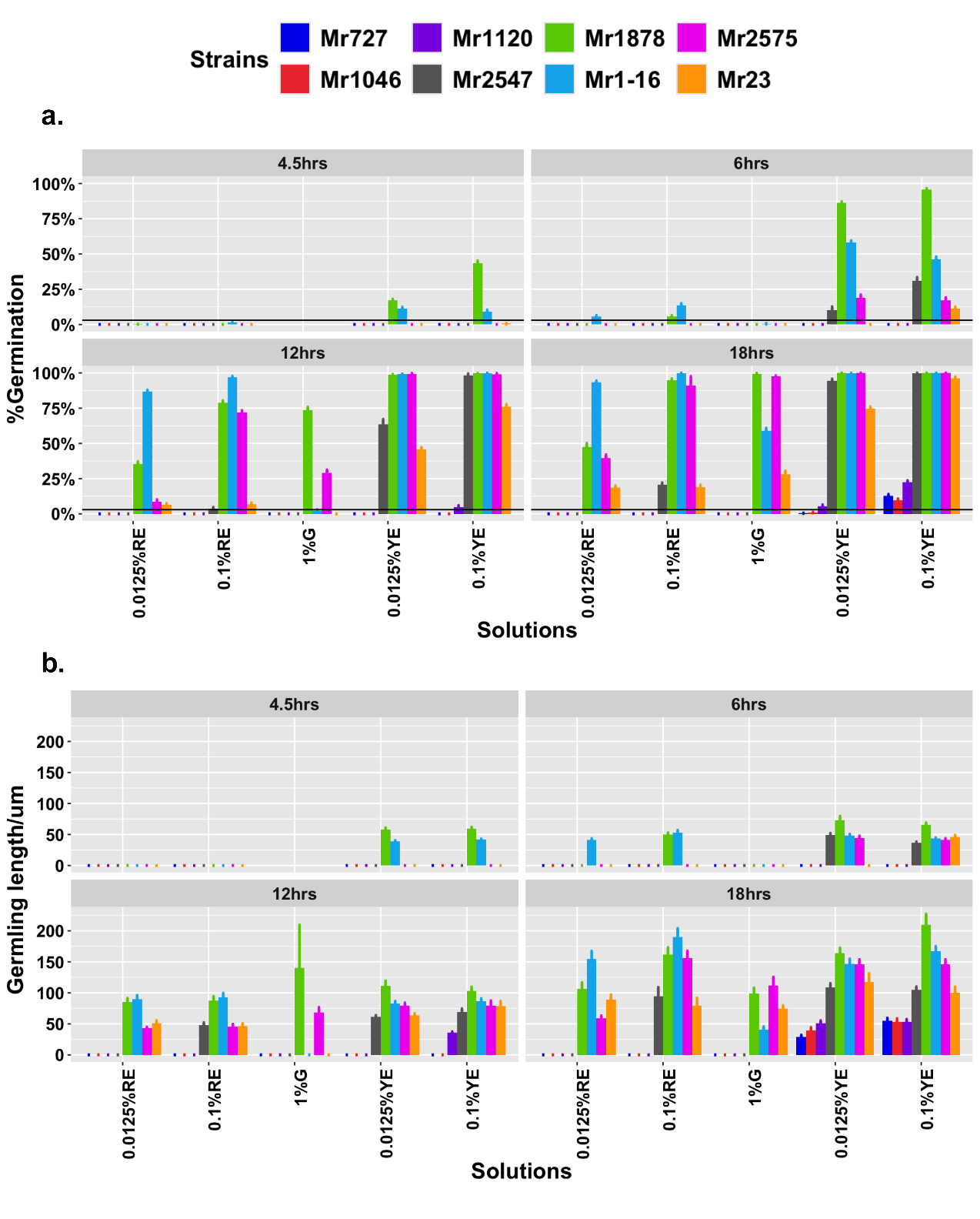

**S11 Figure. Germination of the eight *M. robertsii* strains was assessed across various nutrient sources at intervals of 4.5-, 6-, 12-, and 18-hours post-inoculation.**

RE: root exudate; YE: yeast extracts; G: glucose. Germination rates (mean ± SE) were calculated based on 15 replicates, and greater than 3% germination (black horizontal line) was counted as germinated.

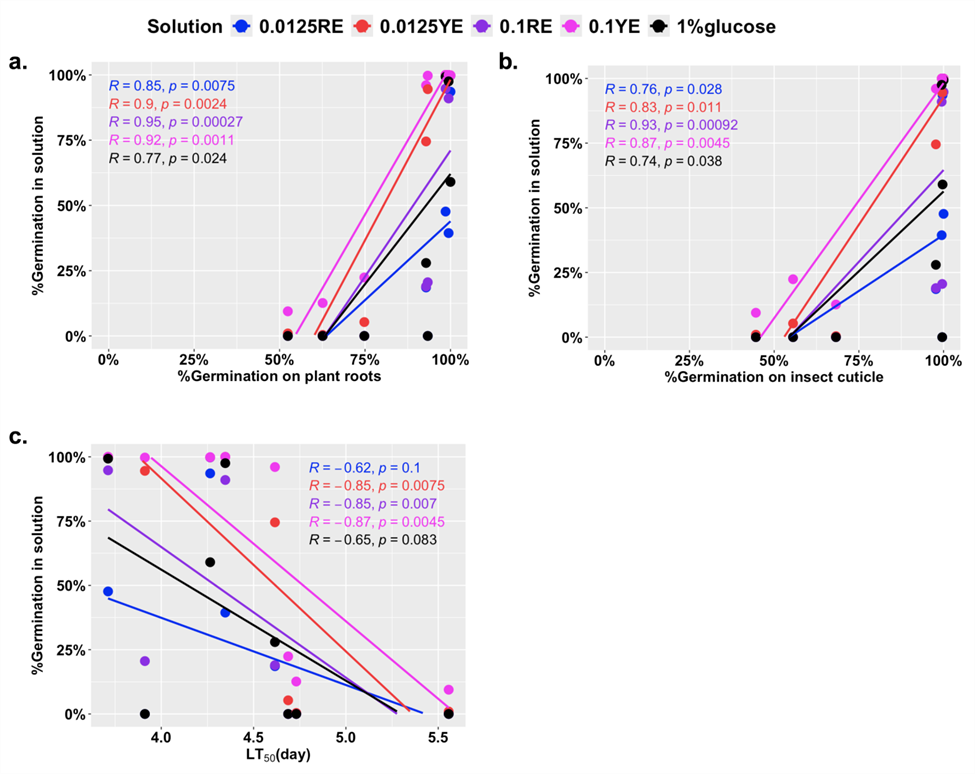

**S12 Figure. Correlations between the percent germination of *M. robertsii* strains 18 hours post-inoculation in various nutrients, plant roots, and insect cuticle, and with their virulence to insects.** All correlations were performed using Spearman’s rank test. a) Germination of *M. robertsii* strains in root exudate (RE), yeast extract (YE), and glucose was significantly correlated with their germination on plant roots. b) and insect cuticle, c) Male Drs-GFP LT_50_ values negatively correlated with *M. robertsii* germination in various nutrients, although some correlations fell short of significance.

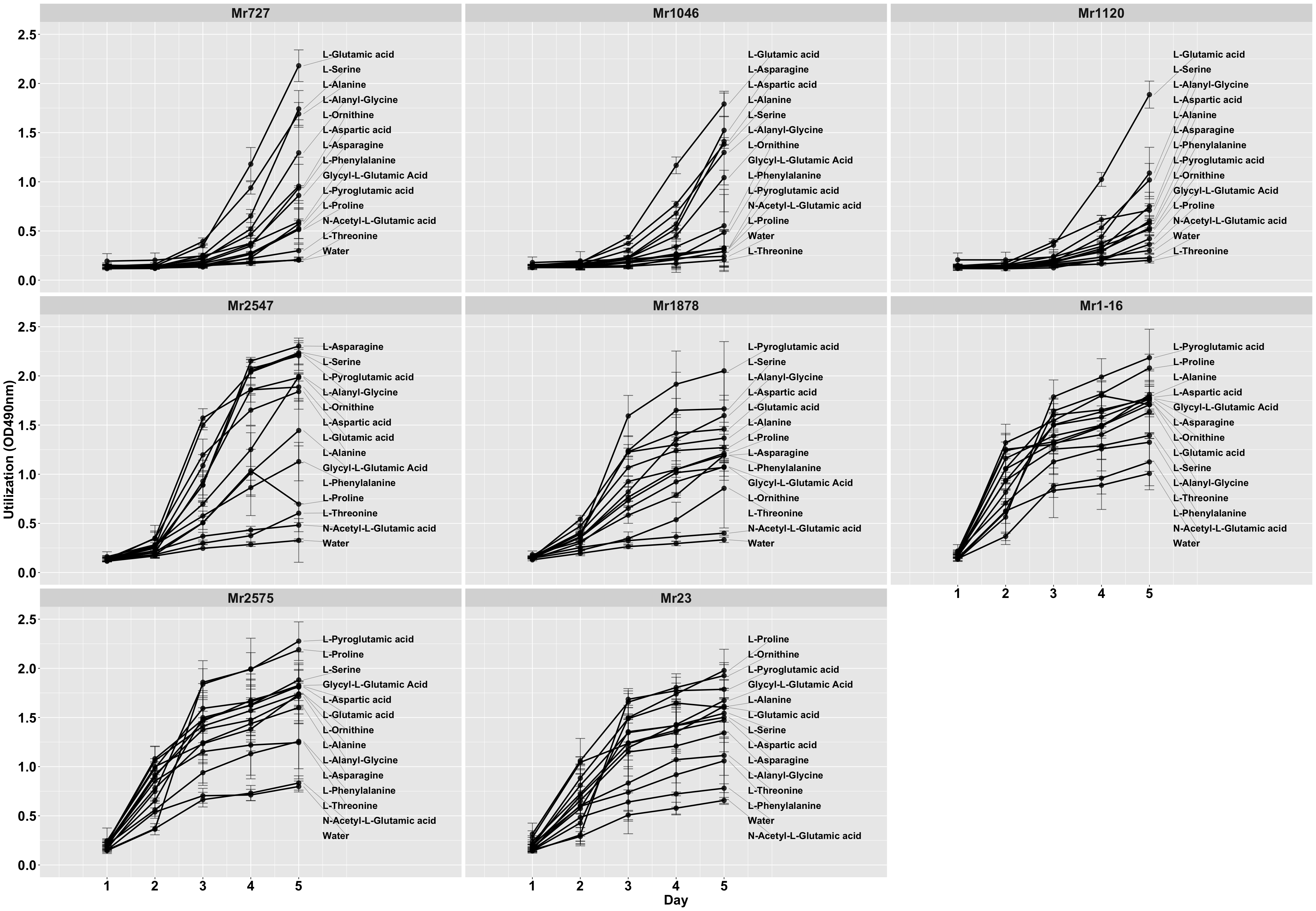

**S13 Figure. Time-course of amino acid utilization by M. robertsii strains.**
Substrate utilization was monitored over five days by measuring absorbance at 490 nm. Each data point represents the mean ± standard deviation from three biological replicates for individual amino acid substrates. On the right, amino acid substrates were ranked based on the strains utilization on day five, from highest (top) to lowest (bottom). Water was included as a control.

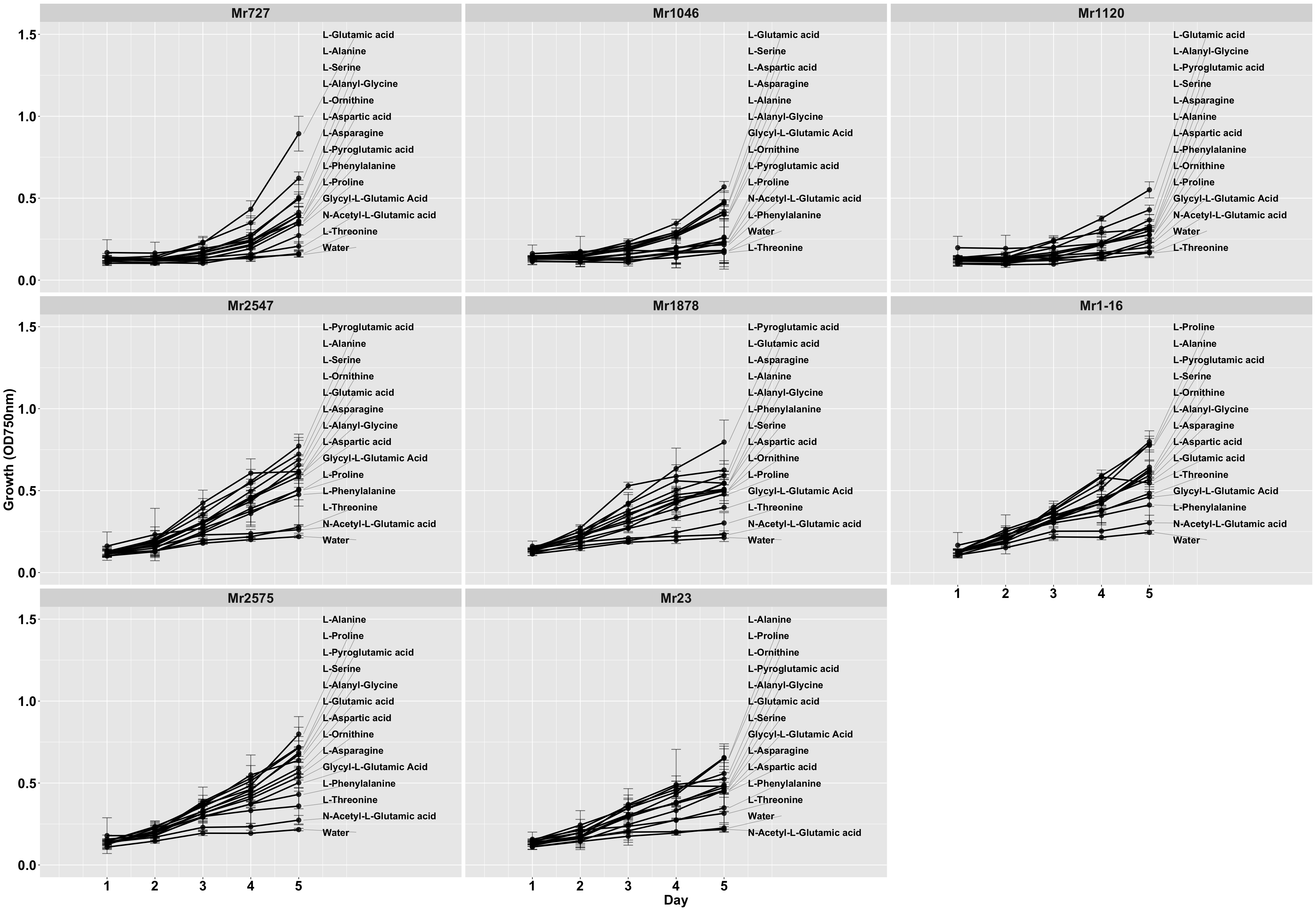

**S14 Figure. Time-course of amino acid growth by M. robertsii strains.**
Substrate growth was monitored over five days by measuring absorbance at 750 nm. Each data point represents the mean ± standard deviation from three biological replicates for individual amino acid. On the right, amino acid substrates were ranked based on the strains growth on day five, from highest (top) to lowest (bottom). Water was included as a control.

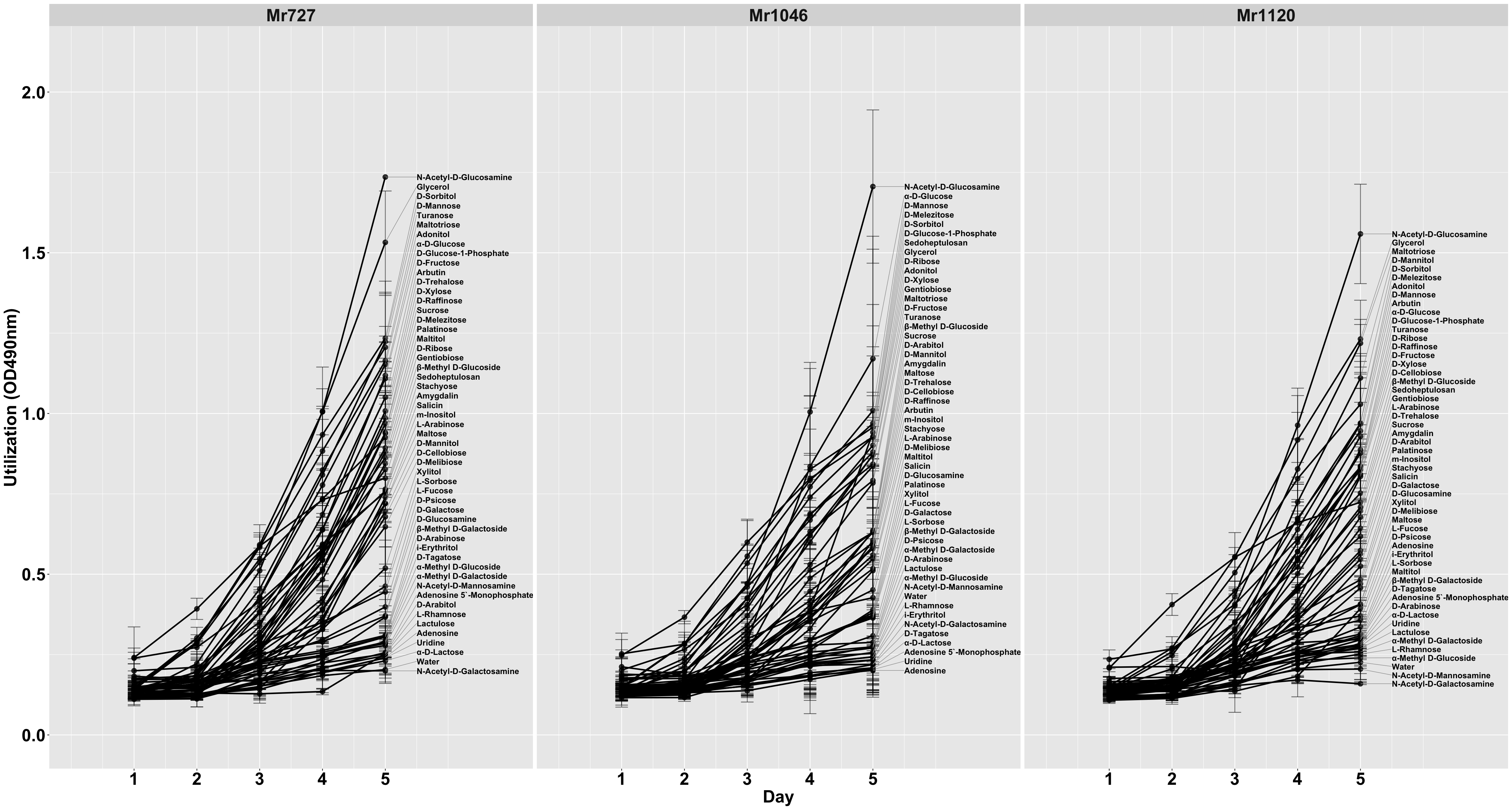

**S15 Figure. Time-course of the utilization of carbohydrate substrates by Mr727, Mr1046 and Mr1120*.***
Substrate utilization was monitored over five days by measuring absorbance at 490 nm. Each data point represents the mean ± standard deviation from three biological replicates for individual **carbohydrate**. On the right, **carbohydrate** substrates were ranked based on the strains utilization on day five, from highest (top) to lowest (bottom). Water was included as a control.

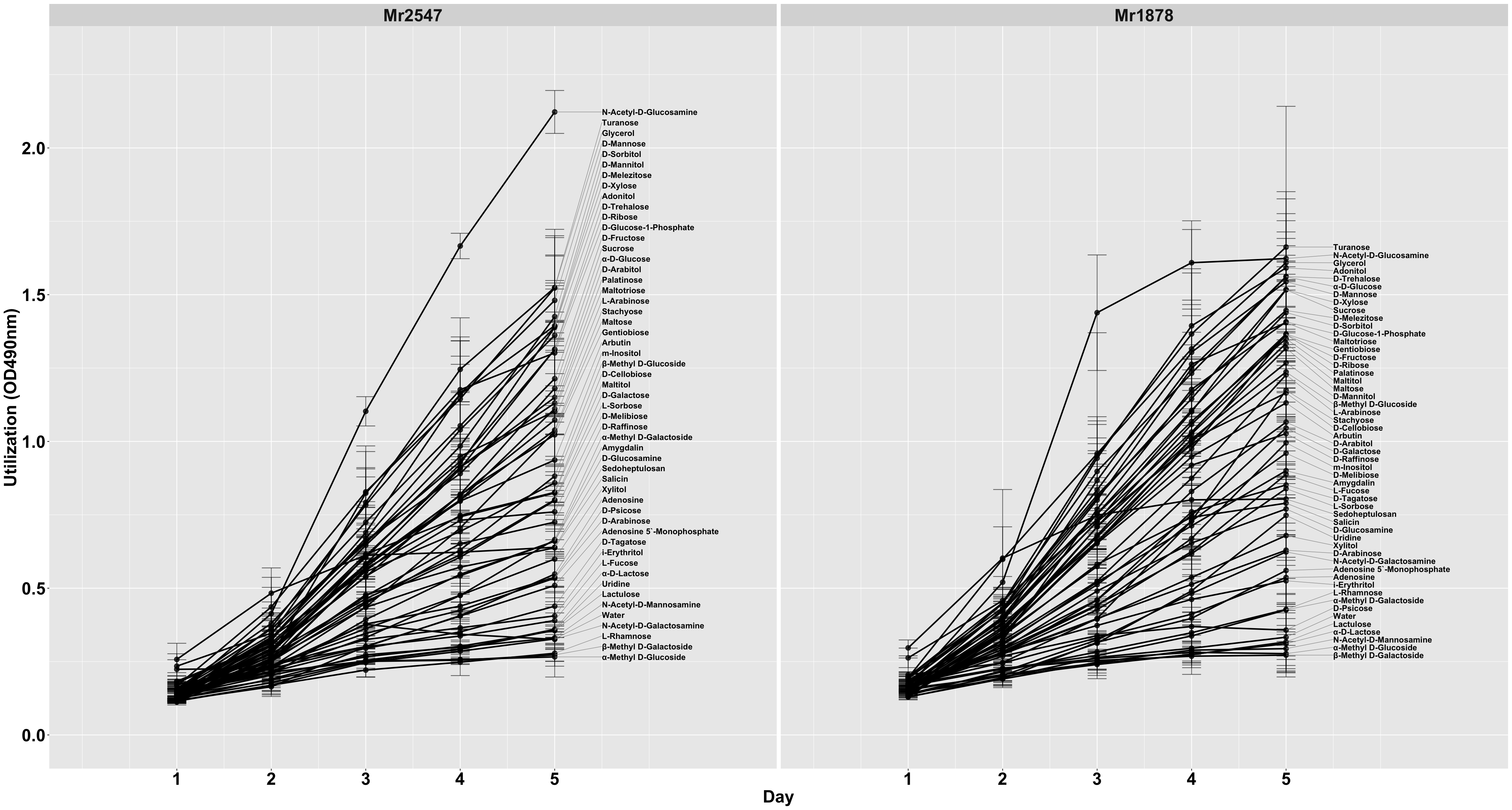

**S16 Figure. Time-course of the utilization of carbohydrate substrates by Mr2547 and Mr1878 in FF MicroPlates.**
Substrate utilization was monitored over five days by measuring absorbance at 490 nm. Each data point represents the mean ± standard deviation from three biological replicates for individual **carbohydrate**. On the right, **carbohydrate** substrates were ranked based on the strains utilization on day five, from highest (top) to lowest (bottom). Water was included as a control.

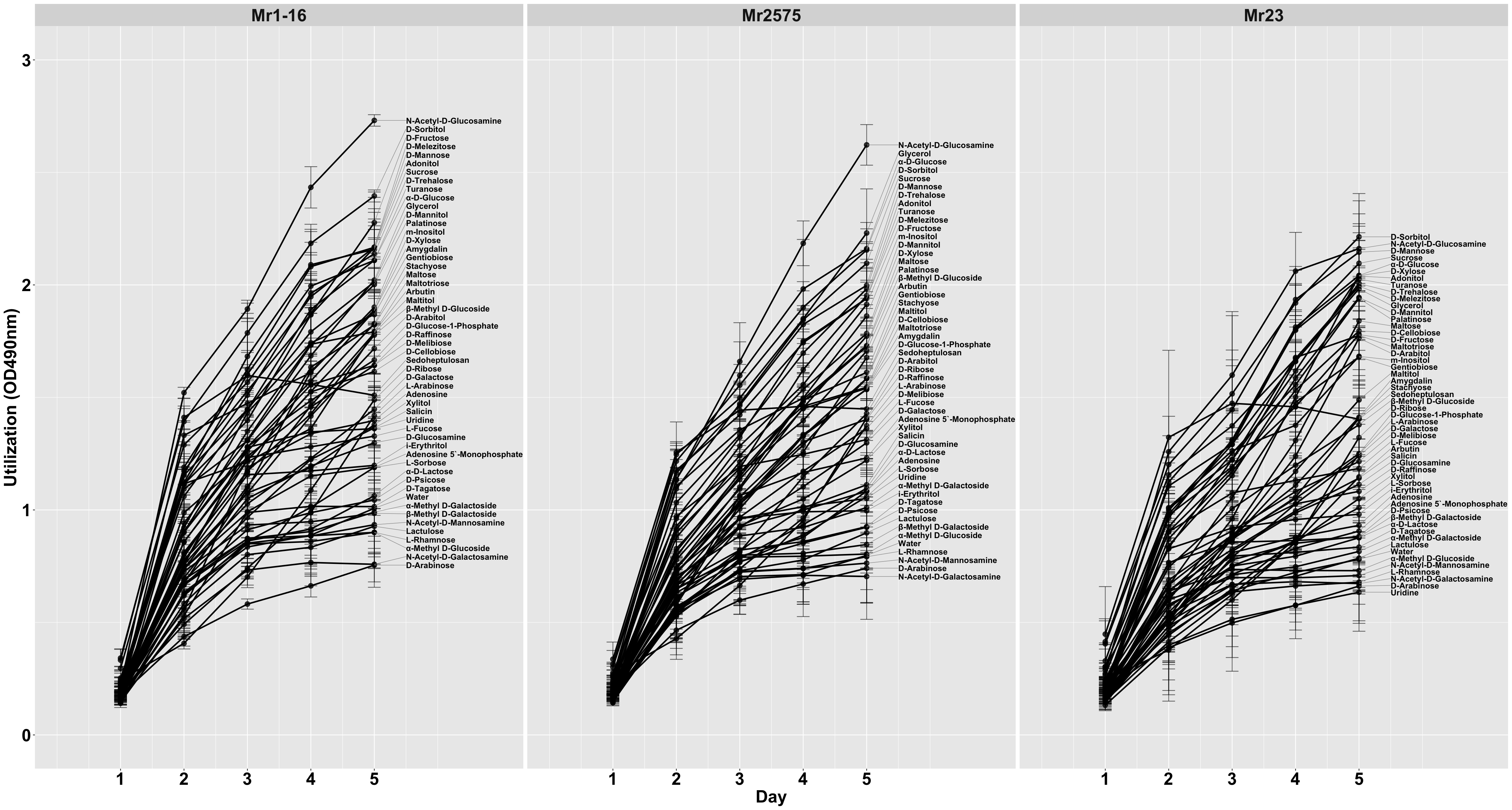

**S17 Figure. Time-course of the utilization of carbohydrate substrates by Mr1-16, Mr2575 and Mr23 in FF MicroPlates.**
Substrate utilization was monitored over five days by measuring absorbance at 490 nm. Each data point represents the mean ± standard deviation from three biological replicates for individual **carbohydrate**. On the right, **carbohydrate** substrates were ranked based on the strains utilization on day five, from highest (top) to lowest (bottom). Water was included as a control.

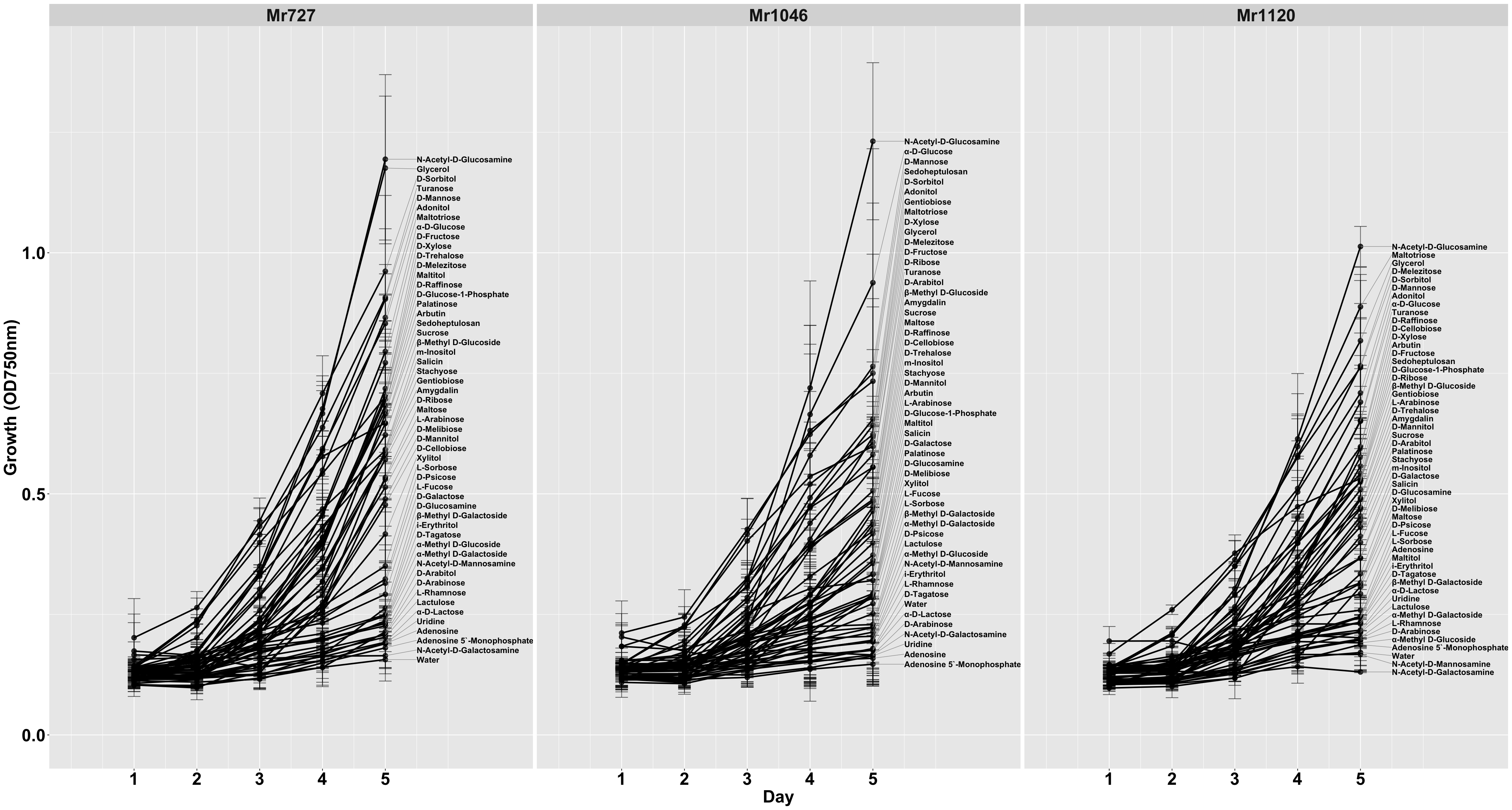

**S18 Figure. Time-course of mycelial growth by Mr727, Mr1046 and Mr1120 on carbohydrate substrates in FF MicroPlates.**
Substrate growth was monitored over five days by measuring absorbance at 750 nm. Each data point represents the mean ± standard deviation from three biological replicates for individual **carbohydrate**. On the right, **carbohydrate** substrates were ranked based on the strains growth on day five, from highest (top) to lowest (bottom). Water was included as a control.

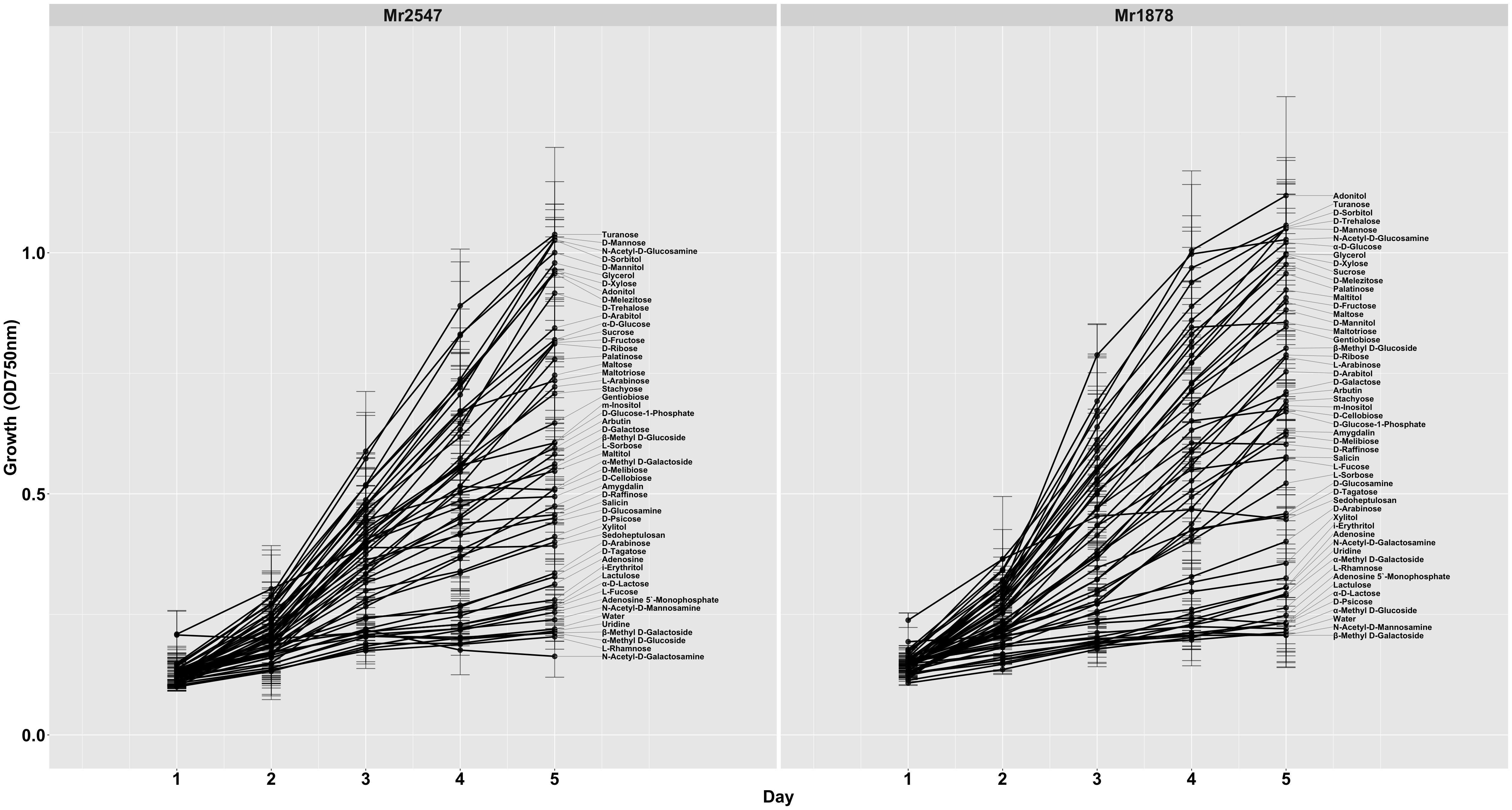

**S19 Figure. Time-course of mycelial growth by Mr2547 and Mr1878 on carbohydrate substrates in FF MicroPlates.**
Substrate growth was monitored over five days by measuring absorbance at 750 nm. Each data point represents the mean ± standard deviation from three biological replicates for individual **carbohydrate**. On the right, **carbohydrate** substrates were ranked based on each *M. robertsii* strains biomass on day five, from highest (top) to lowest (bottom). Water was included as a control.

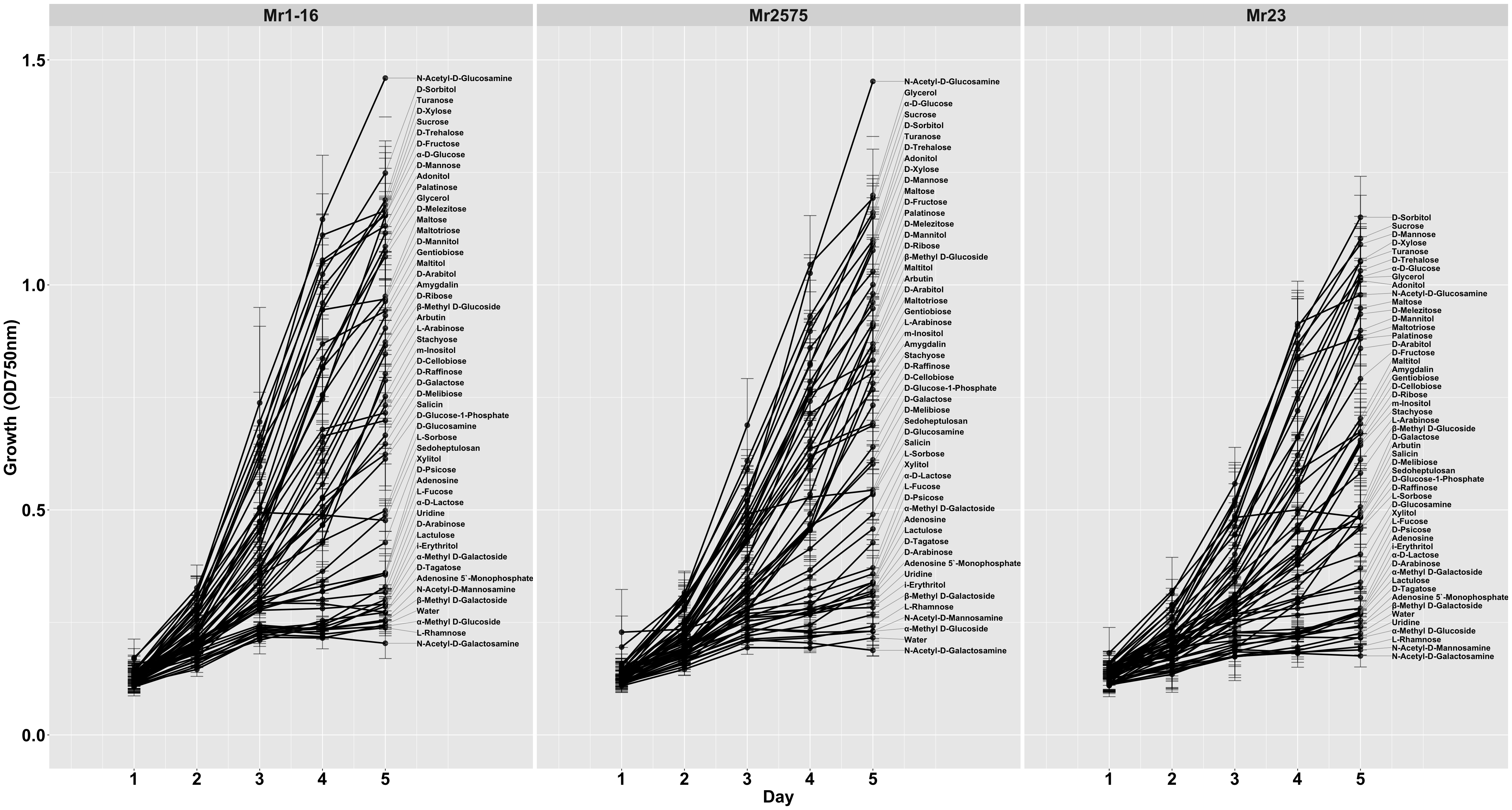

**S20 Figure. Time-course of mycelial growth by Mr1-16, Mr2575 and Mr23 on carbohydrate substrates in FF MicroPlates.**
Substrate growth was monitored over five days by measuring absorbance at 750 nm. Each data point represents the mean ± standard deviation from three biological replicates for individual amino acid carbon substrates. On the right, amino acid substrates were ranked based on each *M. robertsii* strains growth on day five, from highest (top) to lowest (bottom). Water was included as a control.

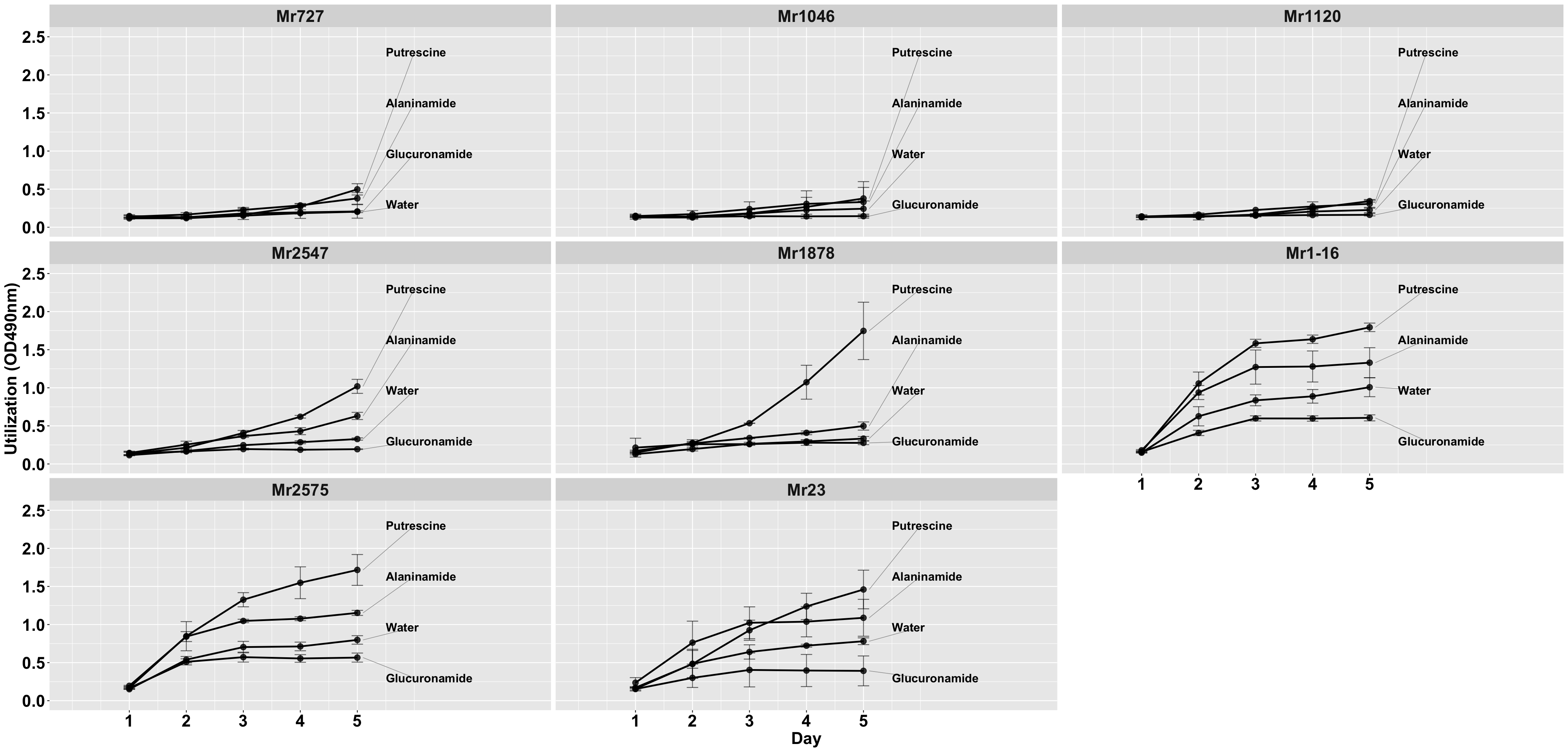

**S21 Figure. Time-course of amine and amide utilization by M. robertsii strains.**
Substrate utilization was monitored over five days by measuring absorbance at 490 nm. Each data point represents the mean ± standard deviation from three biological replicates for individual **amine and amide**. On the right, **amine and amide** substrates were ranked based on the strains utilization on day five, from highest (top) to lowest (bottom). Water was included as a control.

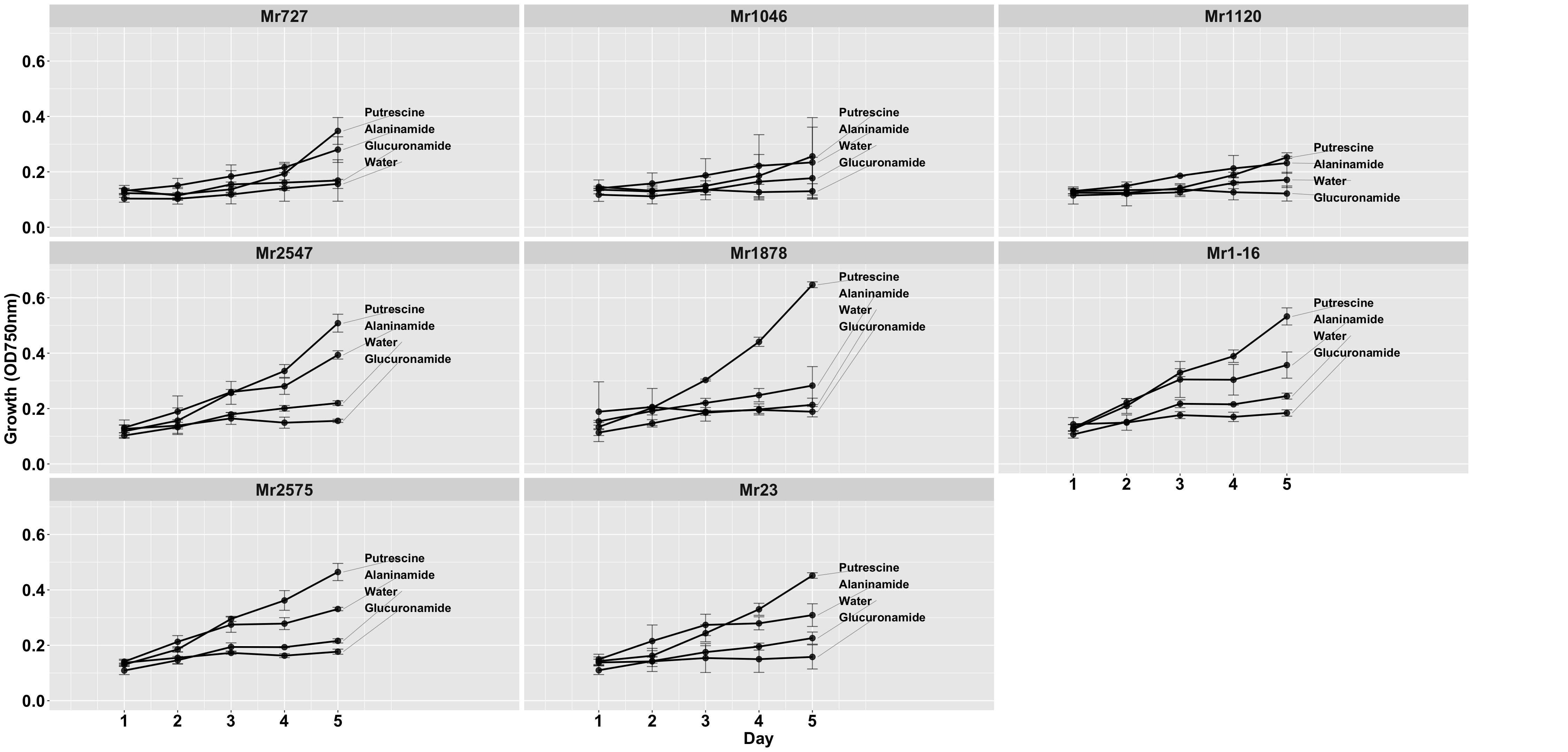

**S22 Figure. Time-course of amine and amide growth by M. robertsii strains.**
Substrate growth was monitored over five days by measuring absorbance at 750 nm. Each data point represents the mean ± standard deviation from three biological replicates for individual **amine and amide**. On the right, **amine and amide** substrates were ranked based on the strains growth on day five, from highest (top) to lowest (bottom). Water was included as a control.

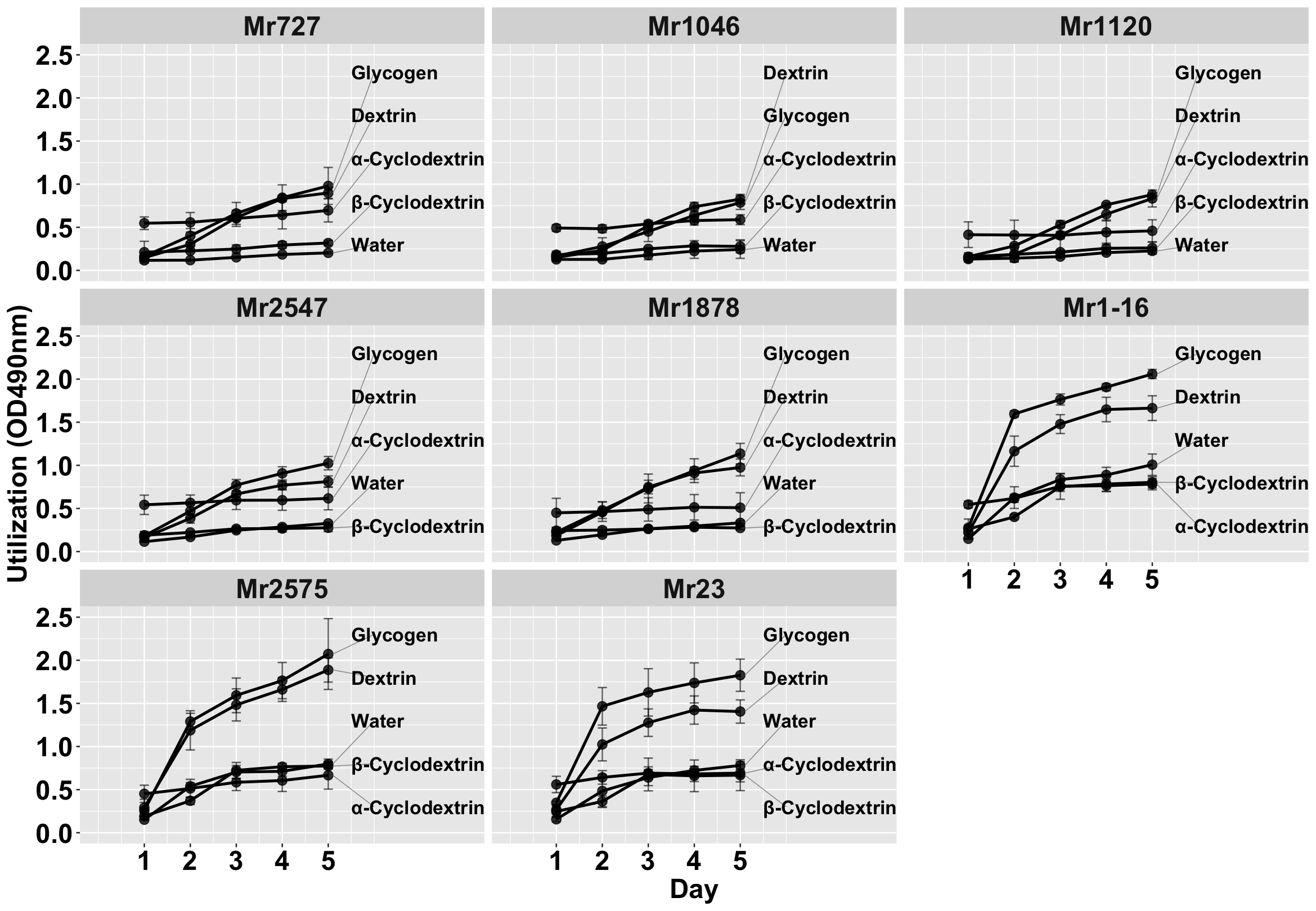

**S23 Figure. Time-course of polymer utilization by M. robertsii strains.**
Substrate utilization was monitored over five days by measuring absorbance at 490 nm. Each data point represents the mean ± standard deviation from three biological replicates for individual **polymer**. On the right, **polymer** substrates were ranked based on the strains utilization on day five, from highest (top) to lowest (bottom). Water was included as a control.

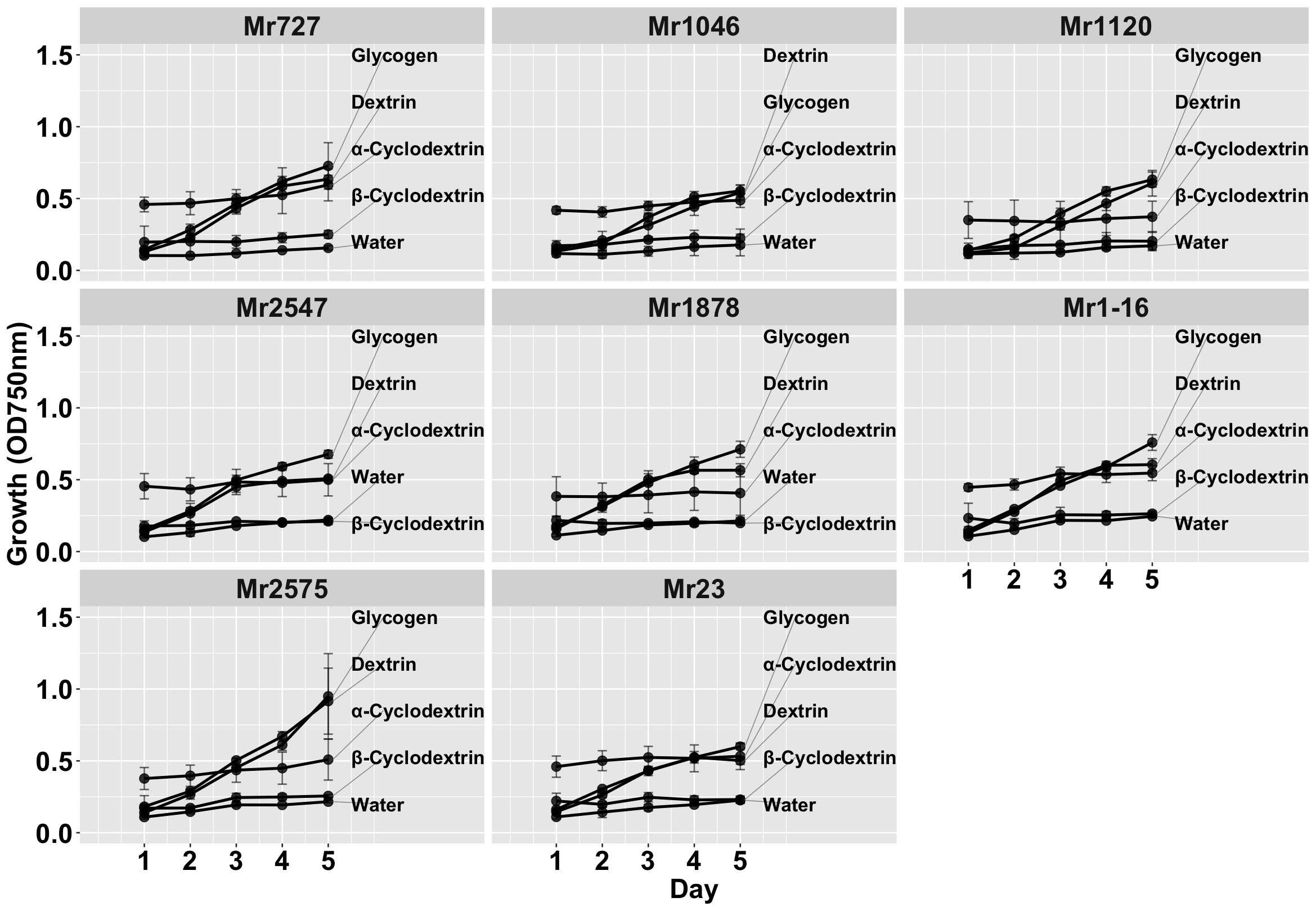

**S24 Figure. Time-course of polymers growth by M. robertsii strains.**
Substrate growth was monitored over five days by measuring absorbance at 750 nm. Each data point represents the mean ± standard deviation from three biological replicates for individual **polymer**. On the right, **polymer** substrates were ranked based on the strains growth on day five, from highest (top) to lowest (bottom). Water was included as a control.

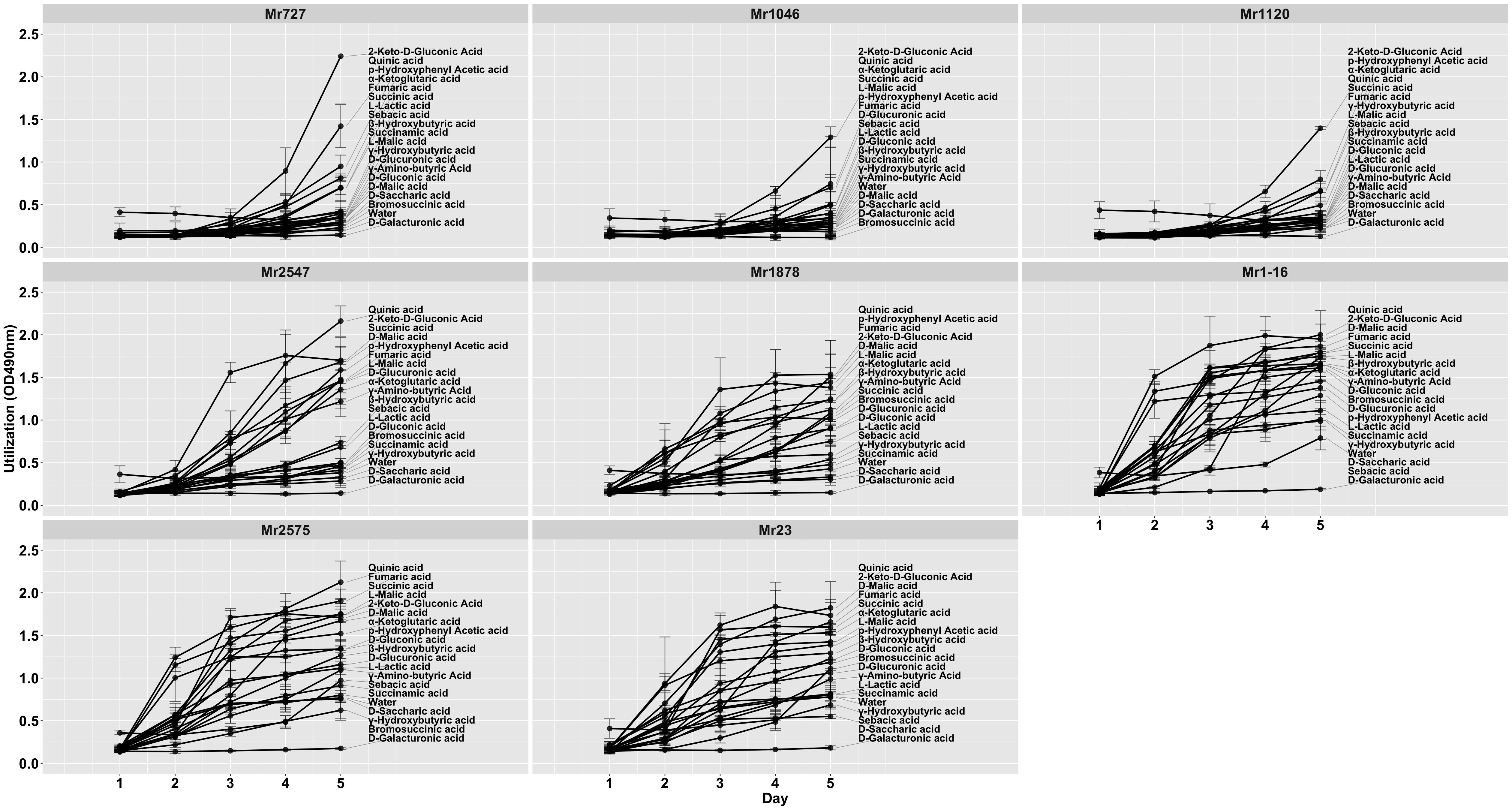

**S25 Figure. Time-course of carboxylic acid** **utilization by M. robertsii strains.**
Substrate utilization was monitored over five days by measuring absorbance at 490 nm. Each data point represents the mean ± standard deviation from three biological replicates for individual **carboxylic acid**. On the right, **carboxylic acid** substrates were ranked based on the strains utilization on day five, from highest (top) to lowest (bottom). Water was included as a control.

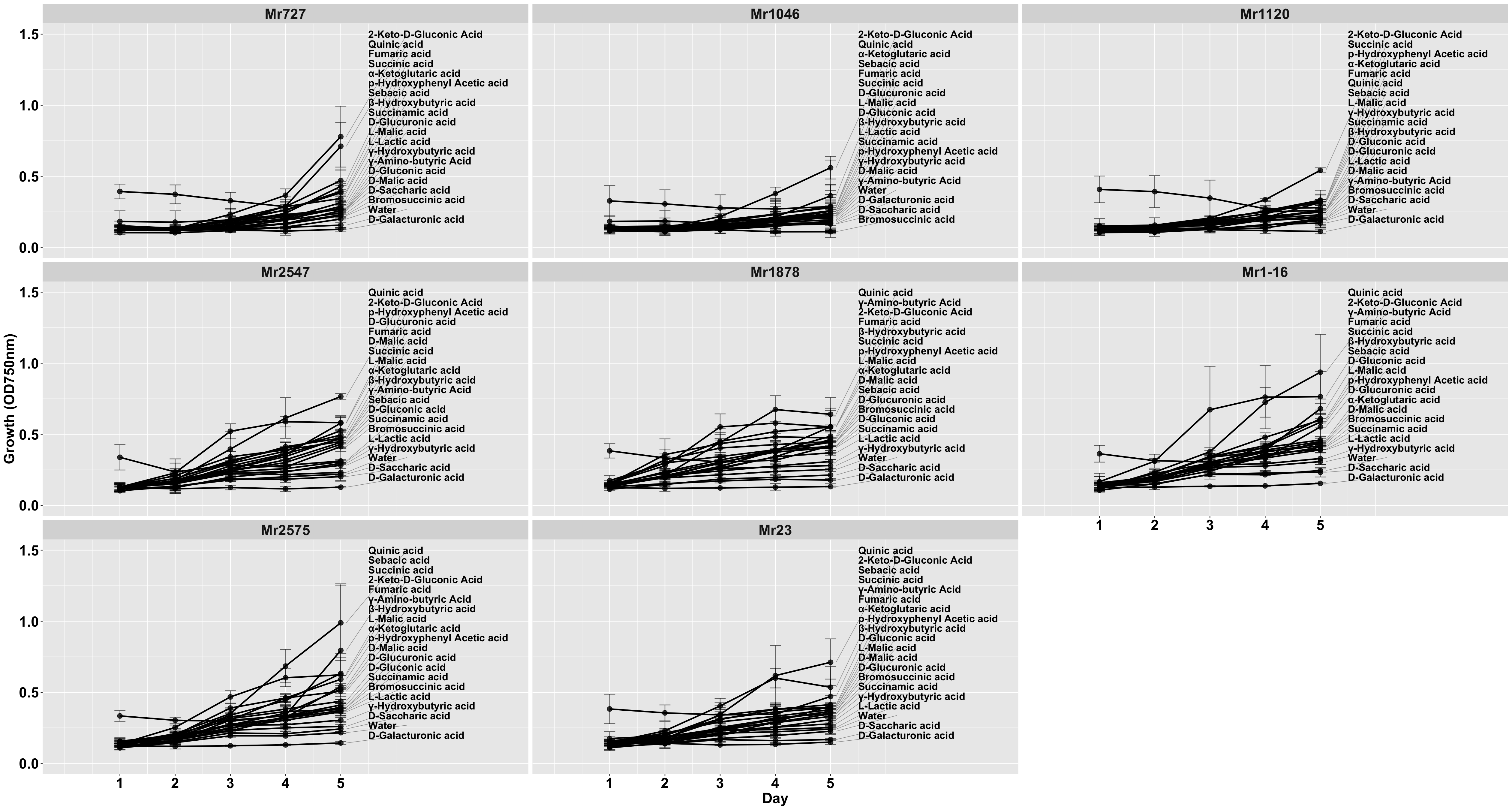

**S26 Figure. Time-course of carboxylic acid growth by M. robertsii strains.**
Substrate growth was monitored over five days by measuring absorbance at 750 nm. Each data point represents the mean ± standard deviation from three biological replicates for individual **carboxylic acid**. On the right, **carboxylic acid** substrates were ranked based on the strains growth on day five, from highest (top) to lowest (bottom). Water was included as a control.

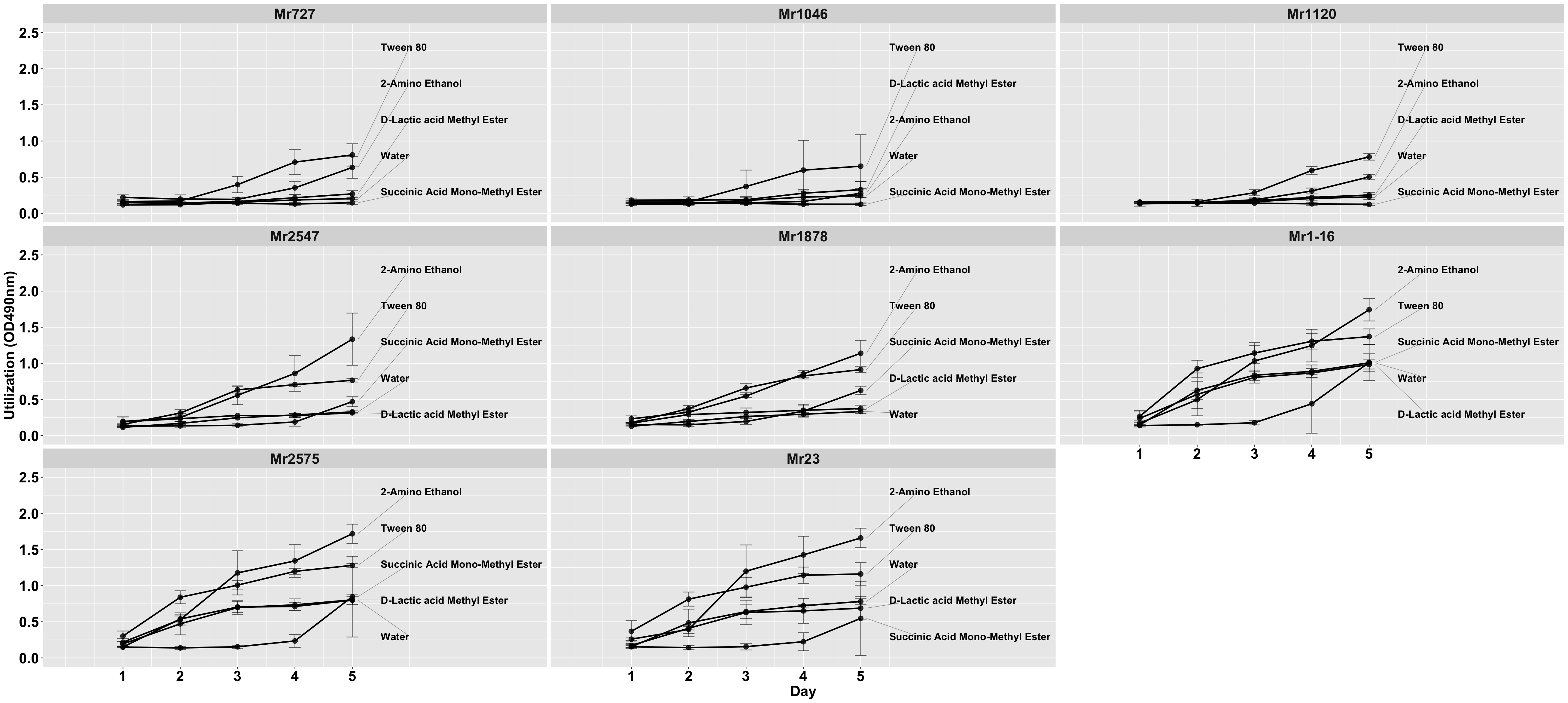

**S27 Figure. Time-course of miscellaneous utilization by M. robertsii strains.**
Substrate utilization was monitored over five days by measuring absorbance at 490 nm. Each data point represents the mean ± standard deviation from three biological replicates for individual **miscellaneous**. On the right, **miscellaneous** substrates were ranked based on the strains utilization on day five, from highest (top) to lowest (bottom). Water was included as a control.

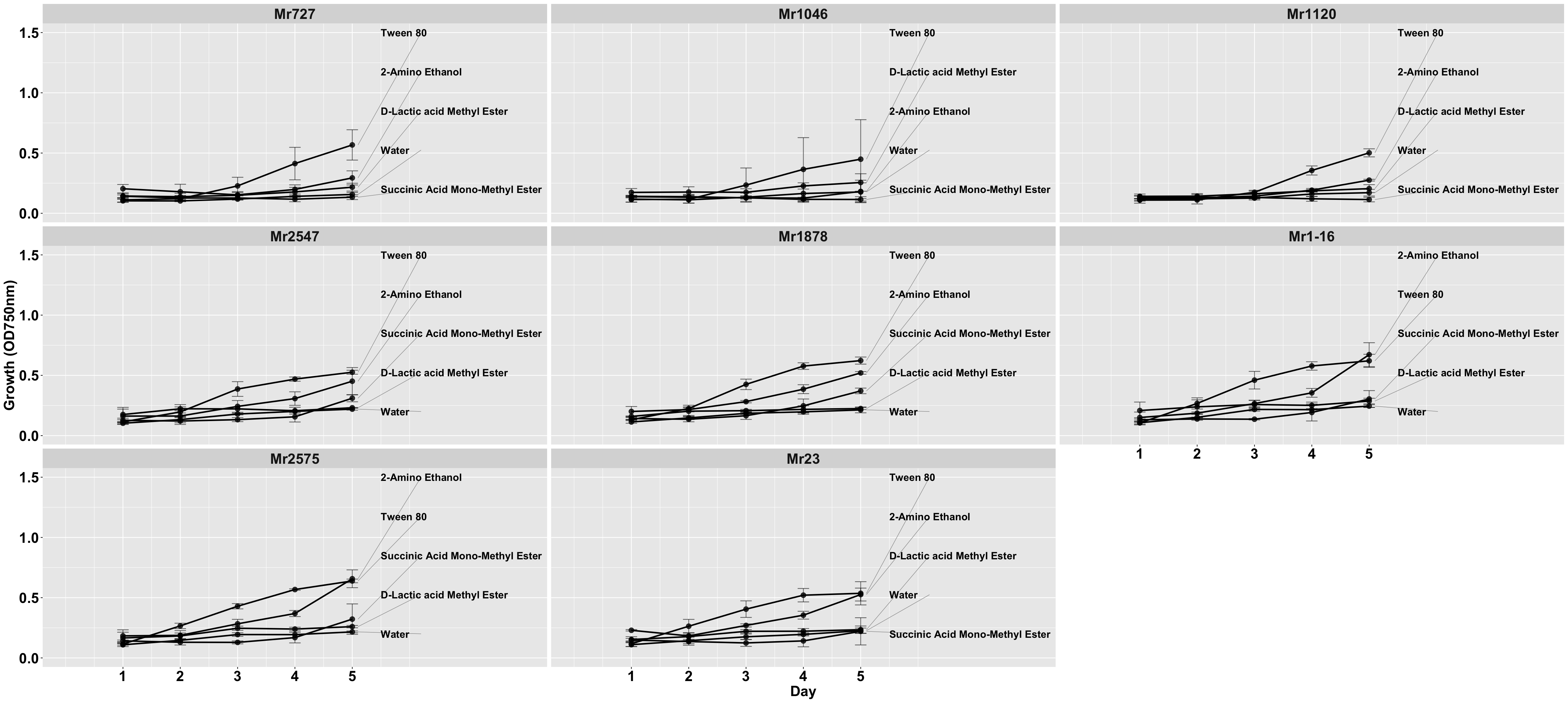

**S28 Figure. Time-course of** **miscellaneous growth by M. robertsii strains.**
Substrate growth was monitored over five days by measuring absorbance at 750 nm. Each data point represents the mean ± standard deviation from three biological replicates for individual miscellaneous. On the right, **miscellaneous** substrates were ranked based on the strains growth on day five, from highest (top) to lowest (bottom). Water was included as a control.

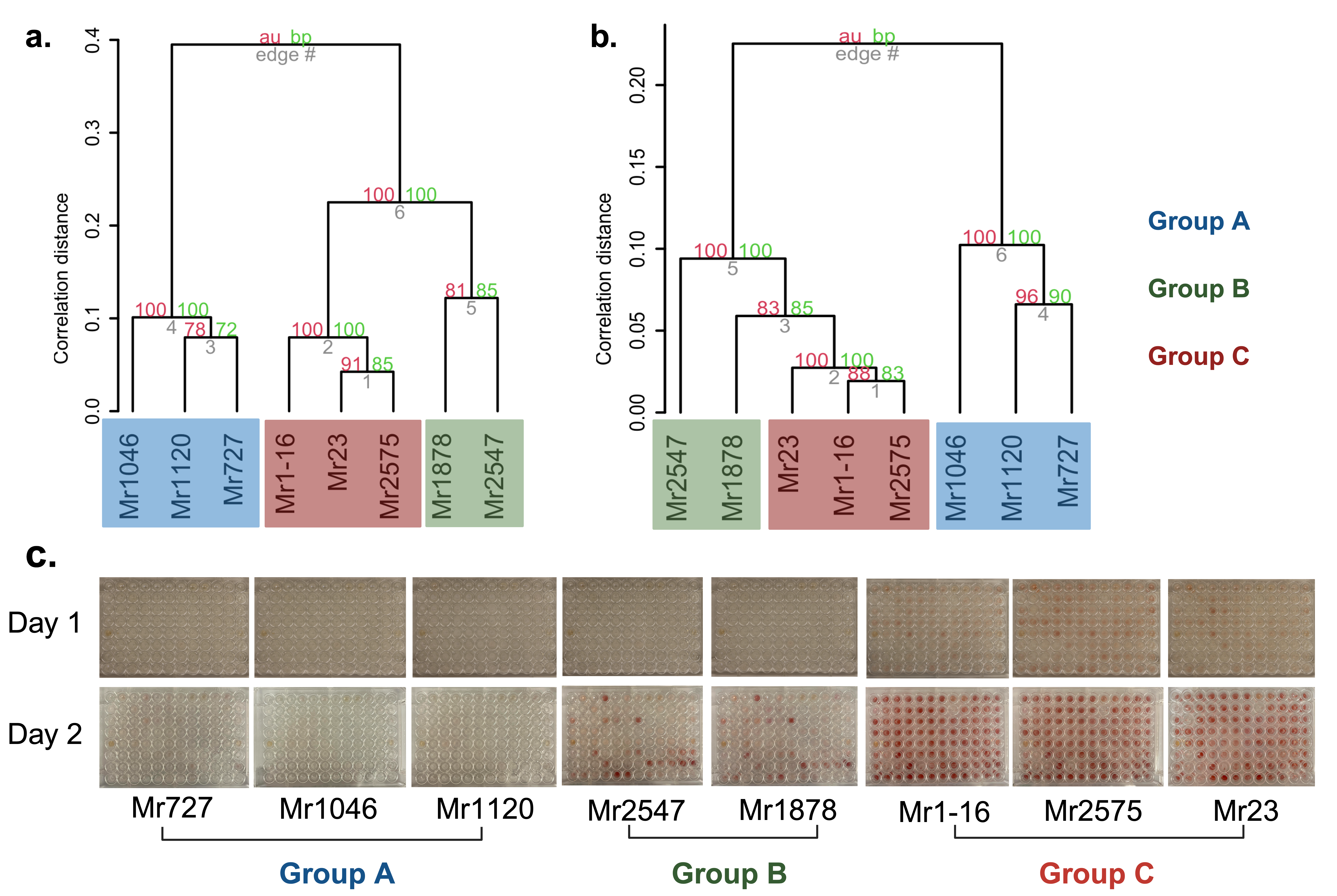

**S29 Figure. Dendrogram showing the similarity of *M. robertsii* FF MicroPlate C-source utilization (a) and growth rates (b).** The utilization rate and mycelial biomass were calculated using absorbance at 490 nm and 750 nm, respectively, on day four. The dendrogram was constructed using correlation distance with approximately unbiased (au) and bootstrap probability (bp) indicated. Based on utilization (a) and growth rates (b), *M. robertsii* strains were subdivided into groups A, B and C that aligned with early (A) and recently (B and C) diverging strains. (c) Images of FF plates from eight *M. robertsii* strains at day one and day two, showing variation in metabolic activity as indicated by color development.

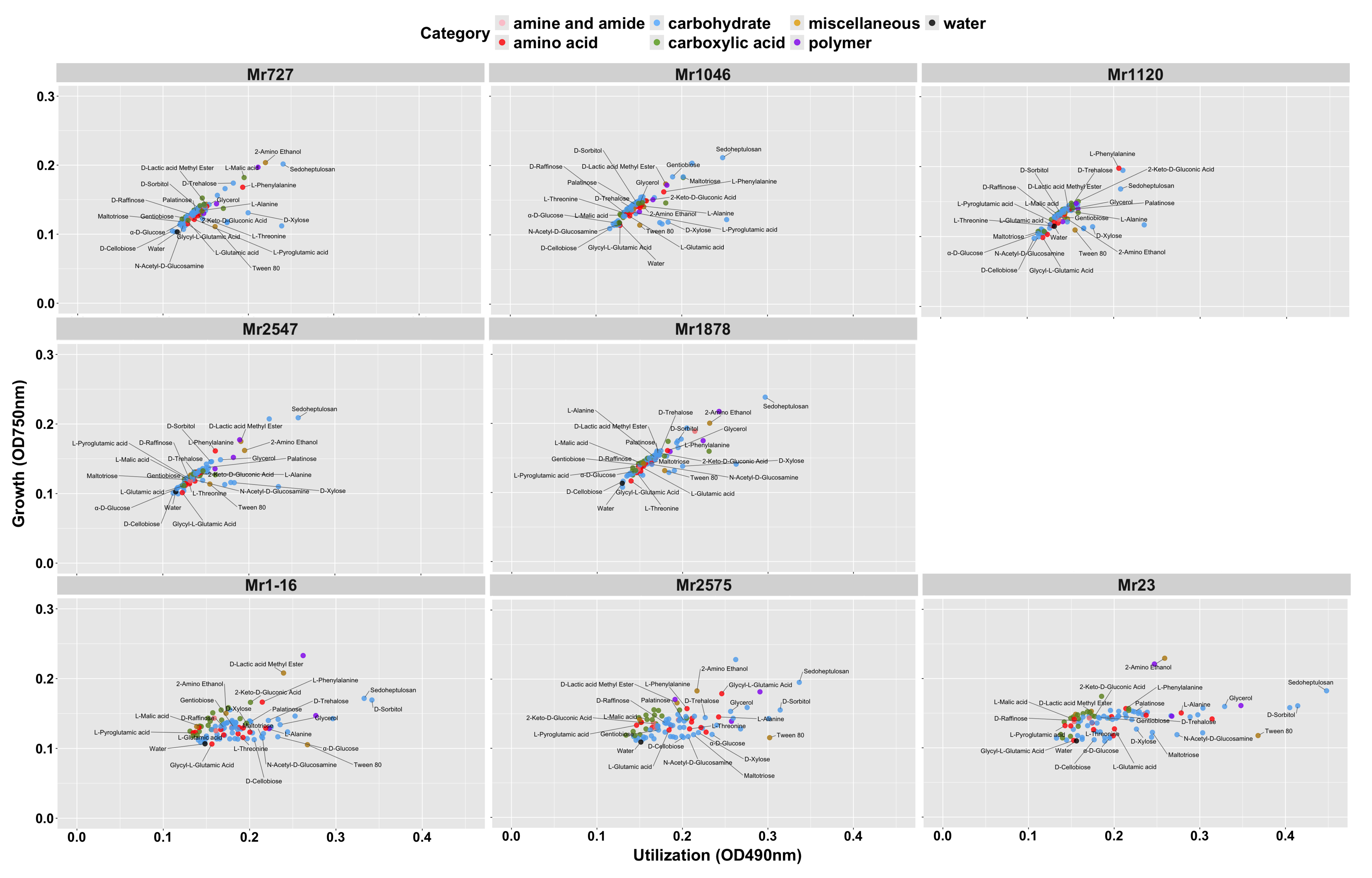

**S30 Figure. Correlation between *M. robertsii* FF MicroPlate C-source utilization (formazan dye production) and mycelial density on day one.** The utilization rate and growth rate were calculated using absorbance at 490 nm and 750 nm, respectively.

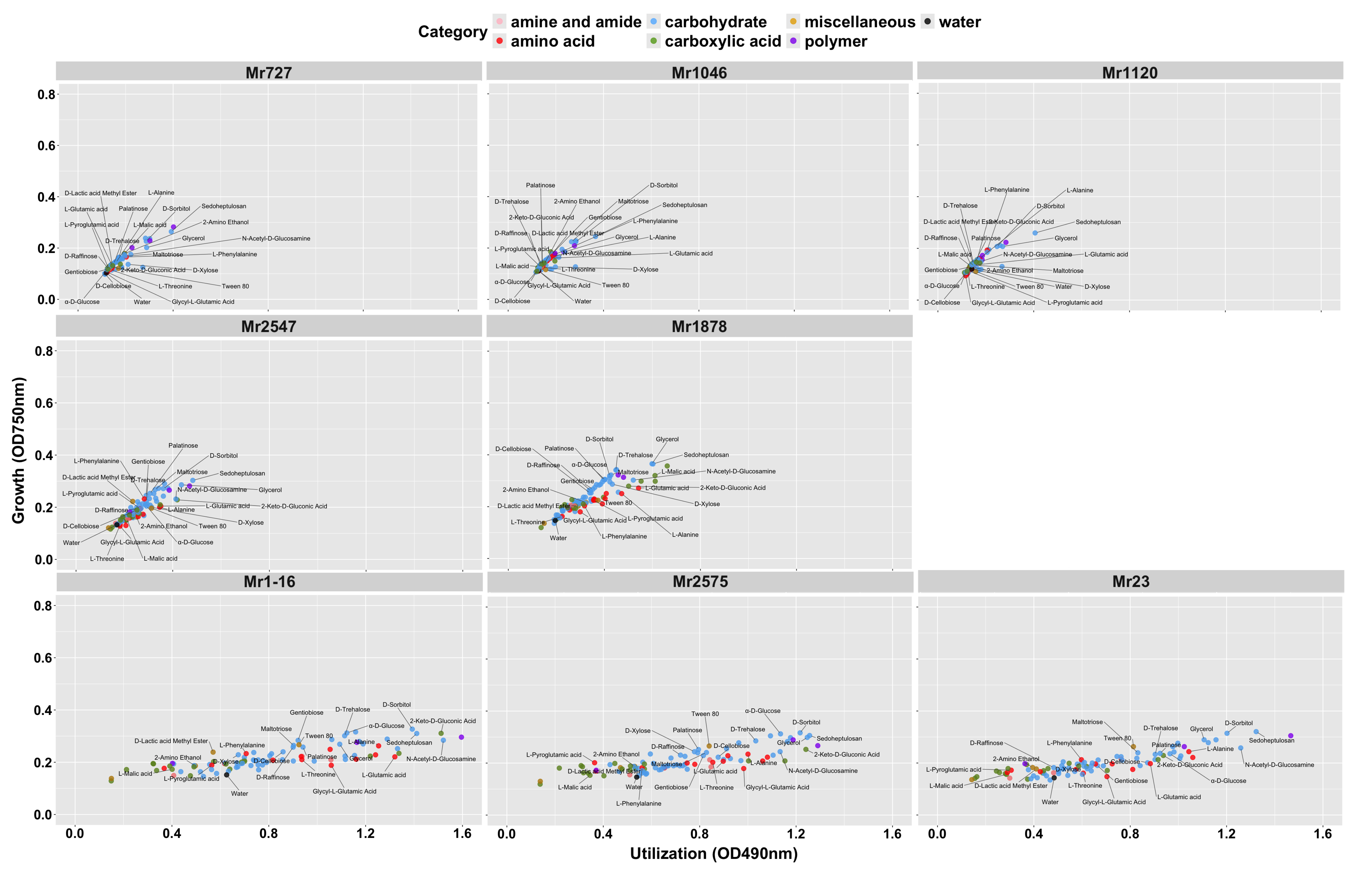
**S31 Figure. Correlation between C-source utilization and mycelial density on day two.** The utilization rate and growth rate were calculated using absorbance at 490 nm and 750 nm, respectively.

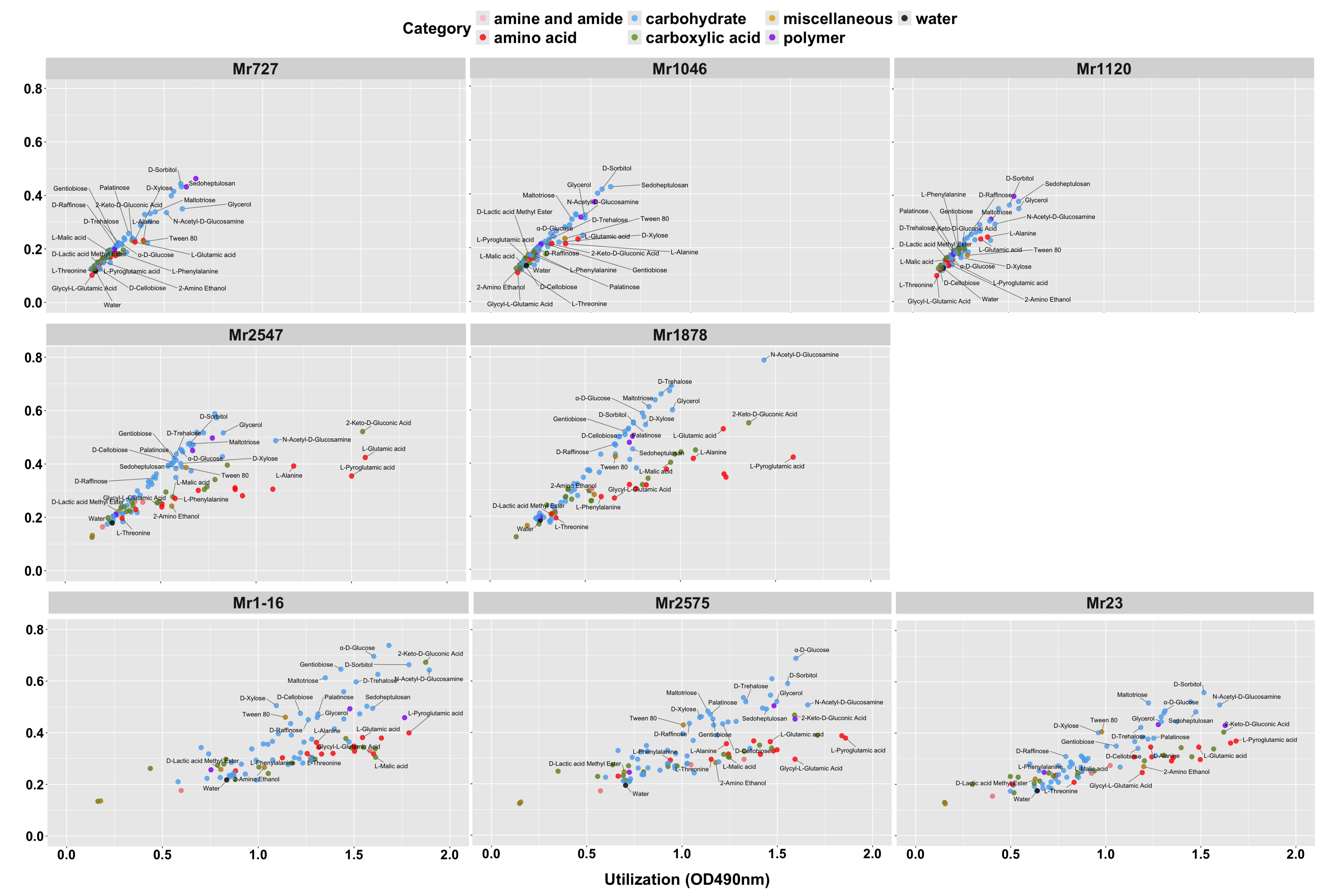

**S32 Figure. Correlation between C-source utilization and mycelial density on day three.** The utilization rate and growth rate were calculated using absorbance at 490 nm and 750 nm, respectively.

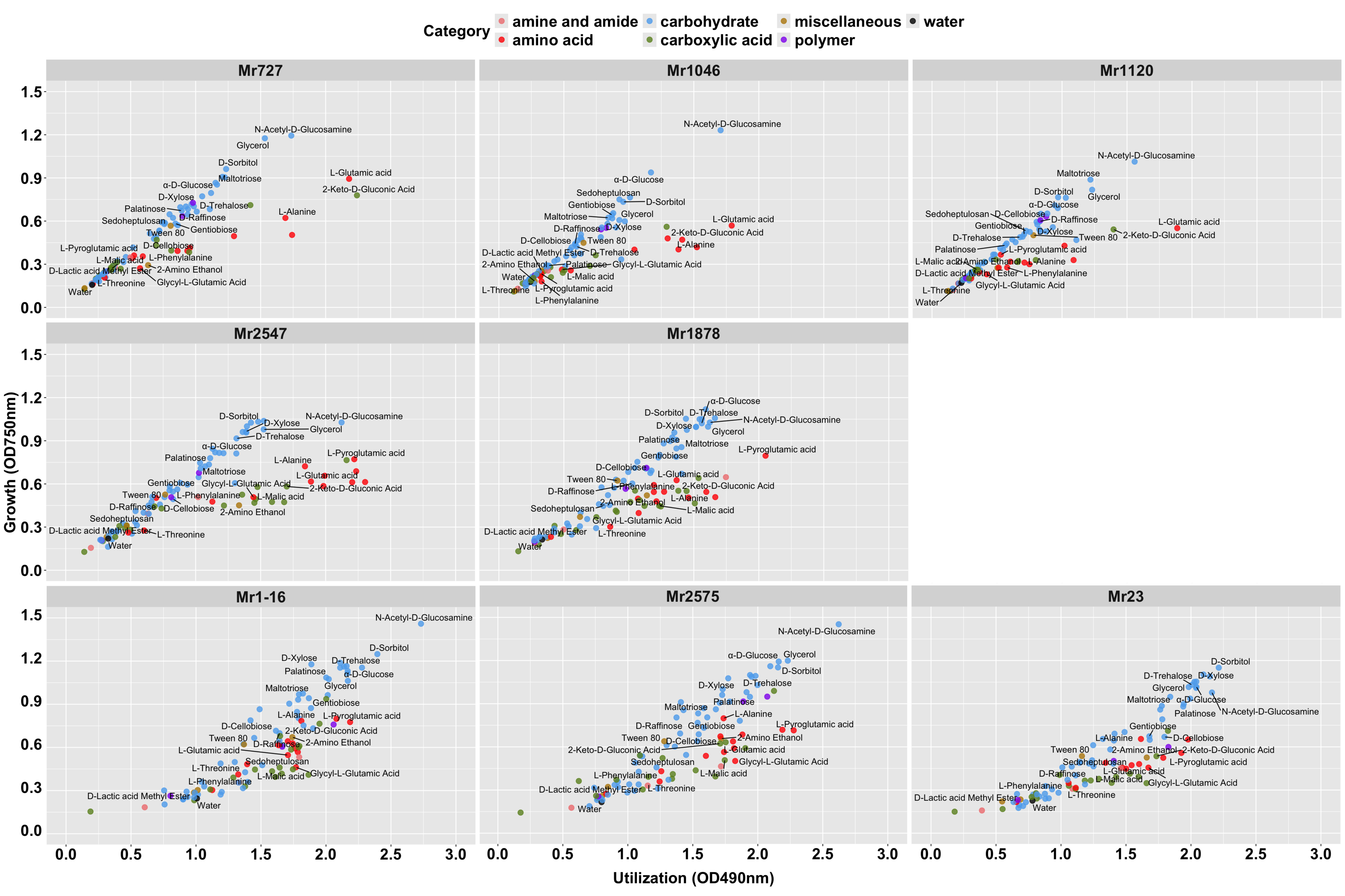

**S33 Figure. Correlation between C-source utilization and mycelial density on day five.** The utilization rate and growth rate were calculated using absorbance at 490 nm and 750 nm, respectively.

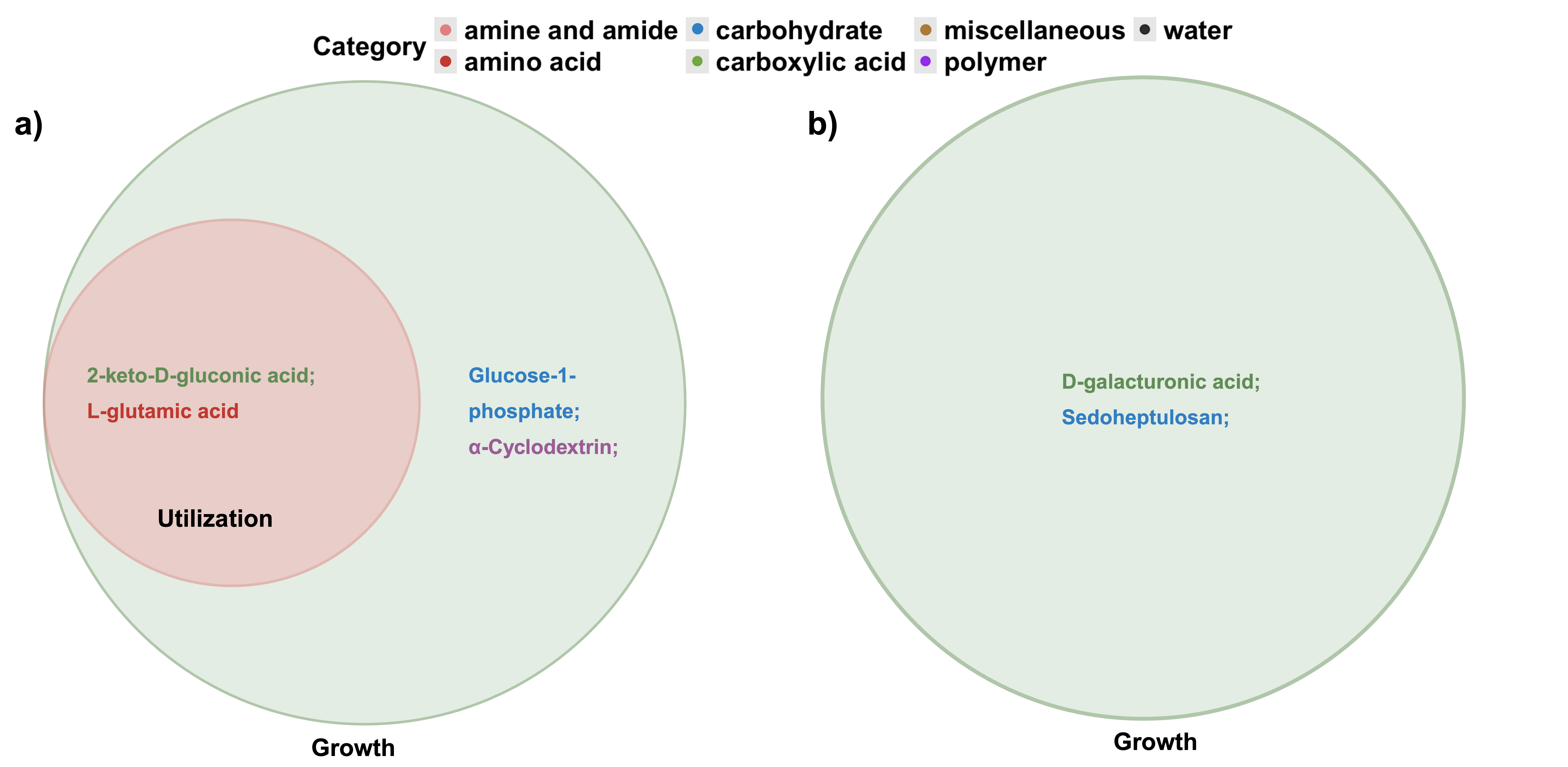

**S34 Figure. Venn diagrams showing the fastest C-substrate utilization and growth of Mr727 (a), and Mr1046(b)**. Fastest utilization is indicated in red, fastest growth in green, and overlaps represent substrates on which strains showed both. Substrates are grouped into amino acids, polymers, carbohydrates, carboxylic acids, amines and amides, miscellaneous, and water.

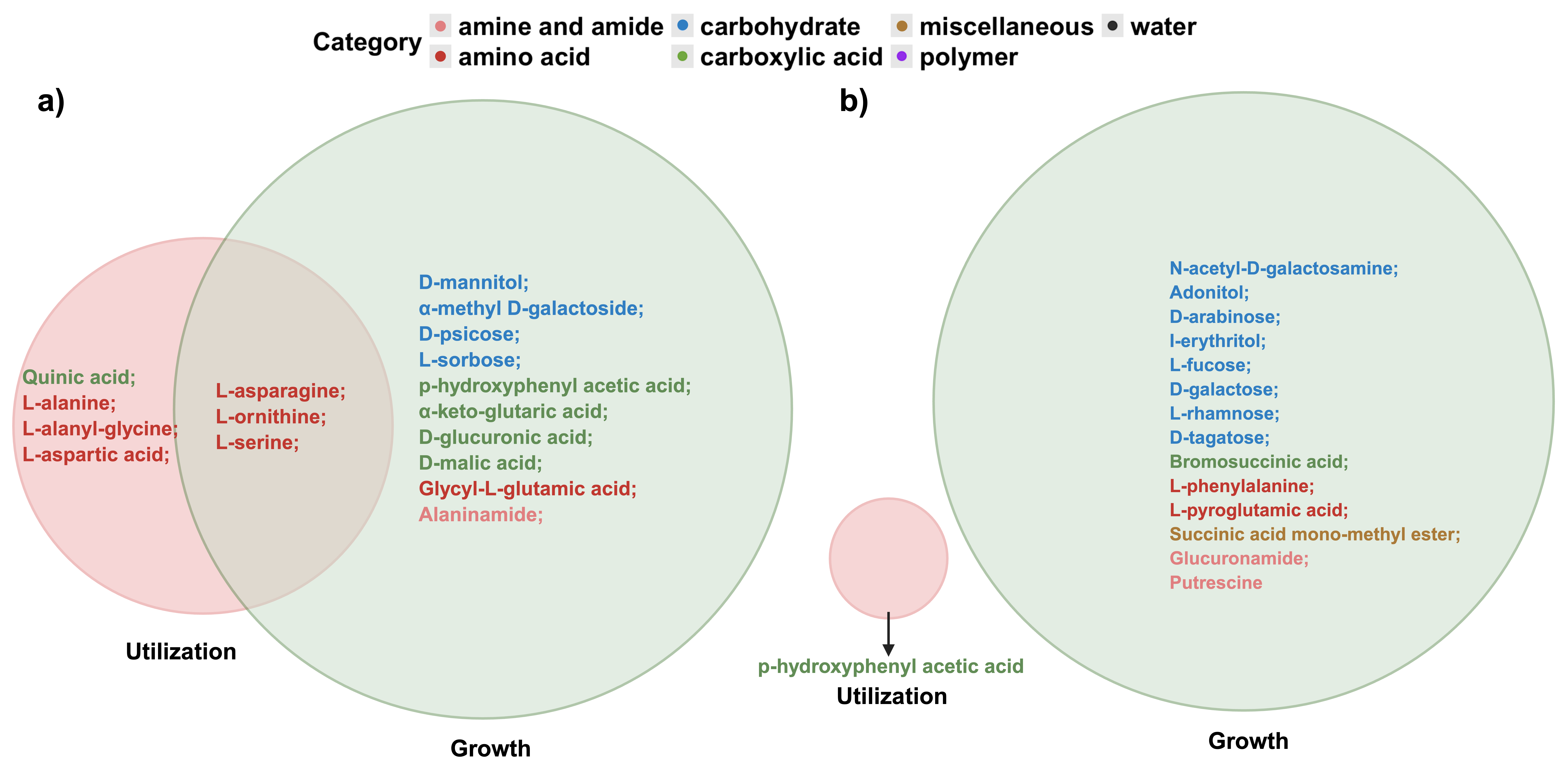

**S35 Figure. Venn diagrams showing the fastest C-substrate utilization and growth of Mr2547 (a), and Mr1878(b)**. Fastest utilization is indicated in red, fastest growth in green, and overlaps represent substrates on which strains showed both. Substrates are grouped into amino acids, polymers, carbohydrates, carboxylic acids, amines and amides, miscellaneous, and water.

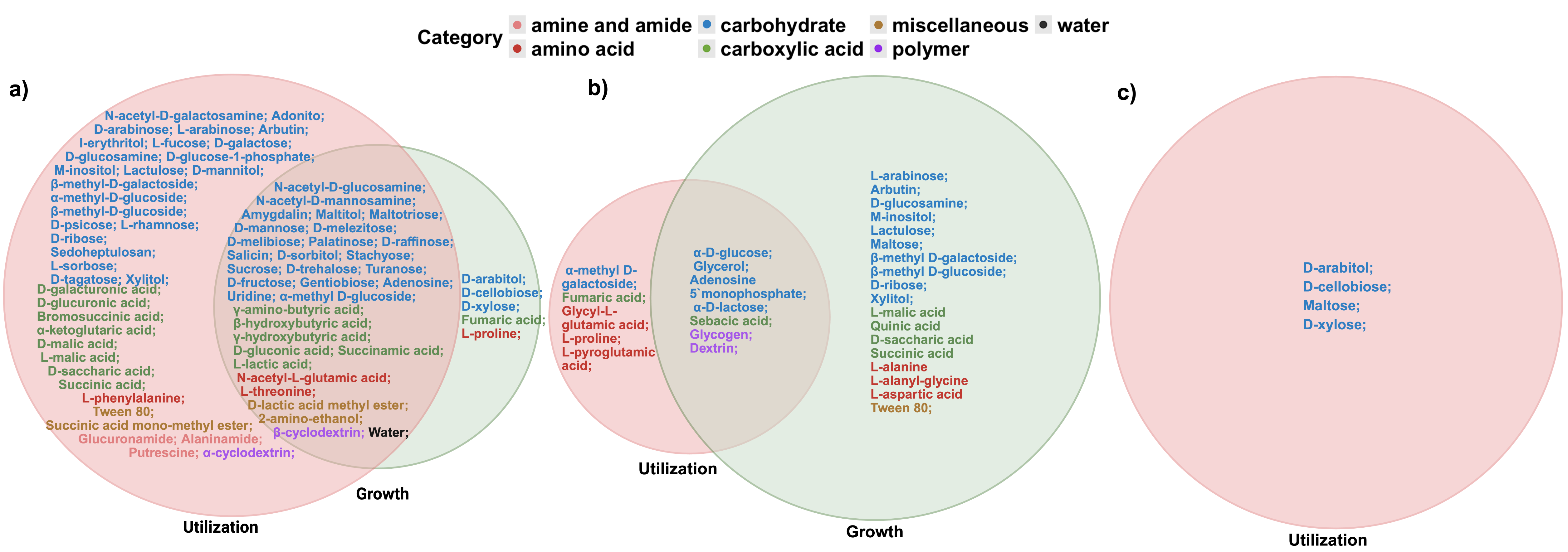

**S36 Figure. Venn diagrams showing the fastest C-substrate utilization and growth of Mr1-16 (a), Mr2575 (b), and Mr23 (c**). Fastest utilization is indicated in red, fastest growth in green, and overlaps represent substrates on which strains showed both. Substrates are grouped into amino acids, polymers, carbohydrates, carboxylic acids, amines and amides, miscellaneous, and water.

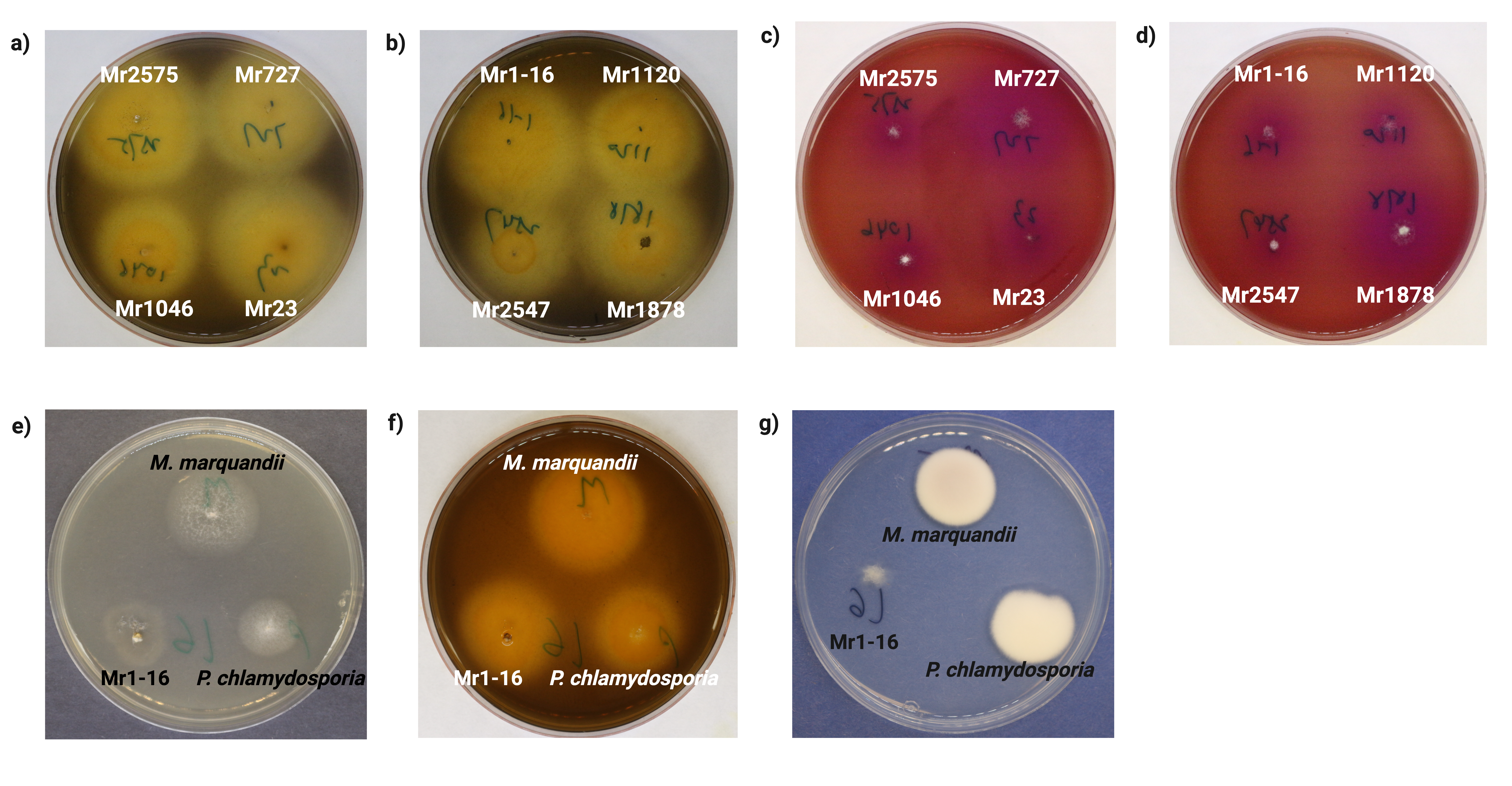

**S37 Figure. Growth of M. robertsii strains, M. marquandii, and P. chlamydosporia six days after inoculating onto minimal Noble agar with apple pectin (a–f) or D-galacturonic acid (g) as sole carbon source.** (a–b) Pectinase activity on 1% pectin visualized with iodine potassium iodide, showing transparent halos around all M. robertsii strains. (c–d) Plates with pH indicator, Bromocresol purple, revealed dark red halos around M. robertsii strains growing on 1% pectin, indicating alkalinization during pectinase activity. (e) Growth of Mr1-16, M. marquandii, and P. chlamydosporia on apple pectin medium showing that Mr1-16 produced less dense, potentially lighter colonies. (f) staining of the same plate in (e) with iodine potassium iodide to visualize pectinase activity. Halos confirmed pectin degradation by all three fungi. (f) Growth of Mr1-16, M. marquandii, and P. chlamydosporia on 1% D-galacturonic acid, the main component of pectin, showing that it supported much less growth of Mr1-16 .

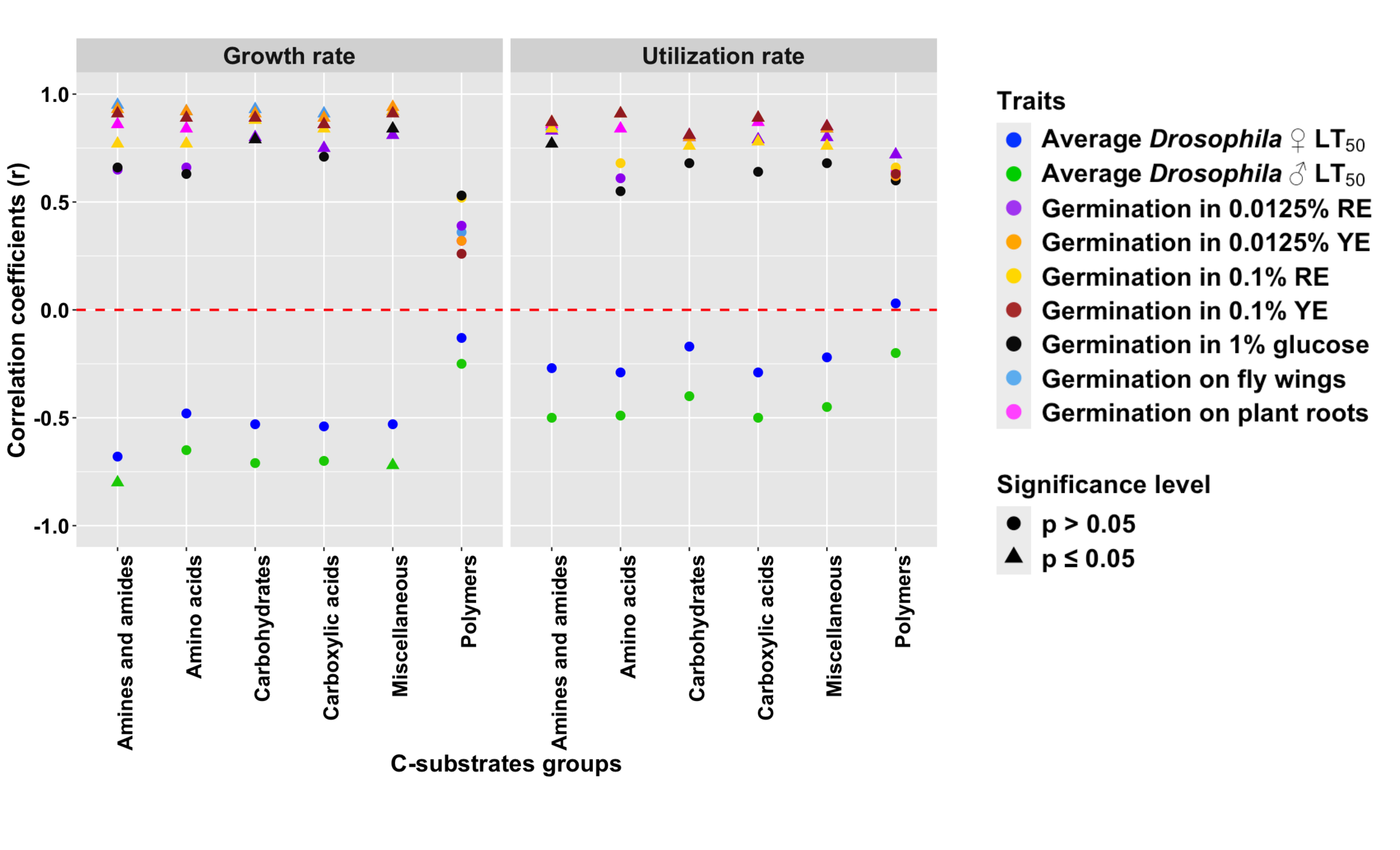
**S38 Fig. Correlation coefficients (r) depicting the relationships between the growth or utilization rate of eight *M. robertsii* strains on C-substrates and their virulence to insects, germination on plant roots, and various nutrients.** The C-substrates are categorized into six major groups: amines and amides, amino acids, carbohydrates, carboxylic acids, polymers, and miscellaneous. RE: root exudate, YE: yeast extract.
